# Purinergic signaling promotes gliomagenesis through nuclear calcium transients

**DOI:** 10.64898/2026.05.11.724407

**Authors:** Shuai Wang, Jenna Richter, Claire D. Kim, Rebecca Ronnen, Julia Cai, Yunfei Song, Sara Haddock, Mark B. Stoessel, Hannah Dada, Suzanne Gross, Adler Guerrero, Theodoris Karnavas, Gabriele Stephan, Greg Belizaire, Luke Bonanni, Danielle Golub, Jonathan Sabio, Hanna Schwarz, Albert Jiang, Nawshin Maleeha, Anuva Ghosh, Madelynn Yung, Eric Grossman-Glover, Jocelyn Tang, Laura Gherghina, Alex Donovan, Andrea Leonard, Luis Chiriboga, Drew Jones, Soomin C. Song, Esteban Mazzoni, Theresa Vincent, Robert J. Schneider, David Zagzag, Yuki Miura, Jayeeta Basu, Andrea H. Brand, Dimitris G. Placantonakis

**Affiliations:** Department of Neurosurgery, NYU Grossman School of Medicine, New York, NY, USA; Neuroscience Institute, NYU Grossman School of Medicine, New York, NY, USA; Department of Cell Biology, NYU Grossman School of Medicine, New York, NY, USA; Department of Pathology, NYU Grossman School of Medicine, New York, NY, USA; Department of Biochemistry and Molecular Pharmacology, NYU Grossman School of Medicine, New York, NY, USA; Department of Microbiology, New York University Grossman School of Medicine, New York, NY, USA; Department of Molecular Pathobiology, NYU College of Dentistry, New York, NY, USA; Department of Psychiatry, NYU Grossman School of Medicine, New York, NY, USA; Center for Neural Science, New York University, New York, NY, USA; Perlmutter Cancer Center, NYU Langone Health, New York, NY, USA

## Abstract

Intracellular Ca^2+^ transients drive key developmental and physiological processes, yet their role in oncogenesis remains incompletely understood. In glioblastoma (GBM), an aggressive brain malignancy, tumor cellular networks exhibit self-sustaining Ca^2+^ transients that promote tumor growth through unclear mechanisms. Using patient-derived GBM models, we show that these transients depend primarily on intracellular Ca^2+^ stores and extend to the nucleus to drive tumorigenesis. A neuromodulator screen identified extracellular purines ATP and ADP as potent inducers of both nuclear and cytosolic Ca^2+^ transients via activation of metabotropic purinergic P2RY1 receptors, whose knockdown attenuates tumorigenicity *in vitro* and *in vivo*. Mechanistically, Ca^2+^ transients promote tumorigenesis via the nuclear Ca^2+^/calmodulin-dependent kinase CAMK4, which regulates transcriptional and epigenetic programs, as well as ribosomal DNA transcription. From the therapeutic perspective, pharmacologic P2RY1 inhibition suppresses tumor growth *in vitro* and *in vivo*. Collectively, these findings reveal a pharmacologically targetable oncogenic mechanism in GBM and possibly other malignancies.

## Introduction

Intracellular Ca^2+^ transients are important for developmental, physiological and disease processes, but their spatiotemporal properties, mechanisms of generation, and effectors vary with cellular and tissue context and remain incompletely characterized in oncogenesis [1–3]. Glioblastoma (GBM), the most common primary brain malignancy, manifests a cellular hierarchy with GBM stem cells (GSCs) at its apex and a complex tumor microenvironment that includes immune lineages within the tumor core and neuroglial lineages in areas of brain infiltration [4, 5]. A combination of GSC-intrinsic and microenvironmental properties generates robust resistance to conventional chemoradiotherapy treatment and a poor prognosis [6, 7], facts that highlight the need for improved understanding of tumor biology and discovery of new treatment targets.

Recent studies indicated that neurons form functional synapses with GBM cells that entrain cytosolic Ca^2+^ transients and oncogenic signaling mechanisms [8–11]. This neuron-tumor interaction is particularly relevant to tumor cells infiltrating brain tissue, but does not account for Ca^2+^-dependent mechanisms within the tumor core [12], where neuronal input is essentially absent. However, previous work indicated that tumor cells can generate cell-autonomous Ca^2+^ transients in the absence of neuroglial interactions [12], raising the possibility that GBM cells in the tumor core assemble cellular networks dependent on tumor-tumor interactions that suffice to generate self-sustaining Ca^2+^ transients and activate oncogenic programs. Here, we tested this hypothesis using patient-derived GBM cells as a cellular platform, and a combination of imaging, electrophysiological, genetic, pharmacologic and sequencing techniques.

## RESULTS

### Ca^2+^ transients extend to the nucleus of GBM cells

Analysis of human GBM biospecimens indicated sparsity of neuronal axons within the tumor core, as opposed to the interface of tumor with brain tissue (**Figure S1A**). This finding justified the study of tumor cell-autonomous Ca^2+^ transients independently of neuronal input. To visualize Ca^2+^ transients in human GBM cells, patient-derived GBM cultures (PDGCs) were lentivirally engineered to express the genetically encoded Ca^2+^ sensor GCaMP6s or GCaMP6f [13] and imaged with live fluorescent microscopy. We developed a quantitative analysis pipeline using Cellpose [14], ImageJ, and Python (**Figure S1B**). Live imaging of spontaneous Ca^2+^ transients with GCaMP6s indicated that they are not confined only to the cytoplasm but also extend to the nucleus of tumor cells, and that the cytoplasmic and nuclear components are synchronous as visualized by Ca^2+^ signal tracing and heatmaps of the ΔF/F_0_ fluorescent signal (**Figure 1A**). To obtain nucleus-specific optical recordings, we lentivirally transduced PDGCs with the closely related Ca^2+^ sensor GCaMP6f engineered with a nuclear localization sequence (NLS) [15] **(Figure S1C**). We confirmed nuclear localization of the Ca^2+^ sensor using a hemagglutinin (HA)-tagged antibody to perform immunofluorescence (IF) staining **(Figure S1D)**. We compared cytoplasmic (imaged with GCaMP6s) and nuclear (imaged with GCaMP6f.3xNLS; **Figure 1B**) Ca^2+^ transients in two PDGCs and two non-neoplastic cell types: astrocytes, and human neural stem cells (NSCs), the putative cell of origin in GBM [16]. In PDGCs, cytoplasmic and nuclear Ca^2+^ transients, each lasting several seconds and similar to prior observations [12], were more pronounced and frequent than in astrocytes and NSCs (**Figure 1C-F, Figure S1E,F**), raising the possibility that they represent a preserved cellular phenomenon co-opted by tumor cells toward oncogenesis. *In vivo* two-photon microscopy of two patient-derived GBM xenografts implanted in the brain of immunodeficient NSG mice and expressing GCaMP6f.3xNLS also manifested spontaneous nuclear Ca^2+^ transients **(Figure 1G-I, Figure S1G)**, suggesting this phenomenon is real and not an artifact of in *vitro* preparations.

**Figure 1:**
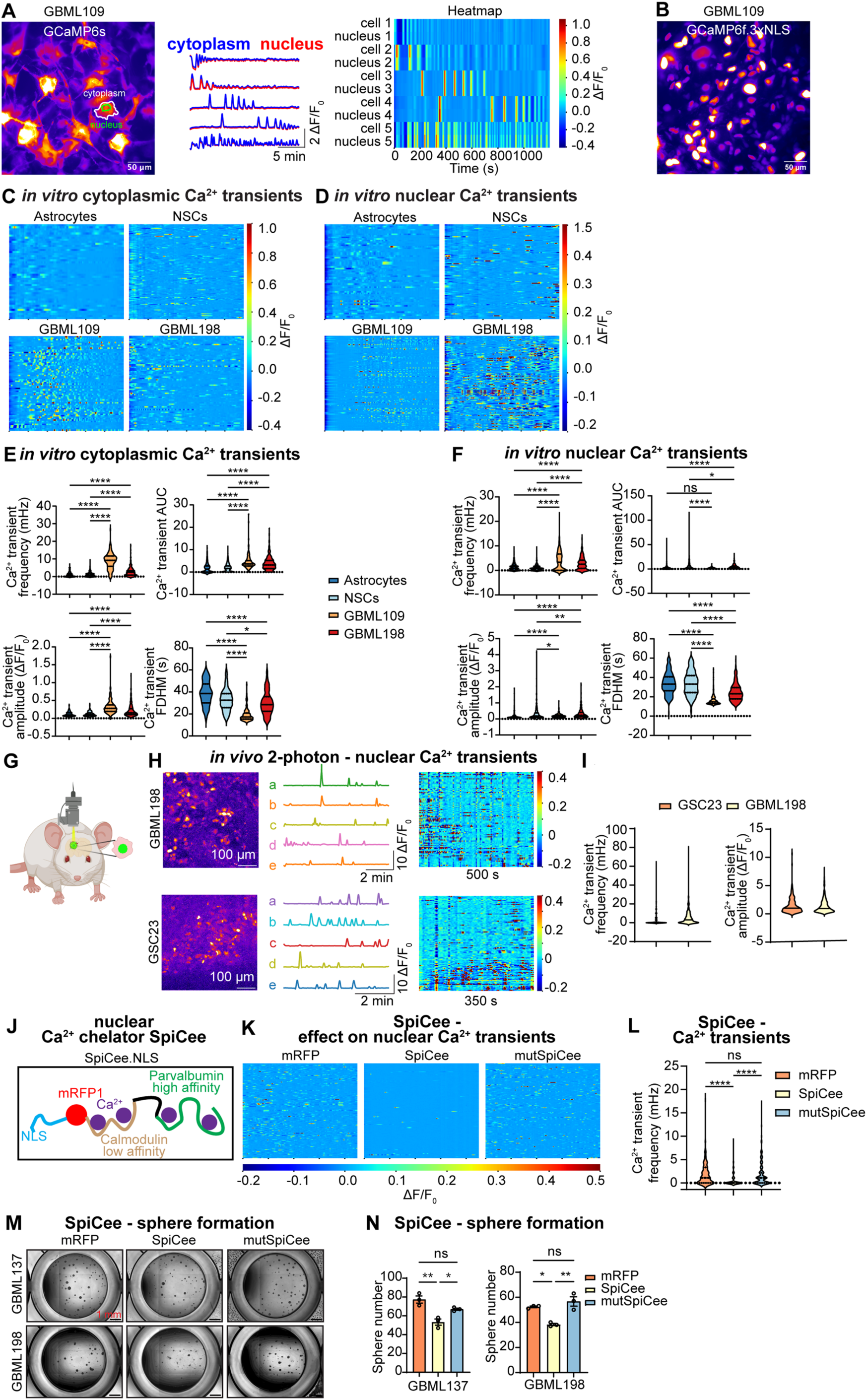
Ca^2+^ transients extend to the nucleus of GBM cells. **(A)** Cytosolic and nuclear Ca^2+^ transients in PDGCs (GBML109) transduced with GCaMP6s lentivirus (top left panel). Time-lapse images were captured with an epifluorescent microscope. Ca^2+^ transients are visualized with Ca^2+^ tracings (top right panel) and a heatmap (bottom panel) showing ΔF/F_0_. **(B)** Representative example of nuclear Ca^2+^ imaging of PDGCs with GCaMP6f.3xNLS. **(C,D)** Ca^2+^ transients in the cytoplasm **(C)** and nucleus **(D)** of immortalized human astrocytes (n_cytoplasm_ = 46, n_nucleus_ = 61), neural stem cells (NSCs) (n_cytoplasm_ = 99, n_nucleus_ = 166), GBML109 PDGCs (n_cytoplasm_ = 66, n_nucleus_ = 134), and GBML198 PDGCs (n_cytoplasm_ = 70, n_nucleus_ = 147) transduced with GCaMP6s or GCaMP6f.3xNLS lentiviruses are visualized with 20-minute heatmaps of ΔF/F_0_. **(E,F)** Violin plots show median and interquartile range of Ca^2+^ transient frequency, area under the curve (AUC), amplitude, and full duration at half-maximum (FDHM) of Ca^2+^ transients in the cytoplasm **(E)** and nucleus **(F)**. Kruskal-Wallis test (H_frequency_ = 327.7, H_AUC_ = 148.3, H_amplitude_ = 190.3, H_FDHM_ = 220.4) with *post hoc* multiple comparisons of cytoplasmic transients in **(E)**: n_astrocyte_ = 157, n_NSC_ = 257, n_GBML109_ = 151, and n_GBML198_ = 267 across three independent experiments for frequency and AUC; n_astrocyte_ = 89, n_NSC_ = 165, n_GBML109_ = 144, and n_GBML198_ = 229 across three independent experiments for amplitude and FDHM. Kruskal-Wallis test (H_frequency_ = 166.5, H_AUC_ = 81.08, H_amplitude_ = 126.3, H_FDHM_ = 435.3) with *post hoc* multiple comparisons of nuclear transients in **(F)**: n_astrocyte_ = 405, n_NSC_ = 404, n_GBML109_ = 346, and n_GBML198_ = 391 across three independent experiments for frequency and AUC; n_astrocyte_ = 263, n_NSC_ = 257, n_GBML109_ = 233, and n_GBML198_ = 296 across three independent experiments for amplitude and FDHM. **(G)** *In vivo* two-photon imaging experimental paradigm. **(H)** Representative imaging fields showing GBML198 and GSC23GBM cells transduced with GCaMP6f.3xNLS lentivirus and implanted in the brain NSG mice (left panel), representative single-cell ΔF/F_0_ traces recorded over two minutes (middle panel), and heat maps (time: GBML198 = 500 s, GSC23 = 350 s) showing nuclear Ca^2+^ transients represented as ΔF/F_0_ (right panel) (n_GSC23_ = 96, n_GBML198_ = 90). **(I)** Violin plots showing median and interquartile range of Ca^2+^ transient frequency and amplitude in the nucleus of GSC23 cells (n_frequency_ = 498, n_amplitude_ = 156) and GBML198 (n_frequency_ = 703, n_amplitude_ = 448) *in vivo* (n = 4 mice per xenograft model). **(J)** Schematic illustration of the SpiCee.NLS construct containing nuclear localization sequence (NLS), mRFP1, two calmodulin domains with low Ca^2+^-binding affinity, a linker sequence, and two high-affinity α-parvalbumin Ca^2+^-binding sites. The effector domains of calmodulin are deleted to avoid the activation of downstream effectors. **(K)** 15-minute heatmaps of ΔF/F_0_ in the nucleus of GBML198 cells transduced with mRFP (n = 169), SpiCee (n = 152), or mutSpiCee (n = 161) lentivirus. **(L)** Violin plots showing median and interquartile range of the frequency of nuclear Ca^2+^ transients in GBML198 cells transduced with mRFP (n = 281 across three recordings), SpiCee (n = 619 across three recordings), or mutSpiCee (n = 279 across three recording) lentivirus (Kruskal-Wallis test (H = 170.9) with *post hoc* multiple comparisons). **(M)** Representative images of tumor sphere formation in GBML137 and GBML198 cultures transduced with mRFP, SpiCee, or mutSpiCee lentivirus. **(N)** Bar plots showing tumor sphere numbers for GBML137 (left) and GBML198 (right) cells transduced with mRFP, SpiCee, or mutSpiCee lentivirus (one-way ANOVA, Tukey’s multiple comparisons test; GBML137, n = 3, F_(2, 6)_ = 16.58; GBML198, n = 3, F_(2, 6)_ = 17.96). * p < 0.05, ** p < 0.01, **** p < 0.0001, ns = not significant.

To determine the function of nuclear Ca^2+^ transients, we modified SpiCee, a genetically encoded Ca^2+^ chelator, fused with a NLS and mRFP1 (**Figure 1J**), and confirmed nuclear localization using confocal fluorescent microscopy (**Figure S1H**). SpiCee combines low- and high-affinity Ca^2+^ binding domains to buffer Ca^2+^ transients [17]. A mutant version, mutSpiCee, which lacks Ca^2+^ chelation ability, and mRFP1 alone served as controls. Analysis of nuclear Ca^2+^ transients indicated that SpiCee significantly reduced their frequency as compared to mRFP1 and mutSpiCee (**Figures 1K,L**). Attenuation of nuclear Ca^2+^ transients by SpiCee further inhibited tumor sphere formation (**Figures 1M,N**) and PDGC growth and viability (**Figure S1I**).

### Mechanisms generating Ca^2+^ transients

Previous studies suggested several mechanisms responsible for generation of Ca^2+^ transients in GBM, including ionotropic glutamate and metabotropic acetylcholine receptors [9, 18–20]. To further characterize the mechanisms responsible for the generation of cytoplasmic and nuclear Ca^2+^ transients in our PDGCs, we tested the contribution of plasma membrane receptors and ion channels using electrophysiological, pharmacologic and genetic approaches [12, 21].

We first measured the membrane potential of PDGCs using whole-cell patch clamp, which showed the membrane potential of PDGCs to vary from -80 mV to -20 mV (**Figure 2A, Figure S2A**). Interestingly, simultaneous patch clamp and GCaMP6s optical recordings indicated that the membrane potential did not change synchronously with Ca^2+^ transients (**Figure 2B**), raising the possibility that ion channels and ionotropic receptors may not be critical drivers of intracellular Ca^2+^ transients. Despite the high expression of a-amino-3-hydroxy-5-methyl-4-isoxazolepropionic acid (AMPA) ionotropic glutamate receptors in GBM cells (**Figure S2B**) and the reported involvement of glutamatergic neurons in the generation of Ca^2+^ transients [20], glutamate, the physiologic agonist of AMPA- and N-methyl-D-aspartate (NMDA)-type ionotropic glutamate receptors, as well as metabotropic glutamate receptors, did not enhance Ca^2+^ transients in our PDGCs (**Figure S2C,D**). Furthermore, several ionotropic receptor and voltage-gated Ca^2+^ channel (VGCC) inhibitors including NBQX (AMPA receptor antagonist), D-APV (NMDA receptor antagonist), mibefradil (T-type VGCC antagonist), ω-conotoxin (N-type VGCC), nifedipine (L-type VGCC), and ω-agatoxin (P-type VGCC) also did not show significant inhibitory effects on Ca^2+^ transients (**Figure S2E,F**). These findings suggest that ion channels and ionotropic receptors likely do not play a central role in the generation of Ca^2+^ transients in PDGCs.

**Figure 2:**
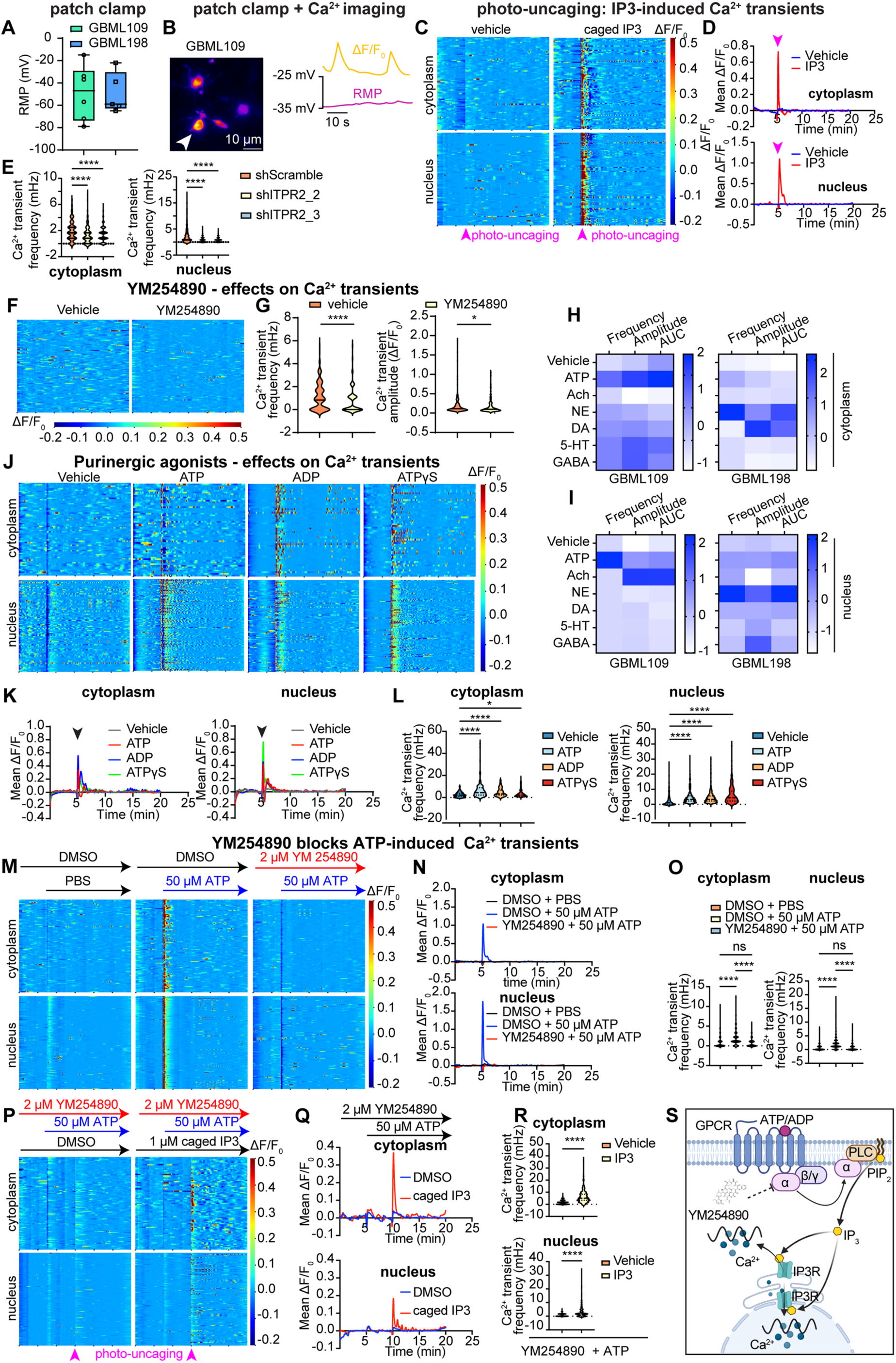
Extracellular ATP/ADP promote nuclear and cytoplasmic Ca^2+^ transients via Gα_q_ and IP_3_R-mediated Ca^2+^ release. **(A)** Box plots showing resting membrane potential (RMP) in GBML109 and GBML198 PDGCs. Box height indicates the interquartile range, the black horizontal line indicates median, and whiskers represent minimal and maximal values (n_GBML109_ = 6, n_GBML198_ = 5). **(B)** Representative example of simultaneous whole-cell patch-clamp recording of RMP and live Ca^2+^ imaging in GBML109 cells transduced with GCaMP6s lentivirus. **(C)** Twenty-minute heatmaps and **(D)** averaged tracing of ΔF/F_0_ in the cytoplasm and nucleus of GBML198 cells treated with caged IP_3_ (ci-IP_3_) or vehicle control (cytoplasm: n_vehicle_ = 78, n_IP3_ = 84; nucleus: n_vehicle_ = 88, n_IP3_ = 106). Pink arrows indicate photo-uncaging at the 5-minute mark. **(E)** Violin plots showing median and interquartile range of the frequency of Ca^2+^ transients in the cytoplasm and nucleus of GBML198 cells transduced with control shScramble and two non-overlapping ITPR2-targeting shRNAs, shITPR2_2 and shITPR2_3. Kruskal-Wallis test (H_cytoplasm_ = 38.49, H_nucleus_ = 125.2) with multiple comparisons of cytoplasmic (n_shScramble_ = 247, n_shITPR2_2_ = 200 and n_shITPR2_3_ = 289 across three recordings) and nuclear (n_shScramble_ = 405, n_shITPR2_2_ = 267, and n_shITPR2_3_ = 424 across three recordings) tracings. **(F,G)** Representative 20-minute heatmaps of ΔF/F_0_ (n_vehicle_ = 114, n_YM254890_ = 107) **(F)** and violin plots showing median and interquartile range of the frequency (n_vehicle_ = 313, n_YM254890_ = 264) and amplitude (n_vehicle_ = 196, n_YM254890_ = 112) of cytoplasmic Ca^2+^ transients **(G)** in GBML198 cells treated with 2 μM G_q_ inhibitor YM254890 or vehicle control (Mann-Whitney U test). **(H,I)** Heatmaps of neuromodulator screening results showing z-scores of Ca^2+^ transient frequency, AUC, and amplitude in the cytoplasm **(H)** and nucleus **(I)** of two PDGCs. Neuromodulator screen concentrations: 100 μM ATP, 1 mM acetylcholine (Ach), 100 μM norepinephrine (NE), 100 μM dopamine (DA), 100 μM serotonin (5-HT), and 100 μM gamma-aminobutyric acid (GABA). **(J,K)** Twenty-minute heatmaps of ΔF/F_0_ (**J**) and averaged Δ*F*/*F*_0_ traces (**K**) in the cytoplasm (n_vehicle_ = 69, n_ATP_ = 57, n_ADP_ = 53, n_ATPγS_ = 54) and nucleus (n_vehicle_ = 90, n_ATP_ = 113, n_ADP_ = 103, n_ATPγS_ = 122) of GBML198 cells in response to vehicle,100 μM ATP, 100 μM ADP, and 100 μM ATPγS. **(L)** Violin plots showing median and interquartile range of the frequency of Ca^2+^ transients in the cytoplasm and nucleus of GBML198 treated with vehicle, ATP, ADP, or ATPγS. Kruskal-Wallis test with multiple comparisons of cytoplasmic transients: n_vehicle_ = 177, n_ATP_ = 202, n_ADP_ = 168, and n_ATPγS_ = 248 across 3 recordings and nuclear transients: n_vehicle_ = 282, n_ATP_ = 509, n_ADP_ = 290, and n_ATPγS_ = 343 across 3 recordings. **(M-O)** Representative 20-minute heatmaps (cytoplasm: n_DMSO + PBS_ = 109, n_DMSO + ATP_ = 107, and n_YM254890 + ATP_ = 113; nucleus; n_DMSO + PBS_ = 166, n_DMSO + ATP_ = 227, and n_YM254890 + ATP_ = 165) of ΔF/F_0_ (**M**), averaged ΔF/F_0_ tracings (**N**), and violin plots indicating median and interquartile range **(O)** of cytoplasmic and nuclear Ca^2+^ transients in GBML198 cells pre-treated with 2 μM G_q_ inhibitor YM254890 or vehicle control (DMSO) after addition of 50 μM ATP or vehicle control (PBS) (cytoplasm: n_DMSO + PBS_ = 353, n_DMSO + ATP_ = 306, and n_YM254890 + ATP_ = 292; nucleus; n_DMSO + PBS_ = 400, n_DMSO + ATP_ = 490, and n_YM254890 + ATP_ = 412 across three separate recordings). **(P-R)** Cytoplasmic and nuclear Ca^2+^ transients in GBML198 treated with 2 μM G_q_ inhibitor YM254890, 50 μM ATP and 1 μM ci-IP_3_ or vehicle control (DMSO), represented as 20-minute heatmaps (cytoplasm: n_vehicle_ = 95, n_IP3_ = 77; nucleus: n_vehicle_ = 141, n_IP3_ = 158) of ΔF/F_0_ **(P)**, averaged ΔF/F_0_ tracings **(Q)**, and violin plots **(R)** indicating median and interquartile range of Ca^2+^ transient frequency (Mann-Whitney U test; cytoplasm: n_vehicle_ = 324, n_IP3_ = 363; nucleus: n_vehicle_ = 397, n_IP3_ = 432)**. (S)** Schematic showing the generation of cytoplasmic and nuclear Ca^2+^ transients in PDGCs by eATP/eADP through activation of G_q_-coupled purinergic receptors. * p < 0.05, ** p < 0.01, *** p < 0.001, **** p < 0.0001, ns = not significant.

In non-excitable cells, Ca^2+^ transients are thought to arise via activation of inositol 1,4,5-trisphosphate receptors (IP_3_R) and Ca^2+^ flux from the endoplasmic reticulum (ER) to the cytosol mutSpiCee [22–24]. We tested whether this paradigm holds true in our PDGCs. The activation of IP_3_ receptors by photo-uncaging ci-IP_3_ with UV light induced robust Ca^2+^ transients in both the cytoplasm and nucleus of PDGCs (**Figure 2C,D, Figure S2G,H**). Single cell RNA-Seq (scRNA-seq) analysis indicated that *ITPR2* is the most highly expressed of the three IP_3_R isoforms in GBM (**Figure S2I**). Immunofluorescent staining demonstrated ITPR2 expression in 2 PDGCs (**Figure S2J**). To evaluate the function of ITPR2 in PDGCs, we used two distinct non-overlapping short hairpin RNAs (shRNAs) to knockdown ITPR2 expression (**Figure S2K**). The frequency of Ca^2+^ transients in both the cytoplasm and nucleus decreased significantly after ITPR2 knockdown (**Figure 2E, Figure S2L**). ITPR2 knockdown also attenuated Ca^2+^ transients induced by photo-uncaged ci-IP_3_ (**Figure S2M,N**). Given that G protein-coupled receptors (GPCRs) coupled to Gα_q_ are a dominant source of IP_3_R-mediated Ca^2+^ release [25], we tested whether G_q_ inhibition influences Ca^2+^ transients in PDGCs. Indeed, the G_q_ inhibitor YM254890 [26] significantly reduced the frequency and amplitude of both cytosolic and nuclear Ca^2+^ transients (**Figure 2F,G, Figure S2O,P**). These data suggest that G_q_-coupled GPCRs, IP_3_ and ITPR2 are crucial to the generation of Ca^2+^ transients in GBM cells.

Recent studies have suggested that neurotransmitters/neuromodulators regulate neural circuits in part through GPCR-induced spontaneous Ca^2+^ events in astrocytes [27, 28]. Based on this information and the fact the GBM cells express receptors to major neurotransmitters and neuromodulators [20], we screened several neurotransmitters to evaluate their effects on cytosolic and nuclear Ca^2+^ transients in two different PDGCs. These included extracellular adenosine triphosphate (eATP), acetylcholine (Ach), norepinephrine (NE), dopamine (DA), serotonin (5-HT), and γ-aminobutyric acid (GABA), all of which have been linked directly or indirectly to regulation of intracellular Ca^2+^ [27–32]. eATP, Ach, and NE increased the frequency, area under the curve (AUC), and amplitude of cytoplasmic Ca^2+^ transients, but only eATP reproducibly enhanced nuclear Ca^2+^ transient frequency, AUC, and amplitude (**Figure 2H,I, Figure S2Q,R, Figure S3A,B**). Both eATP and its metabolite adenosine diphosphate (eADP), obtained from eATP through the enzymatic action of ectonucleotidases, are known to act as direct agonists of purinergic receptors [33]. We therefore tested three different purinergic receptor agonists for their effects on Ca^2+^ transients: eATP, eADP, and the non-hydrolyzable ATP analog eATPγS. All three agonists increased cytoplasmic and nuclear Ca^2+^ transient frequency, amplitude, and AUC (**Figure 2J-L, Figure S3C,D**). We also generated dose-response curves, which indicated that concentrations of eADP and eATP as low as 1-10 μM produce Ca^2+^ transients in both the nucleus and cytoplasm (**Figure S3E-G**).

Knockdown of ITPR2 significantly reduced ATP-driven Ca^2+^ transients in both the cytoplasm and nucleus (**Figure S3H,I**), suggesting eATP operates through the Gα_q_-IP_3_R pathway to mobilize Ca^2+^. To further evaluate this hypothesis, we tested the effects of G_q_ inhibitor YM254890 on eATP-induced cytosolic and nuclear Ca^2+^ transients. Pre-treatment with YM254890 blocked eATP-induced cytosolic and nuclear Ca^2+^ transients in our PDGCs (**Figure 2M-O, Figure S3J,K**). Photo-uncaging ci-IP_3_ still generated Ca^2+^ transients in the presence of YM254890 (**Figure 2P-R, Figure S3L,M**), confirming IP_3_R operates downstream of G_q_ in this pathway. Collectively, these data suggest eATP and eADP activate G_q_-coupled purinergic receptors to generate cytoplasmic and nuclear Ca^2+^ transients in PDGCs (**Figure 2S**).

To confirm that eATP/ADP contribute to the spontaneous cytoplasmic and nuclear Ca^2+^ transients, we treated our PDGCs with apyrase, an ATP- and ADP-hydrolyzing enzyme [34]. Apyrase reduced eATP in a dose-dependent manner in luminescence-based eATP detection assays (**Figure S4A**) and in live fluorescence imaging assays with the fluorescent eATP sensor GRAB [35] (**Figure 3A, Figure S4B**). Importantly, apyrase significantly reduced spontaneous cytoplasmic and nuclear Ca^2+^ transients (**Figure 3B, Figure S4C,D**), suggesting PDGCs release eATP which helps drive spontaneous Ca^2+^ transients.

**Figure 3:**
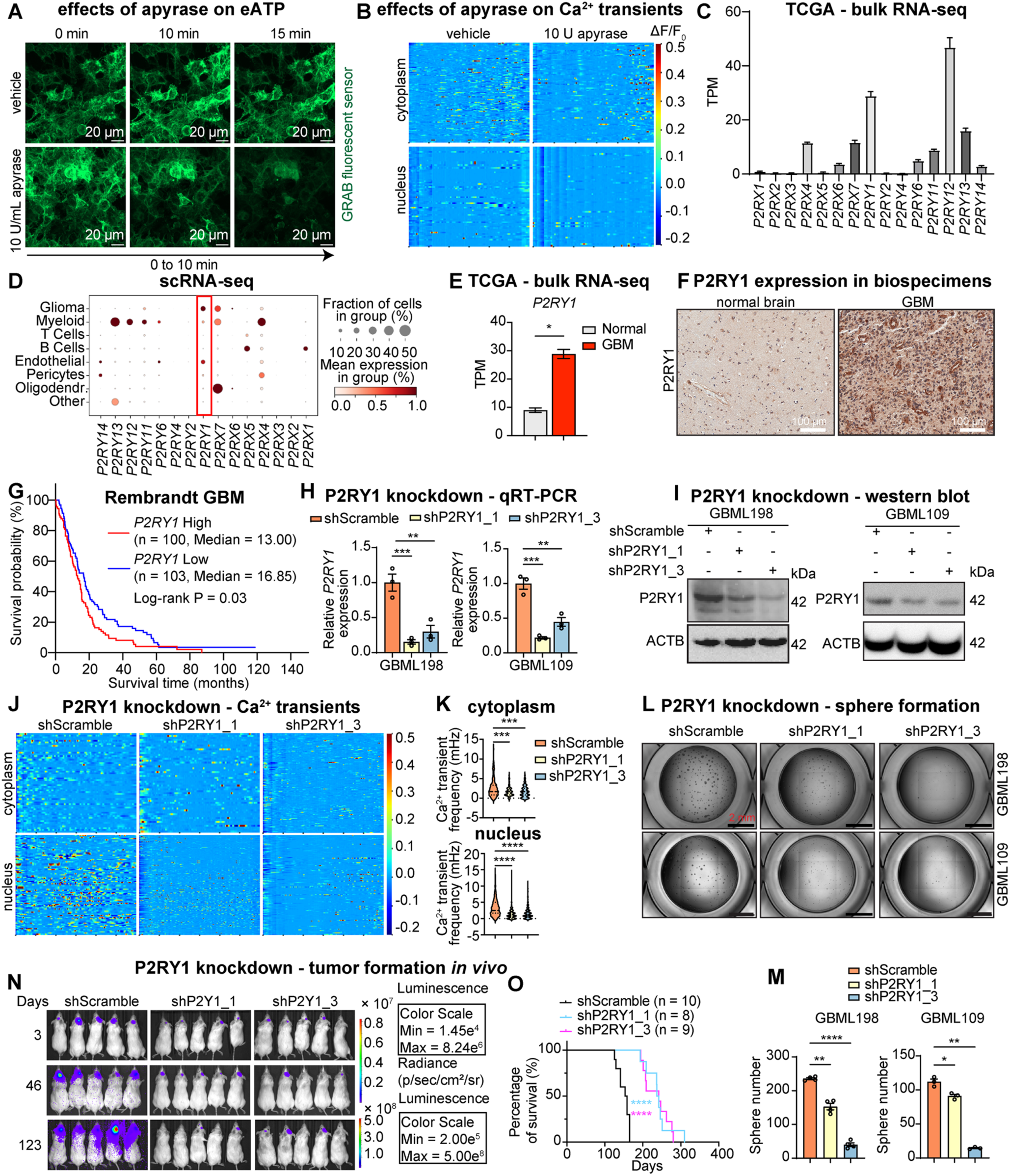
Purinergic receptor P2RY1 promotes Ca^2+^ transients and GBM growth. **(A)** Representative fluorescence images of GBML198 cells transduced with the fluorescent eATP sensor GRAB and treated with either vehicle or 10 U/mL apyrase, an ATP- and ADP-hydrolyzing enzyme. **(B)** Representative 20-minute heatmaps (cytoplasm: n_vehicle_ = 111, n_apyrase_ = 80; nucleus: n_vehicle_ = 68, n_apyrase_ = 62) of ΔF/F_0_ of Ca^2+^ transients in the cytoplasm and nucleus of GBML198 cells treated with vehicle or 10 U/mL apyrase. **(C)** Bar plot showing the expression level (Transcripts per Million, TPM) of purinergic receptors in IDH wildtype GBM specimens (n = 143) from TCGA database. **(D)** Dot plot showing expression of purinergic receptors in single-cell RNA-seq data from operative GBM specimens (GSE182109). **(E)** Bar plot showing the expression level (TPM) of *P2RY1* in normal brain tissue (n = 5) and IDH wildtype GBM specimens (n = 143) from TCGA database (unpaired *t*-test). **(F)** Immunohistochemistry staining of P2RY1 in normal brain and a GBM specimen. **(G)** Kaplan-Meier plot showing overall survival of GBM patients stratified by high (n = 100) vs. low *P2RY1* expression (n = 103) from the Rembrandt database. **(H)** Relative mRNA level of *P2RY1* in GBML198 and GBML109 transduced with shScramble or one of two non-overlapping shRNA lentiviruses targeting P2RY1 (n = 3/group, one-way ANOVA with Dunnett’s multiple comparisons test; GBML198: F_(2, 6)_ = 26.25; GBML109: F_(2, 6)_ = 44.89). **(I)** Immunoblot for P2RY1 and ACTB in GBML198 and GBML109 transduced with shScramble and one of two non-overlapping shRNA lentiviruses targeting P2RY1. **(J)** Heatmaps showing 20-minute recordings of ΔF/F_0_ in the cytoplasm and nucleus of GBML198 cells transduced with shScramble or one of two non-overlapping shRNA targeting P2RY1 (cytoplasm: n_shScramble_ = 58, n_shP2RY1_1_ = 52, n_shP2RY1_3_ = 124; nucleus: n_shScramble_ = 91, n_shP2RY1_1_ = 213, n_shP2RY1_3_ = 135). **(K)** Violin plots showing median and interquartile range of Ca^2+^ transient frequency in the cytoplasm (top panel) and nucleus (bottom panel) across 3 recordings of GBML198 transduced with shScramble, shP2RY1_1, and shP2RY1*_3* lentiviruses (Kruskal-Wallis test (H = 18.49) with *post hoc* multiple comparisons of transients; cytoplasm: n_shScramble_ = 102, n_shP2RY1_1_ = 230, n_shP2RY1_3_ = 289; nucleus: n_shScramble_ = 578, n_shP2RY1_1_ = 414, n_shP2RY1_3_ = 427). **(L)** Representative images of tumor sphere formation assays in GBML198 and GBML109 cultures transduced with shScramble or one of two non-overlapping shRNA targeting P2RY1. **(M)** Bar graphs of tumor sphere formation assays in GBML198 (left) and GBML109 (right) cultures transduced with shScramble or one of two non-overlapping shRNA lentiviruses targeting P2RY1 (one-way ANOVA with Dunnett’s multiple comparisons test; GBML198, n = 4/group, F_(2, 8)_ = 441.1; GBML109, n = 3/group, F_(2, 6)_ = 328.8). **(N)** Longitudinal bioluminescent images of NSG mice injected with GBML198 cells transduced with shScramble, shP2RY1_2, or shP2RY1_3 lentiviruses. **(O)** Kaplan-Meier survival curve showing survival time of mice xenografted with GBML198 cells transduced with shScramble, shP2RY1_2*, or* shP2RY1_3 lentiviruses (n = 10, 8, and 9 mice/group; log-rank test). * p < 0.05, ** p < 0.01, *** p < 0.001, **** p < 0.0001.

### P2RY1 knockdown attenuates Ca^2+^ transients and tumor growth

Given our findings that eATP/ADP promote Ca^2+^ transients in an G_q_/ITPR2-dependent manner, we next sought to determine the relevant upstream G_q_-coupled purinergic receptor. Bulk RNAseq analysis of IDH (Isocitrate DeHydrogenase) wildtype GBM in the TCGA suggested that *P2RY1, P2RY12* and *P2RY13* are the most highly expressed metabotropic purinergic receptors (**Figure 3C**). Single-cell RNA-seq analysis demonstrated that *P2RY1* is highly expressed in tumor cells, whereas *P2RY12* and *P2RY13* are expressed in immune cells of the tumor microenvironment (**Figure 3D**). *P2RY1* was also found to be highly expressed in GBM compared to normal brain in TCGA (**Figure 3E**). Immunohistochemistry (IHC) staining of our operative GBM specimens showed that P2RY1 is highly expressed in GBM tumors but not normal brain (**Figure 3F, Figure S4E**). Immunofluorescence (IF) staining demonstrated P2RY1 expression in PDGCs *in vitro* (**Figure S4F**). Differentiation of GSCs with serum showed persistent and even increased expression of *P2RY1* (**Figure S4G**), suggesting its expression is maintained whether tumor cells are stem-like or differentiated. The differentiation of GSCs was validated by the significant decrease in *OLIG2* transcript [36] in DGCs (**Figure S4G**). Survival analysis using the Rembrandt and CGGA databases indicated that high expression of *P2RY1* predicted poorer prognosis in GBM patients (**Figure 3G, Figure S4H**).

We knocked down P2RY1, whose ligands include both eATP and ADP [37], with two non-overlapping shRNAs to study its function in PDGCs (**Figures 3H,I**). Knockdown of P2YR1 attenuated both spontaneous (**Figures 3J,K**) and eATP-induced (**Figures S4I,J**) nuclear and cytosolic Ca^2+^ transients. P2RY1 knockdown also inhibited tumor sphere formation *in vitro* (**Figure 3 L,M**), tumor xenograft growth *in vivo* (**Figure 3N**, **Figure S4K**), and prolonged survival (**Figure 3O**). These findings suggest that eATP/ADP activate P2RY1 to generate Ca^2+^ transients in the cytoplasm and nucleus and promote tumor growth.

### Ca^2+^/calmodulin signaling is an important effector of nuclear Ca^2+^ transients

Calmodulin, a principal intracellular sensor of elevations in Ca^2+^ levels, complexes with Ca^2+^ to activate Ca^2+^/calmodulin-dependent kinases [38]. Treatment of two PDGCs with KN93, an inhibitor of Ca^2+^/calmodulin-dependent kinase activity, reduced their growth and viability, whereas its inactive analog KN92 [39] had no significant effect (**Figure 4A**). To test whether Ca^2+^/calmodulin signaling is activated by nuclear Ca^2+^ transients, we lentivirally transduced PDGCs with CaMBP4, which consists of four copies of the M13 calmodulin-binding peptide derived from myosin light chain kinase, and prevents nuclear Ca^2+^/calmodulin signaling [40]. Immunofluorescence microscopy validated the localization of CaMBP4 in the nucleus, predominantly in the nucleolar regions (**Figure 4B**). CaMBP4 inhibited stem-like properties of PDGCs, as measured by extreme limiting dilution (**Figure 4C**) and tumor sphere formation assays (**Figures 4D,E**), suggesting the potential tumorigenic role of Ca^2+^/calmodulin signaling. Furthermore, CaMBP4 co-localized with fibrillarin, a nucleolar region marker (**Figure 4F**), and reduced the number of nucleolar organizer regions per cell (**Figure 4G**). Consistent with its localization in nucleolar regions, CaMBP4 also decreased ribosomal RNA (47S and 5.8S) levels, as quantified by qRT-PCR (**Figure 4H**). Importantly, perturbing Ca^2+^/calmodulin signaling either with CAMBP4 or chelation of nuclear Ca^2+^ with SpiCee.NLS inhibited the invasive properties of PDGCs (**Figure 4I,J**). In summary, these findings suggest that nuclear Ca^2+^/calmodulin signaling is important for GBM growth and invasion

**Figure 4:**
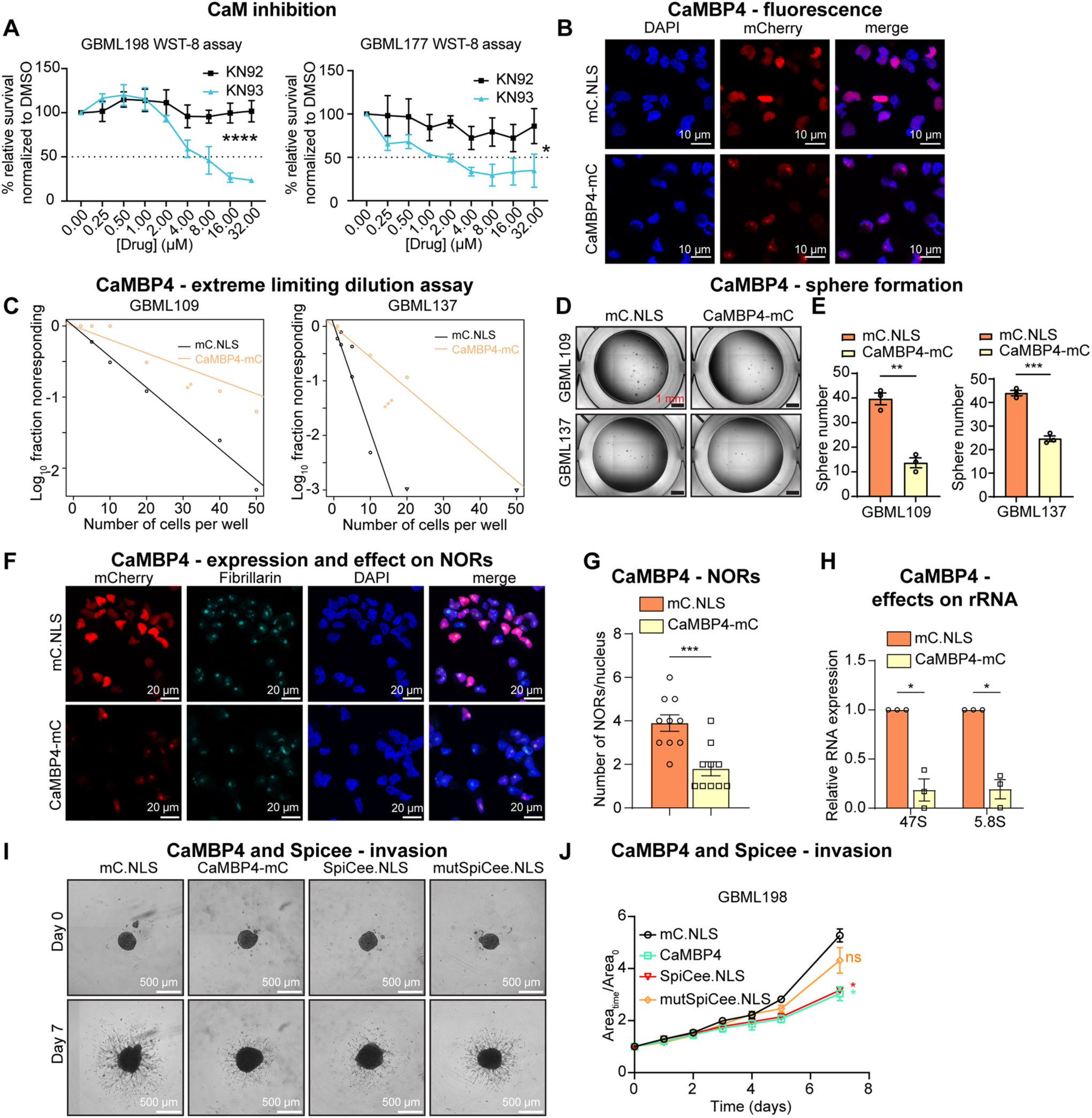
Oncogenic actions of nuclear Ca^2+^/calmodulin signaling. **(A)** WST-8 assays in GBML198 (left) and GBML177 (right) cells treated with KN92 or KN93 (n = 3/concentration; two-way ANOVA; GBML198: F_(1,36)_ = 30.90, p < 0.0001; GBML177: F_(1,34)_ = 28.30, p < 0.05). **(B)** Representative fluorescent images showing signals of mCherry, and DAPI in GBML137 cells transduced with mC.NLS, or CaMBP4-mC lentiviruses. **(C)** Plot showing initially seeded number of cells/well against the Log_10_ fraction of non-responder wells without any detected spheres in GBML109 and GBML137 cells transduced with mC.NLS or CaMBP4-mC (ξ^2^ = 9.8, ξ^2^ = 16.5). **(D)** Representative images of tumor sphere formation assays in GBML109 and GBML137 transduced with mC.NLS or CaMBP4-mC lentivirus. **(E)** Bar graphs of tumor sphere formation assays in GBML109 (left) and GBML137 (right) cells transduced with mC.NLS, or CaMBP4-mC lentiviruses (n = 3/group, unpaired t-test). **(F)** Representative IF images showing mCherry, fibrillarin, and DAPI in GBML137 cells transduced with mc.NLS or CaMBP4-mC lentiviruses. **(G)** Bar plot showing the number of nucleolar organizer regions (NORs) per nucleus, as labeled with fibrillarin, in GBML137 cells transduced with mC.NLS, or CaMBP4-mC lentiviruses (unpaired *t*-test, n = 10/group). **(H)** Bar graph depicting 47S and 5.8S rRNA expression relative to *ACTB*, as measured by qRT-PCR in GBML137 cells transduced with mC.NLS or CaMBP4-mC lentiviruses (n = 3/group, paired t-test). **(I)** Representative images of 3-dimensional (3D) invasion assays at days 0 and 7 in GBML198 cells transduced with mc.NLS, CaMBP4-mC, SpiCee.NLS, and mutSpiCee.NLS lentiviruses. **(J)** Line graph showing area_time_/area_0_ (y-axis) over time (x-axis) of 3D invasion assays in GBML198 cells transduced with mc.NLS, CaMBP4-mC, SpiCee.NLS, and mutSpiCee.NLS lentiviruses (two-way repeated measures (RM) ANOVA, n = 3/group, F_(24, 60)_ = 8.515). * p < 0.05, ** p < 0.01, *** p < 0.001, ns = not significant.

### Knockdown of CAMK4 inhibits tumor growth

CAMK4 is the only Ca^2+^/calmodulin dependent kinase that localizes to the nucleus [41] and is known to mediate lymphocyte maturation [42] and neuronal plasticity [43]. We next tested its involvement in GBM tumorigenesis. Immunohistochemistry staining of our GBM operative specimens and normal brain showed CAMK4 expression in tumors and neocortex (**Figure 5A, Figure S5A**). Single-cell RNA-seq analysis of GBM tumors showed that the expression of *CAMK4* is highest in tumor cells and tumor-associated T cells (**Figure 5B**). Immunofluorescence staining of CAMK4 in PDGCs demonstrated that CAMK4 mainly localizes to the nucleus of GBM cells, including nucleolar regions (**Figure 5C, Figure S5B**). Importantly, CAMK4 protein levels were found to be higher in PDGCs compared to NSCs, the presumed cell of origin in GBM (**Figures 5D,E**), suggesting a role in gliomagenesis. *CAMK4* expression trended to lower levels after serum-induced differentiation of GSCs into DGCs (**Figure S5C**), suggesting a role in GSC maintenance.

**Figure 5:**
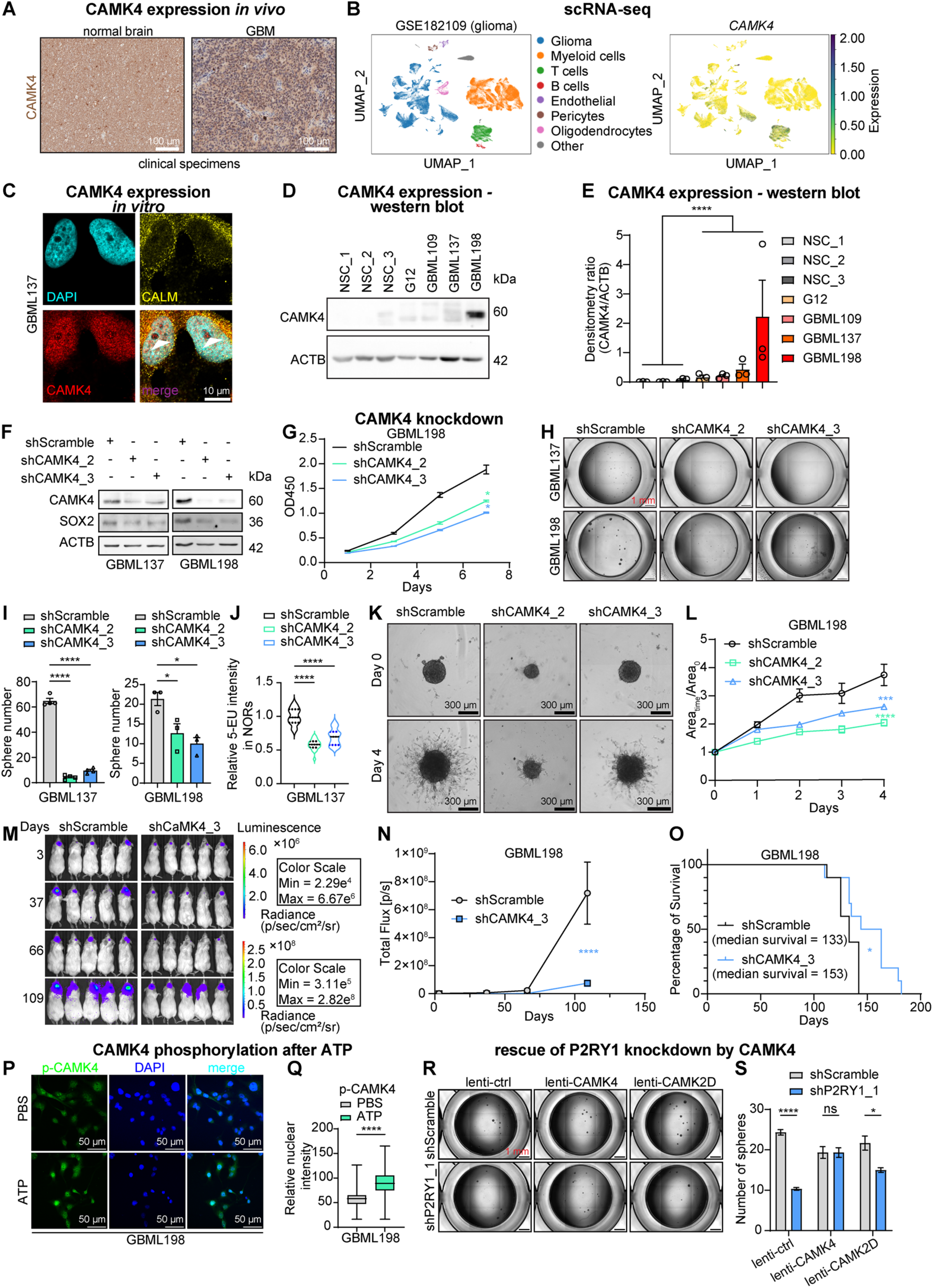
CAMK4 is essential for GBM growth. **(A)** Immunohistochemistry staining of CAMK4 in normal brain and an operative GBM specimen. **(B)** UMAP showing the expression of CAMK4 in different cell types in a GBM scRNA-seq dataset (GSE182109). **(C)** Representative fluorescent images of DAPI, CALM (calmodulin), and CAMK4 in GBML137 cells. White arrowheads in the merged panel indicate CAMK4 localization in nucleolar regions. **(D)** Representative immunoblot for CAMK4 and ACTB in three human embryonic stem cell (hESC)-derived NSC preparations and 4 PDGCs. **(E)** Bar graph depicting densitometry ratio of CAMK4 relative to ACTB in immunoblots from NSCs and PDGCs (unpaired t-test, n = 3 for each NSC or PDGC). **(F)** Immunoblot for CAMK4, SOX2, and ACTB from GBML137 and GBML198 cells transduced with shScramble and two non-overlapping shRNA lentiviruses targeting CAMK4. **(G)** Line graph depicting absorbance (OD450) over time in WST-8 assays in GBML198 cells transduced with shScramble or one of two non-overlapping shRNA targeting CAMK4 (two-way ANOVA with Dunnett’s multiple comparisons test, n = 3/group, F_(6, 18)_ = 41.22). **(H)** Representative images of tumor sphere formation assays in GBML137 and GBML198 cells transduced with shScramble or one of two non-overlapping shRNA lentiviruses targeting CAMK4. **(I)** Bar graphs of sphere formation assays in GBML137 (left) and GBML198 (right) transduced with shScramble or one of two non-overlapping shRNA targeting CAMK4 (one-way ANOVA with Tukey’s multiple comparisons test; GBML137, n = 4/group, F_(2, 9)_ = 542.8; GBML198, n = 3/group, F_(2, 6)_ = 9.673). **(J)** Violin plot showing relative 5-EU intensity in nucleolar organizer regions (NORs) of GBML137 cells transduced with shScramble or one of two non-overlapping shRNA targeting CAMK4. Horizontal dash lines indicate the interquartile range, and medians are indicated by a black horizontal line (shScramble, n = 29; shCAMK4_2, n = 23; shCAMK4_3, n = 51; Kruskal Wallis H = 62.95 with *post hoc* multiple comparisons). **(K)** Representative images of 3D invasion assays at days 0 and 4 in GBML198 cells transduced with shScramble or one of two non-overlapping shRNA targeting CAMK4. **(L)** Line graph showing area_time_ /area_0_ over time of 3D invasion assays in GBML198 cells transduced with shScramble or shRNAs targeting CAMK4 (n = 4/group, two-way ANOVA with Šídák’s multiple comparisons test, F_(8, 45)_ = 4.717). **(M)** Longitudinal bioluminescent images of NSG mice injected with GBML198 cells transduced with shScramble or shCAMK4_3 lentiviruses. **(N)** Line graph depicting total bioluminescence flux as a function of time from the *in vivo* bioluminescent imaging experiment in (**M**) (n = 10/group, two-way ANOVA with Šídák’s multiple comparisons test, F_(3, 54)_ = 8.245). **(O)** Kaplan-Meier curve shows improved survival of mice xenografted with GBML198 cells transduced with shCaMK4_3 vs. shScramble (n = 10 mice/group; log-rank test). **(P)** Representative IF images of phospho-CAMK4 (p-CAMK4) staining in GBML198 cells treated with PBS or 100 μM ATP for 5 minutes. **(Q)** Box plot showing the fluorescence intensity of p-CAMK4 in GBML198 cells treated with PBS (n = 56) or 100 μM ATP (n = 55) for 5 minutes (Mann-Whitney U test). **(R)** Representative images of tumor sphere formation assays in shScramble or shP2Y1_1 G12 cells transduced with either lenti-ctrl, lenti-CaMK4, or lenti-CaMK2D lentiviruses. **(S)** Bar plot showing tumor sphere numbers of G12 cultures in **(R)** (multiple t-tests). * p < 0.05, *** p < 0.001, **** p < 0.0001, ns = not significant.

Overexpression of CAMK4 in PDGCs (**Figure S5D**) resulted in reduced sensitivity to chemical Ca^2+^ chelator BAPTA-AM (**Figure S5E**), suggesting that CAMK4 may represent an important effector of Ca^2+^ transients. Knockdown of CAMK4 using two non-overlapping shRNA constructs in PDGCs (**Figure 5F**) impeded cellular growth/viability (**Figure 5G**) and tumor sphere formation (**Figures 5H,I, Figure S5F**). Consistent with the effects of CaMBP4, knockdown of CAMK4 also suppressed ribosome biogenesis, as measured by 5-EU incorporation assays [44] (**Figure 5J, Figure S5G**), and nucleolar size (**Figure S5H**). CAMK4 knockdown also reduced invasive properties of PDGCs (**Figure 5K,L, Figure S5I,J**). *In vivo* experiments demonstrated that CAMK4 knockdown impeded xenograft tumor growth (**Figure 5M,N, Figure S5K,L**) and prolonged the survival of tumor-bearing mice (**Figure 5O, Figure S5M**).

We then tested whether CAMK4 functions as a downstream effector of eATP/ADP and P2RY1 signaling in PDGCs. Immunofluorescence microscopy demonstrated that activation of purinergic signaling with eATP significantly increases phosphorylation of CAMK4 (Thr200) in the nucleus (**Figure 5P,Q**), a modification associated with increased CAMK4 enzymatic activity. Given that Ca^2+^ transients in the cytoplasm likely also have tumorigenic functions, we explored the function of cytosolic Ca^2+^/calmodulin-dependent kinases, such as CAMK2. Single-cell RNA-seq analysis indicated that *CAMK2D* expression is the highest among the CAMK2 isoforms in GBM (**Figure S5N**). Knockdown of CAMK2D (**Figure S5O**) reduced the tumor sphere formation ability of PDGCs (**Figure S5P**). However, only the overexpression of CAMK4, but not CAMK2D (**Figure S5Q**), was able to rescue the growth impediment associated with P2RY1 knockdown (**Figure 5R,S**). These observations suggest that CAMK4 is essential for PDGC growth *in vitro* and *in vivo* and that it functions downstream of P2RY1.

### Oncogenic mechanisms of CAMK4

To investigate how CAMK4 regulates GBM growth, we performed RNA-Seq analysis using two non-overlapping shRNAs targeting CAMK4. Principal component analysis (PCA) of raw count data revealed sample separation based on CAMK4 knockdown (shCAMK4) vs. shScramble control (**Figure S6A**). Normalized counts of *CAMK4* transcript confirmed on-target activity of both shRNAs (**Figure S6B**). Volcano plots additionally showed the shRNAs to produce similar upregulated and downregulated genes, as evidenced by the fact that 9 of the top 10 differentially expressed genes (|log_2_FoldChange| > 0.5, FDR < 0.05) ranked by false discovery rate (FDR) were shared between the two shRNA groups (**Figure S6C**). Correlation analysis of log_2_-transformed fold change (Log_2_FC) between shCaMK4_2 and shCaMK4_3 compared to shScramble further indicated that the two shRNAs produced similar transcriptomic effects (**Figure S6D**). Of all the differentially expressed genes between shCAMK4 and shScramble, 253 down-regulated genes and 349 up-regulated genes were common to both CAMK4 shRNAs (**Figure S6E**). Gene ontology (GO) analysis of the shared downregulated genes indicated that knockdown of CAMK4 suppressed genes related to the oncogenic PLC and ERK (MAPK) pathways and wound healing (possibly related to the mesenchymal subtype of GBM [45]) (**Figure 6A**). Conversely, CAMK4 knockdown upregulated biological processes related to neuronal differentiation, suggesting a shift from the stem-like GSC state (**Figure 6A**). Further unbiased gene set enrichment analysis (GSEA) using GO biological process (GO BP) datasets showed upregulation of synapse assembly across both shRNA knockdown groups (**Figure 6B, Figure S6F**). GSEA analysis with Hallmark gene sets indicated the depletion of gene sets related to epithelial mesenchymal transition (EMT), oncogenic signaling, immune responses and metabolism in both shRNA groups (**Figure S6G**). Collectively, RNA-seq analysis using several methods suggests that CAMK4 regulates tumorigenic signaling, tumor immunity and metabolism, and cellular behaviors associated with the mesenchymal subtype (EMT, wound healing, immune suppression) while preserving the stem-like GSC state and suppressing differentiation.

**Figure 6:**
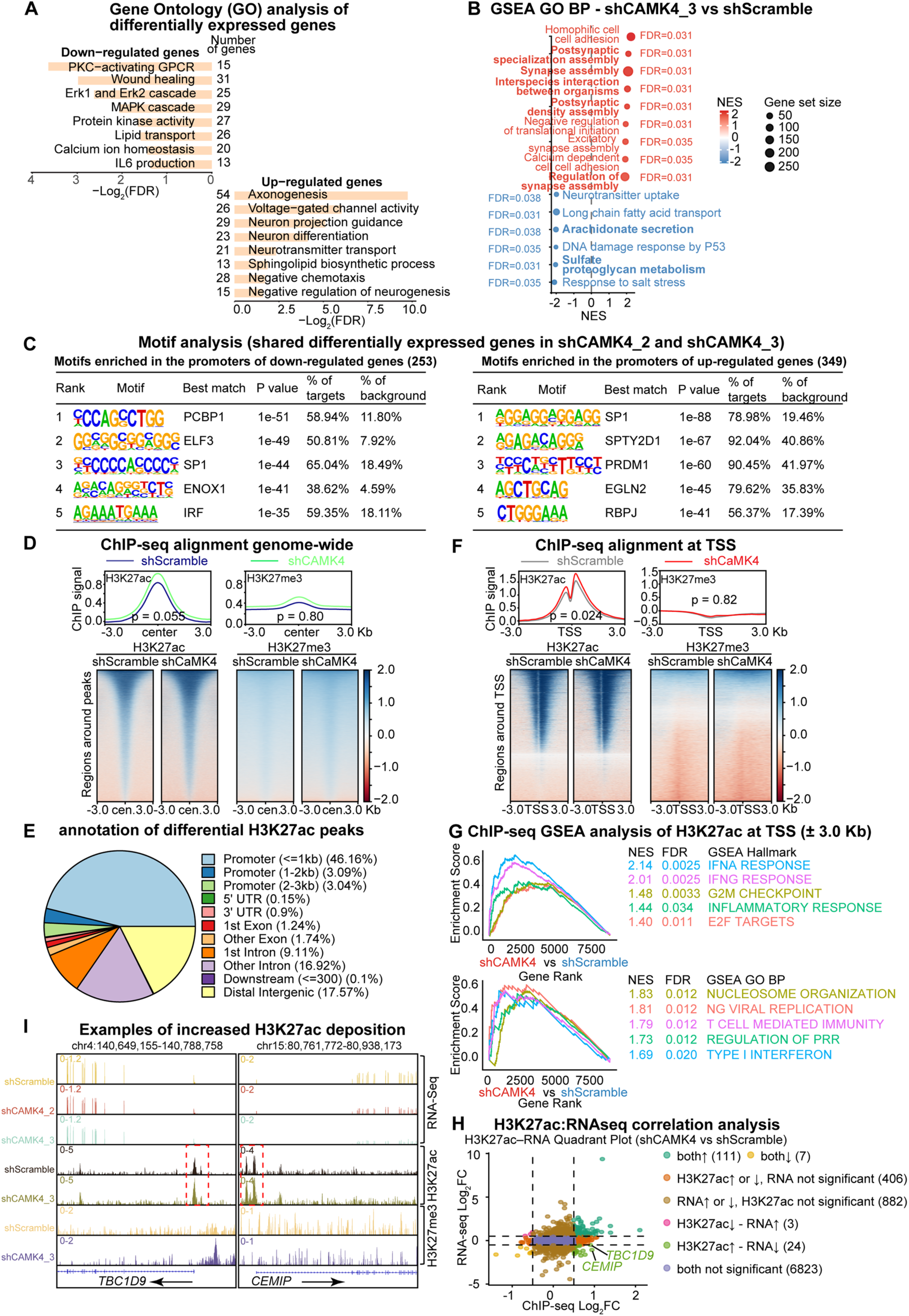
CAMK4 regulates the transcriptome of GBM cells. **(A)** Representative GO terms enriched in shared downregulated genes (top left) and upregulated genes (bottom right) from two independent shRNAs targeting CAMK4 compared to shScramble. **(B)** Bubble plots showing top GO BP terms in GSEA comparisons of the shCAMK4_3 and shScramble transcriptomes. **(C)** Motif analysis of promoter sequences of overlapping upregulated and downregulated genes between the two CAMK4 knockdown groups shows enrichment of specific motifs. **(D)** Average profile and heatmaps of H3K27ac and H3K27me3 ChIP-Seq high-confidence signals genome-wide (Wilcoxon rank-sum test; n = 2/group). **(E)** Pie chart showing annotation of high-confidence differential H3K27ac peaks between the shCAMK4 and shScramble groups. **(F)** Average profile and heatmaps of ChIP-Seq signals aligned with transcriptional start sites (TSS) ± 3 kb genome-wide (Wilcoxon rank sum test; n = 2/group). **(G)** Representation of the top-ranked Hallmark gene sets (upper) and GO terms (lower) identified by GSEA of genes ranked according to differential H3K27ac ChIP-Seq signal within 3.0 kb of the TSS in the CAMK4 knockdown group as compared to shScramble. **(H)** Quadrant plot showing correlation analysis of log_2_FC of H3K27ac ChIP signals and log_2_FC RNA-Seq of CAMK4 knockdown groups generated by DESeq2. **(I)** Representative examples of gene loci (*TBC1D9*, *CEMIP*) with decreased transcription (RNA-seq) but increased H3K27ac deposition (ChIP-seq) after CAMK4 knockdown.

To understand the mechanisms mediating this transcriptional reprogramming, we took two approaches. First, based on the possibility that CAMK4 may regulate the activity of transcription factors via phosphorylation, we performed motif analysis in the promoters of genes that were either down- or up-regulated by CAMK4 knockdown. This analysis identified motifs for several transcription factors. PCPB1, ELF3, and SP1 motifs were enriched in the promoters of downregulated genes, while SP1, SPTY2D1, and PRDM1 motifs were enriched in the promoters of upregulated genes (**Figure 6C**). The enrichment of SP1 motifs in the promoters of both up- and down-regulated genes may reflect its function as both an activator and repressor of transcription [46].

Second, we tested whether CAMK4 regulates the epigenome in PDGCs. CAMK4 in known to phosphorylate CREB Binding Protein (CREBBP or CBP or KAT3), which through its acetyltransferase activity deposits acetyl groups on lysine 27 of histone 3 (H3K27ac) to promote transcriptional activity [47–49]. We, therefore, tested whether CAMK4 knockdown alters the balance of euchromatic (H3K27ac) and heterochromatic (H3K27me3) histone modifications in PDGCs. We performed chromatin immunoprecipitation (ChIP)-Seq for H3K27ac and H3K27me3 in PDGCs with and without CAMK4 knockdown. Analysis indicated an overall increase in genome-wide H3K27ac deposition, while there was no change in the overall H3K27me3 signal (**Figure 6D**). Annotation of the H3K27ac peaks in the two groups indicated they primarily localized to the promoter regions (38.75%) (**Figure S7A**). Further annotation of the differentially called peaks by DESeq2 and ChIPSeeker between the two groups demonstrated an even stronger localization to promoters (52.29%) (**Figure 6E**). Analysis of the H3K27ac intensity in the promoter regions ± 3 kb around the transcriptional start site (TSS) genome-wide demonstrated a significantly higher level of H3K27ac in the CAMK4 knockdown group as compared to the control (**Figure 6F**). In contrast, we did not detect significant changes in H3K27me3 deposition after CAMK4 knockdown.

GSEA analysis of genes with differential H3K27ac peaks in their promoters suggested CAMK4 knockdown was positively correlated with genes within pro-inflammatory pathways, such as IFN-α and IFN-γ responses (**Figure 6G**). Analysis of the correlation between promoter-specific H3K27ac and RNA-Seq data demonstrated an overall positive association between H3K27ac and RNA transcription (**Figure 6H**). However, 24 genes demonstrated increased H3K27ac despite decreased RNA transcription (**Figure 6H**), while only 3 genes showed the opposite pattern, again pointing in the direction of dysregulated increase in H3K27ac deposition. Among these 24 genes were *TBC1D9*, which regulates TBK1-mediated innate immune responses through Ca^2+^ signaling [50], and *CEMIP,* which promotes tumor growth and invasion through the Wnt and EGFR pathways [51] (**Figure 6I**). These data suggest that CAMK4 helps regulate gene expression by selective deposition of H3K27ac at specific gene loci.

To gain insight into the increased H3K27ac deposition brought about by CAMK4 knockdown, we first grossly assessed the overall amount of H3K27ac and H3K27me3 by immunoblot in CAMK4 knockdown and control cells but could not detect any large-scale change (**Figure S7B**). Protein-protein interaction (STRING) analysis showed that CAMK4 interacts with histone deacetylases (HDAC4 and HDAC5) (**Figure S7C**) [52, 53], besides its known regulation of CREBBP. RNA-Seq indicated that CAMK4 knockdown resulted in transcriptional upregulation of *CREBBP*, *HDAC5*, and *HDAC7,* as well as the writer and eraser of the H3K27me3 modification *EZH2* and *KDM6B* respectively (**Figure S7D**), hinting at global epigenetic dysregulation.

To determine whether CAMK4 exerts effects on gene transcription and the epigenetic machinery via site-specific chromatin occupancy or diffuse actions in the nucleus, we utilized the DamID approach. DamID uses a bacterial Dam methylase fused to the protein of interest to methylate GATC motifs neighboring chromatin occupancy sites, which enables identification of bait-genome interactions, similar to ChIP-seq [54]. We transduced PDGCs with Dam-CAMK4 and Dam control lentivirus (**Figure S7E**). Dam-CAMK4 methylation enrichment analysis suggested that CaMK4 did not show any specific occupancy patterns (**Figure S7F**), a finding replicated with ChIP-Seq using Flag-tagged CAMK4 as bait (**Figure S7G**). Collectively, these data suggest that CAMK4 reprograms the transcriptome of GBM cells by regulating the activity of transcription factors and the epigenetic machinery, without targeting specific sites on chromatin itself.

### Pharmacological perturbation of P2RY1 by BPTU inhibits GBM growth

To therapeutically exploit the potent P2RY1-Ca^2+^ transient-CAMK4 pathway in GBM, we pharmacologically inhibited P2RY1 using the lipophilic allosteric inhibitor BPTU [55]. BPTU reduced Ca^2+^ transients in both the cytoplasm and nucleus (**Figures 7A-F**) and impaired the growth and viability of PDGCs *in vitro* (**Figure 7G**). BPTU also drastically inhibited tumor sphere formation in PDGCs (**Figure 7H,I**). To understand the transcriptional implications of BPTU’s effects and compare them to CAMK4 knockdown, we performed RNA-seq analysis in PDGCs treated with sublethal concentrations of BPTU or its vehicle control (**Figure 7J**). GO analysis indicated that BPTU treatment inhibited cell cycle processes, calcium ion transport, and the ERK pathway, but promoted the response to ER stress, apoptosis, innate immune responses, and differentiation programs in PDGCs (**Figure 7K**). GSEA analysis similarly showed that BPTU treatment positively correlated with interferon and inflammatory responses, but negatively correlated with chromosome separation, nuclear division and cell cycle (**Figure S8A-C**). A combined RNA-Seq analysis of CAMK4 knockdown and BPTU treatment showed overlap of 124 downregulated genes and 99 upregulated genes (**Figure 7L**). Among these overlapping differentially expressed genes, GO analysis showed positive ERK regulation, calcium ion transport, and GPCR pathways are downregulated by both P2RY1 and CAMK4, while complement activation, phagocytosis, and negative regulation of MAPK (ERK), are upregulated (**Figure 7M**).

**Figure 7:**
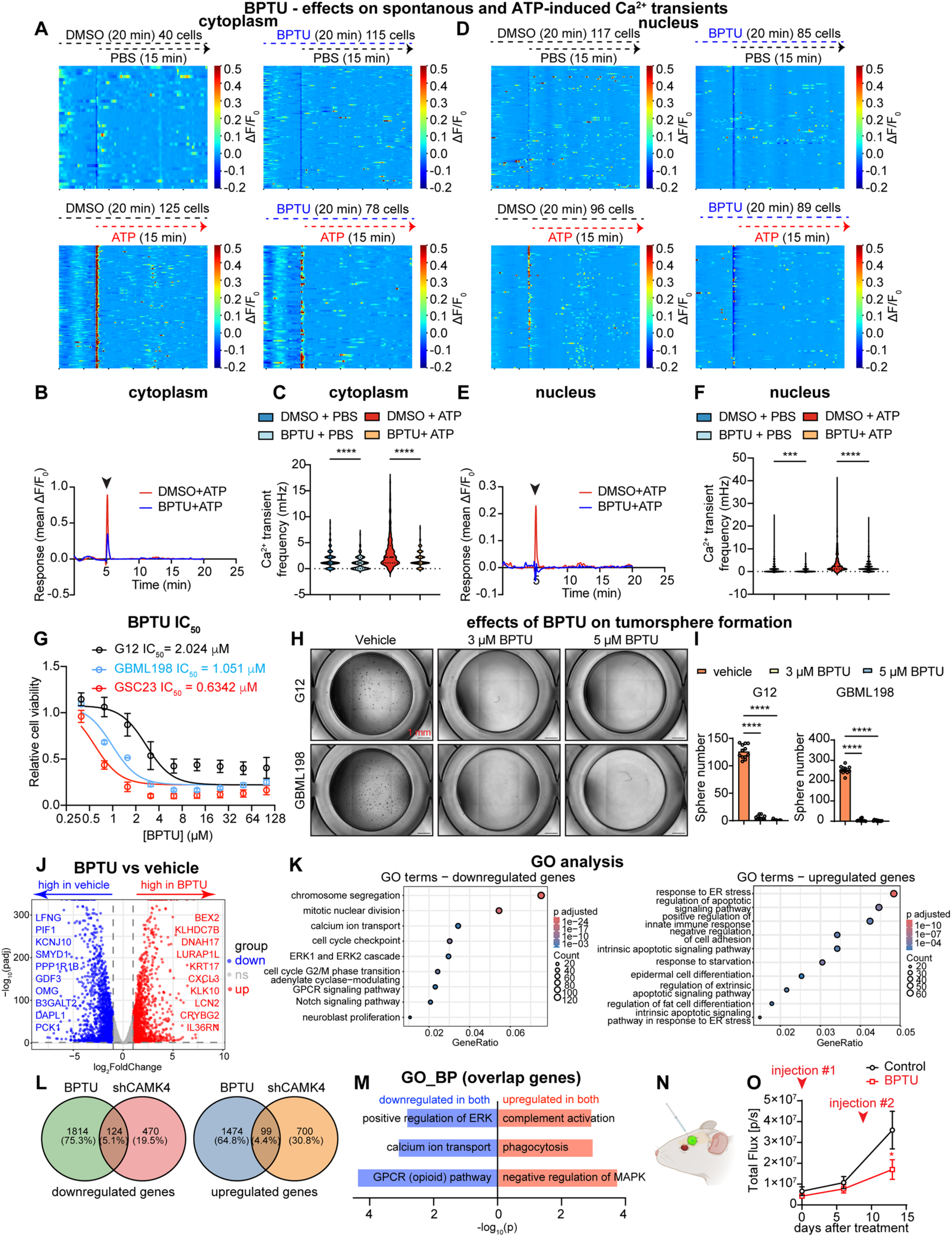
BPTU inhibits GBM growth. **(A - F)** BPTU attenuates spontaneous and ATP-induced Ca^2+^ transients in the cytoplasm and nucleus of PDGCs. **(A,D)** Heatmaps showing 20-minute recordings of ΔF/F_0_ in the cytoplasm **(A)** and nucleus **(D)** of GBML198 cells treated with 50 μM ATP in the presence or absence of 10 μM BPTU. **(B,E)** Average intensity tracings of GCaMP6s **(B)** and GCaMP6f.3xNLS **(E)** signal in GBML198 cells in response to 50 μM ATP with or without BPTU. **(C,F)** Violin plots show median and interquartile range of cytoplasmic **(C)** and nuclear **(F)** Ca^2+^ transient frequency in GBML198 cells treated with 50 μM ATP in the presence or absence of 10 μM BPTU (Kruskal-Wallis test with Dunn’s multiple comparisons test; cytoplasm: n_DMSO + PBS_ = 147, n_BPTU + PBS_ = 247, n_DMSO + ATP_ = 281, and n_BPTU + ATP_ = 233 across three recordings; nucleus: n_DMSO + PBS_ = 317, n_BPTU + PBS_ = 287, n_DMSO + ATP_ = 380, and n_BPTU + ATP_ = 313 across three recordings). **(G)** Dose-response curves of G12, GBML198, and GSC23 cells treated with BPTU, as measured with WST-8 assays. Relative cell viability was normalized to DMSO control. **(H)** Representative images of tumor sphere formation assays in G12 and GBML198 cultures treated with different concentrations of BPTU. **(I)** Bar graphs of tumor sphere formation assays in G12 (n = 12/group from three biological replicates) and GBML198 (n = 11, 11, and 10 from 3 biological replicates) cells treated with different concentrations of BPTU (one-way ANOVA with Dunnett’s multiple comparisons test; G12: F_(2, 33)_ = 1024; GBML198: F_(2, 29)_ = 1581). **(J)** Volcano plots showing differentially expressed genes in GBML198 cells treated with 5 μM BPTU vs. DMSO vehicle for 2 days. The top 10 upregulated and downregulated genes ranked by *P* value are marked (log_2_FC > 1 was used as criterion for differential expression). **(K)** Bubble plots showing representative GO terms enriched in downregulated (left) and upregulated genes (right) of BPTU-treated cells compared to DMSO-treated cells. **(L)** Venn diagram plots showing the overlap of upregulated and downregulated genes between BPTU treatment and CAMK4 knockdown. **(M)** GO terms enriched in overlapping upregulated and downregulated genes represented in **(L)**. **(N)** Schematic demonstrating the intratumoral injection of BPTU. **(O)** Line graph depicting total bioluminescence flux (y-axis) over time (x-axis) in an *in vivo* bioluminescent imaging experiment in NSG mice orthotopically implanted with luciferase-expressing GSC23 cells and treated with two intratumoral injections of vehicle (DMSO) or BPTU 8 days apart. Each injection contained 4 μg BPTU (n = 8/group, two-way ANOVA with Šídák’s multiple comparisons test, F_(2, 42)_ = 2.173). * p < 0.05, *** p < 0.001, **** p < 0.0001.

We then sought to evaluate the efficacy of BPTU *in vivo*. Administration of BPTU via oral gavage in mice for two weeks was found to produce no detectable levels in the brain, as measured by mass spectrometry (**Figure S8D,E**), suggesting lack of blood-brain barrier penetration. To mitigate this limitation, we administered BPTU intracranially within patient-derived xenografts in immunodeficient NSG mice. In a proof-of-concept study, two intratumoral injections of BPTU 8 days apart were sufficient to significantly reduce tumor burden in treated mice (**Figure 7N,O**). These data indicate that pharmacological inhibition of P2RY1 is a feasible strategy to control GBM growth.

## Discussion

Ca^2+^ transients have been identified in a variety of cell types, including plant cells [56], oocytes [2], stem cells [57], and differentiated cells [58] in a variety of tissues, where they regulate developmental and physiological programs. Ca^2+^ transients have also been implicated in oncogenesis, including gliomagenesis, although the mechanisms of their generation and their effectors remain incompletely characterized [8–10, 12, 59]. In GBM and other forms of glioma, neuron-tumor synapses entrain tumorigenic cytosolic Ca^2+^ transients mediated by neurotransmitters and their cognate receptors, including glutamate and acetylcholine, which ultimately promote tumor growth through a variety of signaling cascades and transcriptional effects [9, 11]. While such neuron-tumor interactions are relevant for tumor initiation by brain-infiltrating GBM cells [60], they cannot account for oncogenic processes within the tumor core, which is devoid of neuronal inputs. There, instead, networks of tumor cells exhibit tumor cell-autonomous Ca^2+^ transients with similar oncogenic potential as those induced by neuronal synapses [12]. In essence, these findings suggest that emerging properties of tumor cellular networks produce signals sufficient for the generation of self-sustaining oncogenic Ca^2+^ transients [12, 60]. However, what these signals are and how Ca^2+^ signaling cascades influence transcriptional programs remains unclear.

Our work provides answers to both questions. Given that GBM cells express neurotransmitter /neuromodulator receptors [61], we performed a functional neuromodulator screen to assess influences on tumor cell-autonomous Ca^2+^ transients. We discovered that extracellular purines ATP and ADP, which are known to be orders of magnitude more abundant in the extracellular space of tumors relative to healthy tissues [62], are potent inducers of Ca^2+^ transients in GBM via activation of metabotropic Gq- coupled P2RY1 receptors. This was consistent with our observation that Ca^2+^ transients did not correlate with changes in the membrane potential of GBM cells, as one would expect if ionotropic receptors were involved, but instead depended on IP3 receptor (IP3R)-mediated Ca^2+^ release from internal stores. While it is incontestable that cytosolic Ca^2+^ transients act through multiple effectors to influence cellular phenotypes and transcriptional programs, we discovered that these IP3R-dependent transients were represented robustly in the nucleus and that P2RY1 activation was both necessary and sufficient for these nuclear Ca^2+^ transients. Within the nucleus of GBM cells, Ca^2+^ transients activate the Ca^2+^/calmodulin-dependent kinase CAMK4, which in turn regulates transcriptional programs related to stemness/differentiation, anti-tumor immunity, metabolism and mesenchymal phenotypes, via effects on the epigenetic machinery and likely post-translational modifications of transcription factors as well. CAMK4 additionally regulates transcription of ribosomal DNA, thus influencing not just the transcriptome, but the translatome as well. These findings provide a detailed mechanistic explanation for the transduction of the extracellular purinergic signals to the nuclear machinery. Importantly, this novel oncogenic mechanism is targetable therapeutically. Our expression profiling of P2RY1 and CAMK4 showed that the former is significantly upregulated in GBM relative to healthy brain tissue, as opposed to the latter, which is expressed in neurons and is required for neuronal plasticity [43]. Given the anticipated favorable therapeutic window for P2RY1, we tested the P2RY1-specific allosteric inhibitor BPTU [55], previously used in experimental rodent models of Alzheimer’s disease [63], for anti-tumor effects. Our pharmacokinetic interrogation of BPTU showed no penetration into brain tissue, which prompted us to test whether direct intratumoral delivery impairs tumor growth. Indeed, intermittent intratumoral administration of BPTU slowed tumor growth in our orthotopic patient-derived GBM xenograft models in mice, raising the possibility that optimization of the delivery can produce long-lasting anti-tumor benefits and improved survival.

Previous reports have suggested that P2RY1 exerts tumor suppressive effects in a variety of malignancies, such as prostate cancer, gastric cancer and even GBM [64–67]. In addition, eATP/ADP are known to serve as DAMPs (damage-associated molecular patterns) sensed by purinergic receptors, such as P2RY1 to trigger inflammatory responses [68]. Our findings, however, indicate that eATP/ADP act as potent tumor-promoting signals through P2RY1 receptors, nuclear Ca^2+^ transients and CAMK4 activation to bias the transcriptome and epigenome of GBM cells toward proliferation, stem-like states, and immune evasion. This mechanism, along with the established action of ectonucleotidases to break down eATP/ADP to adenosine, a potent immunosuppressive signal acting on immune cells in the tumor microenvironment [69], establish extracellular purines as a critical tumorigenic driver. Collectively, our work reveals a novel and pharmacologically targetable oncogenic mechanism in GBM and possibly other malignancies, in which extracellular purines regulate the transcriptome and epigenome of tumor cells via nuclear Ca^2+^ transients.

## Materials and Methods

### Cell lines and cell culture

Patient-derived GBM cultures (PDGCs) (GBML109, GBML137, GBML177 and GBML198) were established and maintained as previously described with slight modifications [70–72]. In brief, fresh IDH-wild type GBM specimens were obtained from patients after informed consent (NYU IRB study 12-01130). Specimens were mechanically minced using autoclaved surgical scissors followed by enzymatic dissociation using Accutase (Innovative Cell Technologies, Cat# AT104) in 37°C incubator for about 20 minutes with pipetting up and down every 5 minutes until the minced specimens didn’t block P1000 tips. Red blood cells were removed using Red Blood Cell Lysis Buffer (Sigma, Cat# 11814389001). Finally, PDGCs were filtered with 70 μm cell strainer (Sigma, Cat# CLS352350) before long-term culture in Neurobasal medium (Gibco, Cat# 21103049) supplemented with 1:1000 N2 (Gibco, Cat# 17-502-049), 1:1000 B27 (Gibco, Cat# 12587010), nonessential amino acids (Gibco, Cat# 11140050), GlutaMax (Gibco, Cat# 35050061), 20 ng/ml recombinant basic Fibroblast Growth Factor (bFGF; PeproTech, Cat# 10018B1MG), 20 ng/ml Epidermal Growth Factor (EGF; PeproTech, Cat# AF100151MG) and Penicillin-Streptomycin (Gibco, Cat# 15140122). GSC23 and G12 PDGCs were gifts from Drs. Erik Sulman and Jann Sarkaria, respectively. PDGCs were cultured as floating tumor spheres in low-attachment vessels and were dissociated with Accutase for passaging.

Neural stem cells (NSCs) were derived from H9 human embryonic stem cells (hESCs) using a previously established protocol [73]. All hESC related experiments were approved by New York University (NYU) Embryonic Stem Cell Research Oversight Committee (ESCRO) (Protocol 14-00267). Human ESCs were cultured on mouse embryonic fibroblasts (Thermo Fisher, Cat# A34962) using gelatin (Sigma, Cat# G1890-100G)-coated cell culture dishes. NSCs were cultured on Poly-L-Ornithine (R&D systems, Cat# 3436-100-01) and Laminin (Corning, Cat# 354232)-coated dishes using Dulbecco’s modified Eagle’s medium/Nutrient Mixture F-12 (DMEM/F-12; Gibco, Cat# 11330057) supplemented with N2, 20 µg/mL Insulin (Sigma, Cat# I0516), 1:1000 B27, 1.6 g/L glucose (Thermo Fisher, Cat# A2494001) and 20 ng/mL EGF and bFGF.

HEK293T (Takara, Cat# 632180) cells were cultured in Dulbecco’s modified Eagle’s medium (DMEM; Gibco, Cat# 11965-118) supplemented with 10% fetal bovine serum (FBS; Corning, Cat# 35-015-CV), Penicillin-Streptomycin and sodium pyruvate (Gibco, Cat# 11360070). All cells were cultured in humidified 37°C incubator balanced with 21% O_2_ and 5% CO_2_.

### Plasmids and molecular cloning

For all shRNA plasmids cloning, pLKO.1 - TRC control was a gift from David Root (Addgene plasmid # 10879; RRID:Addgene 10879). A published shScramble sequence was chosen as a negative control hairpin [74]. shP2RY1_1, shP2RY1_3, shCAMK4_2, shCAMK4_3, shCAMK2D_1, shITPR2_2, and shITPR2_3 were designed using GPP Web Portal (https://portals.broadinstitute.org/gpp/public/). pLKO.1 - TRC control was digested with AgeI-HF (NEB, Cat# R3552S) and EcoRI-HF (NEB, Cat# R3101S) for 2 hours at 37°C. Digested products were purified using QIAquick Gel Extraction Kit (Qiagen, Cat# 28704) after gel electrophoresis. ShRNA oligonucleotide annealing mixture consisting of forward and reverse oligonucleotides (100 nmol/mL each) and 1x annealing buffer (10 mM Tris-HCI, pH 7.5, 0.1 M NaCI and 1 mM EDTA) was heated to 95°C for 10 minutes on a dry heater, then cooled slowly to room temperature after the heater was turned off. Annealed oligonucleotides were ligated into purified fragments with T4 DNA ligase (NEB, Cat# M0202M) for 20 minutes at room temperature. Ligation products (1 μL) were transformed into homemade competent Stbl3 competent cells generated using Mix & Go! E.coli Transformation Kit (Zymo research, Cat# T3001).

All the plasmids are summarized in **Tables 1** and **2**, with primer sequences listed in **Table 3**. Restriction digest fragments and PCR-amplified inserts were purified using QIAquick Gel Extraction Kit (Qiagen, Cat# 28704) after gel electrophoresis. Ligation was performed using T4 ligase as described above. All the plasmids used in the paper were verified by Sanger sequencing or whole-plasmid Nanopore sequencing.

**TABLE 1:** Plasmids.

| Plasmid | Source | Identifier |
| --- | --- | --- |
| hSyn:GCaMP6f.3xNLS.HA | Hilmar Bading | N/A |
| pHAGE-RSV-tdTomato-2A-GCaMP6s | Darrell Kotton | Addgene plasmid # 80316 |
| pMD2.G | Didier Trono | Addgene plasmid # 12259 |
| psPAX2 | Didier Trono | Addgene plasmid # 12260 |
| pCX-SpiCee-NLS | Xavier Nicol | Addgene plasmid # 140900 |
| pCX-mutSpiCee | Xavier Nicol | Addgene plasmid # 140903 |
| pLV-EF1a-IRES-Blast | Tobias Meyer | Addgene plasmid # 85133 |
| pLKO.1 - TRC cloning vector | David Root | Addgene Plasmid #10878 |
| pDisplay-GRAB_ATP1.0-IRES-mCherry-CAAX | Yulong Li | Addgene Plasmid # 167582 |
| pLenti PGK Blast V5-LUC (w528-1) | Eric Campeau,<br>Paul Kaufman | Addgene Plasmid # 19166 |
| pHIV-LUC-mCherry | Timothy Chan |  |
| 722:hSyn:CaMBP4.Flag.mCherry | Hilmar Bading |  |
| pLVX-EF1α-mCherry-N1 | - | TAKARA Plasmid# 631986 |
| pLenti-SFFV-mScar1-Dam | Andrea Brand |  |

**TABLE 2:** PCR-based cloning.

| Insert | Backbone | Cloning method or restriction enzymes | Final plasmid |
| --- | --- | --- | --- |
| Not-GCaMP6f.3xNLS.HA-BamHI | pHAGE-RSV-tdTomato-2A-GCaMP6s | Not-HF, BamHI-HF | pHAGE-GCaMP6f.3xNLS.HA |
| shITPR2_2-F, shITPR2_2-R | pLKO.1 - TRC cloning vector | EcoRI-HF, AgeI-HF | pLKO.1-shITPR2_2 |
| shITPR2_3-F, shITPR2_3-R | pLKO.1 - TRC cloning vector | EcoRI-HF, AgeI-HF | pLKO.1-shITPR2_3 |
| CAMK4 | pLenti-SFFV-mScar1-Dam | Gibson Assembly | pLenti-SFFV-mScar1-Dam.CaMK4 |
| mRFP1.NLS | pLV-EF1a-IRES-Blast | Gibson Assembly | pLV-EF1a-mRFP1.NLS-IRES-Blast |
| SpiCee.mRFP1.NLS | pLV-EF1a-IRES-Blast | Gibson Assembly | pLV-EF1a-SpiCee.mRFP1.NLS-IRES-Blast |
| mutSpiCee.mRFP1.NLS | pLV-EF1a-IRES-Blast | Gibson Assembly | pLV-EF1a-mutSpiCee.mRFP1.NLS-IRES-Blast |
| ATP1.0-IRES-mCherry | pHAGE-RSV-tdTomato-2A-GCaMP6s | Gibson Assembly | pHAGE-RSV-GRAB_ATP1.0-IRES-mCherry |
| shScramble-F, shScramble-R | pLKO.1 - TRC cloning vector | EcoRI-HF, AgeI-HF | pLKO.1-shScramble |
| shP2RY1_1-F, shP2RY1_1-R | pLKO.1 - TRC cloning vector | EcoRI-HF, AgeI-HF | pLKO.1-shP2RY1_1 |
| shP2RY1_3-F, shP2RY1_3-R | pLKO.1 - TRC cloning vector | EcoRI-HF, AgeI-HF | pLKO.1-shP2RY1_3 |
| shCAMK4_2-F, shCAMK4_2-R | pLKO.1 - TRC cloning vector | EcoRI-HF, AgeI-HF | pLKO.1-shCAMK4_2 |
| shCAMK4_3-F, shCAMK4_3-R | pLKO.1 - TRC cloning vector | EcoRI-HF, AgeI-HF | pLKO.1-shCAMK4_3 |
| shCAMK2D_1-F, shCAMK2D_1-R | pLKO.1 - TRC cloning vector | EcoRI-HF, AgeI-HF | pLKO.1-shCAMK2D_1 |
| BclI-CAMK4-Mfe I | pLV-EF1a-IRES-Blast | EcoRI-HF, BamHI-HF | pLV-EF1a-CAMK4-IRES-Blast |
| BamHI-CAMK2D-EcoRI | pLV-EF1a-IRES-Blast | EcoRI-HF, BamHI-HF | pLV-EF1a-CAMK2D-IRES-Blast |
| P2A | pHIV-LUC-mCherry | Gibson Assembly | pHIV-luc-P2A-mCherry |
| BamHI-mCherry-XbaI | pLVX-EF1α-mCherry-N1 | BamHI-HF, XbaI-HF | pLVX-EF1α-CaMBP4.Flag.mCherry |
| EcoRI-mCherry-XbaI | pLVX-EF1α-mCherry-N1 | EcoRI-HF, XbaI-HF | pLVX-EF1α-mCherry.NLS |
| BclI-flag-CAMK4-MfeI | pLV-EF1a-IRES-Blast | EcoRI-HF, BamHI-HF | pLV-EF1a-Flag.CAMK4-IRES-Blast |

**TABLE 3.**
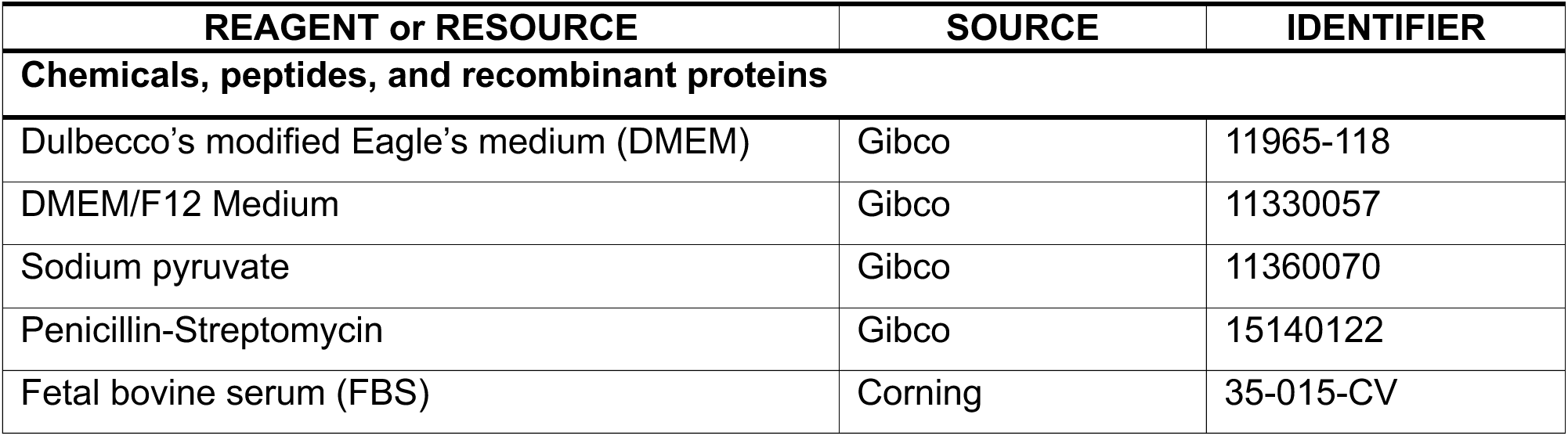
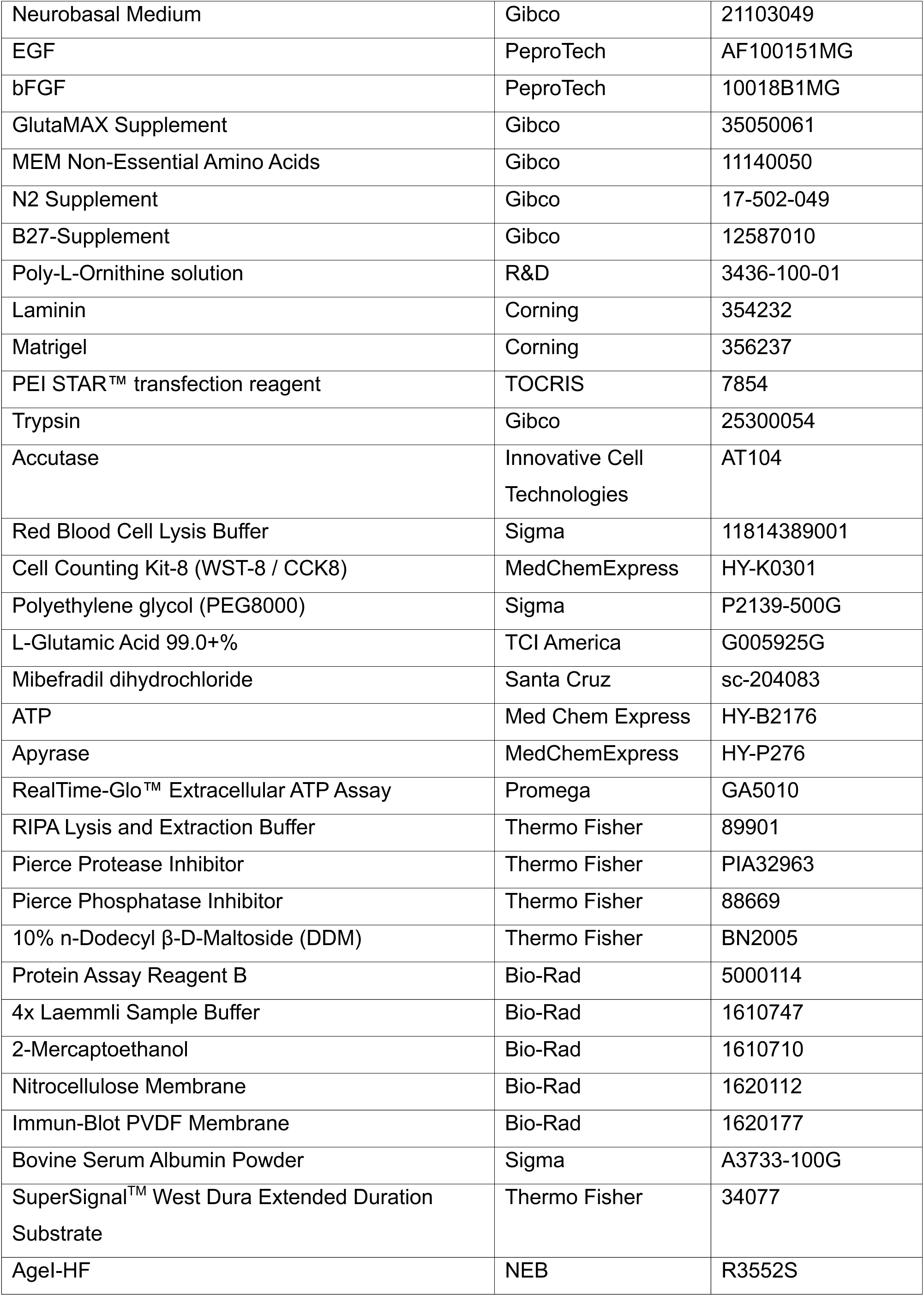
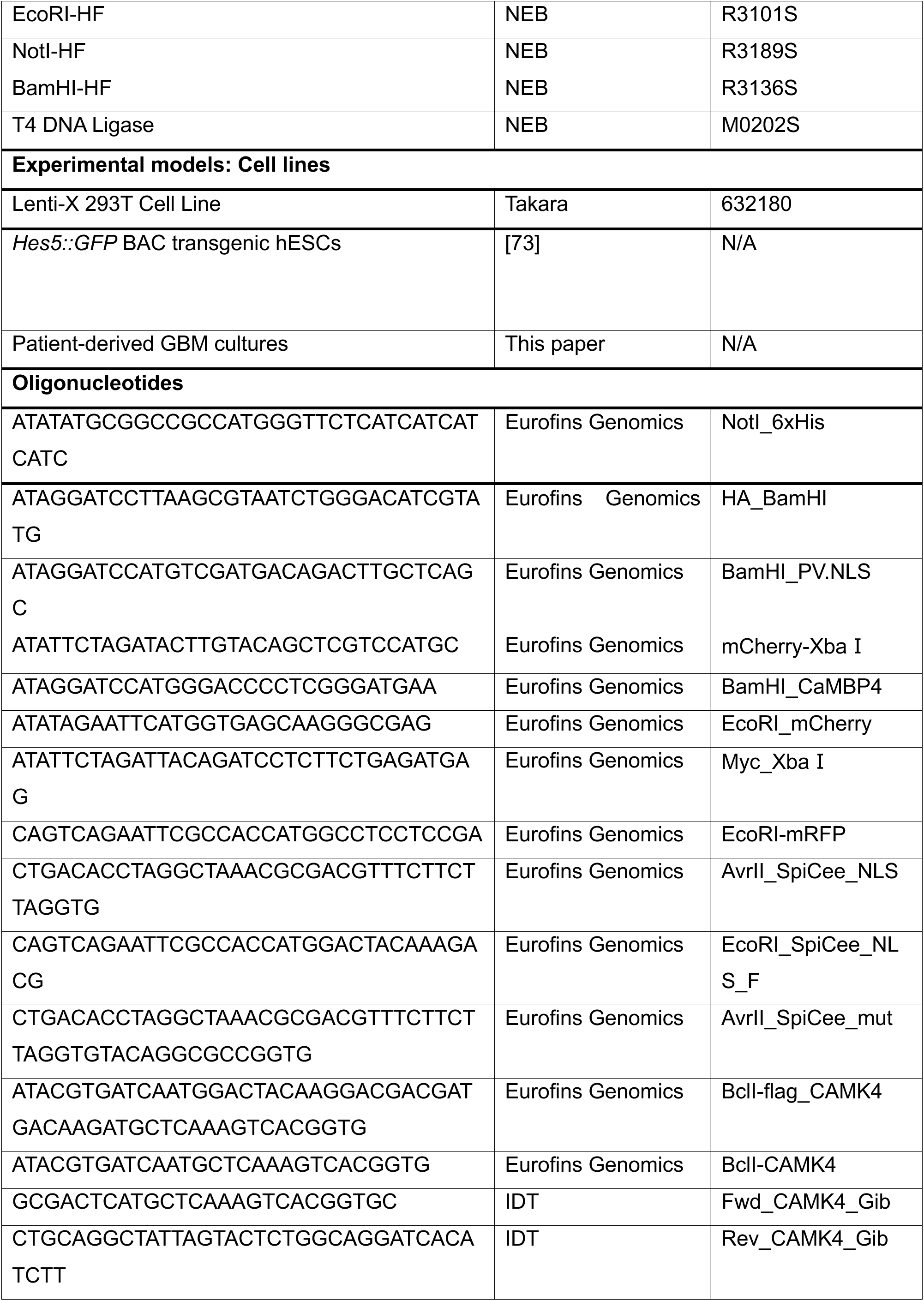
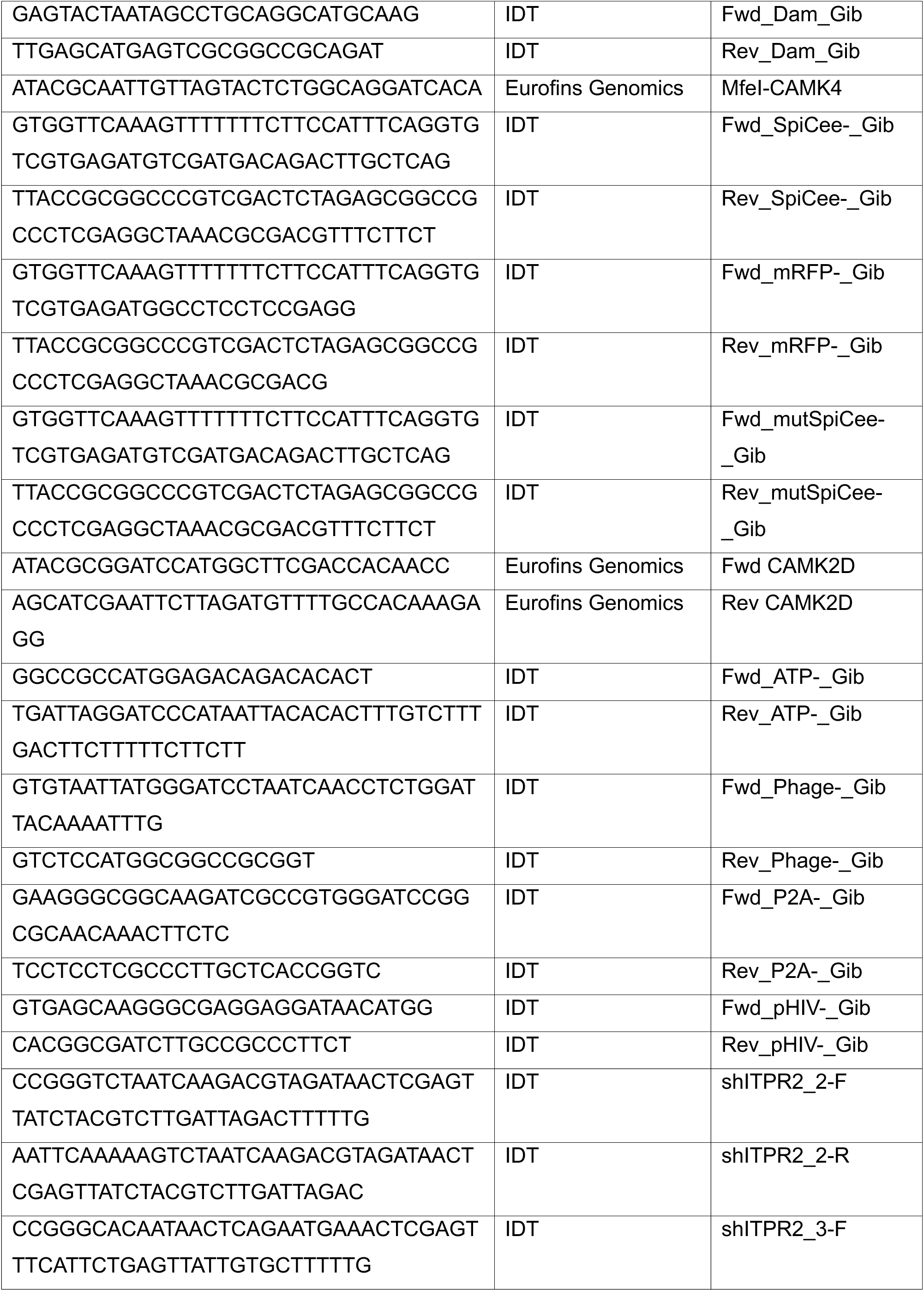
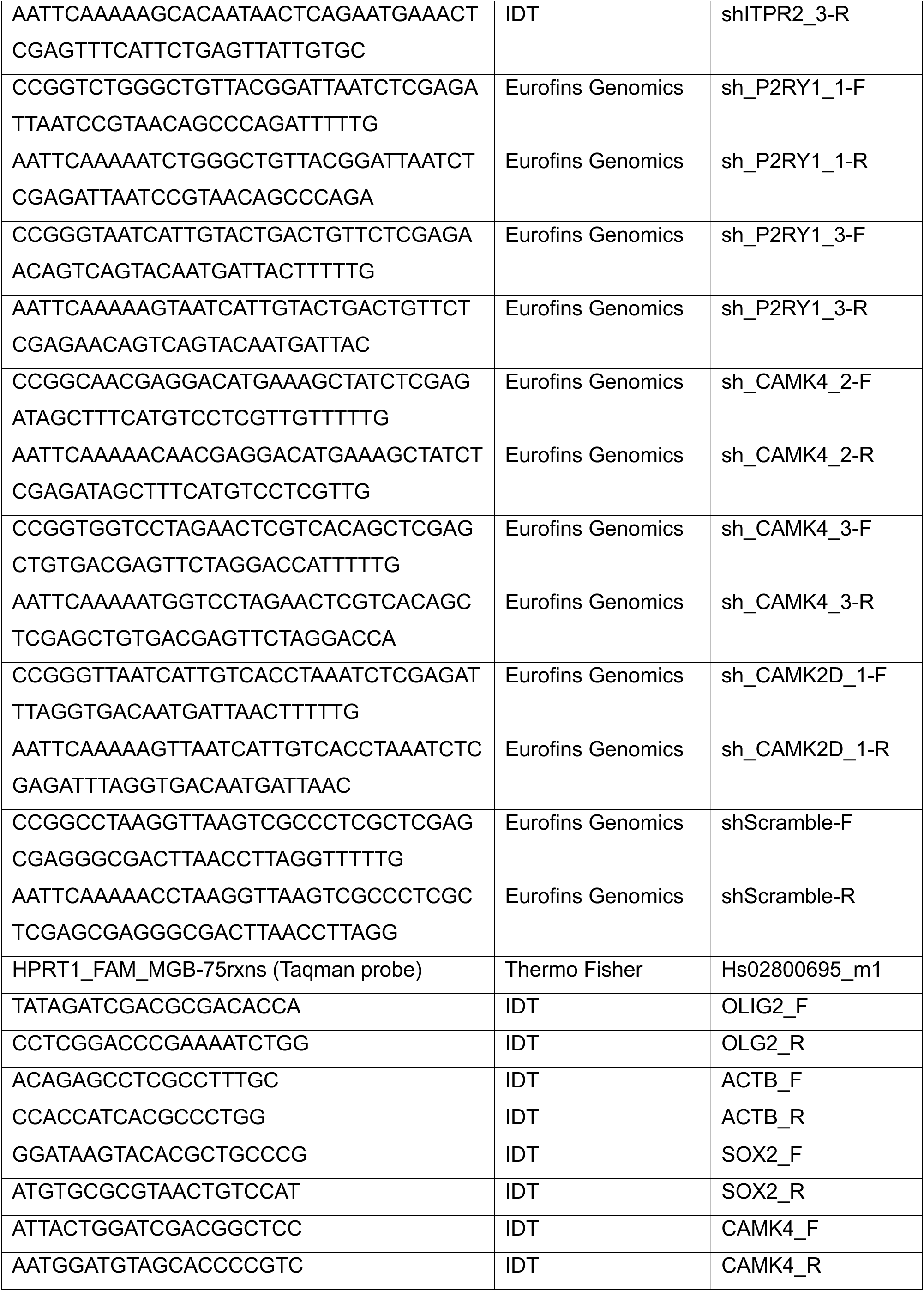
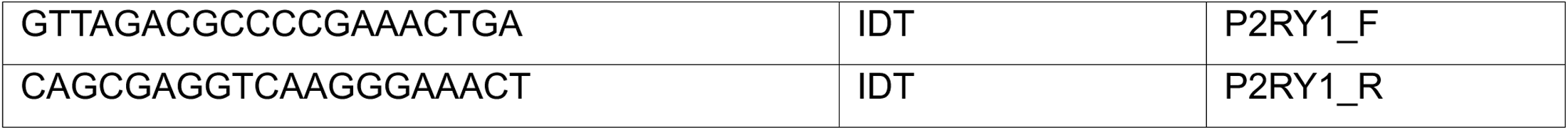

pHAGE-RSV-tdTomato-2A-GCaMP6s was a gift from Darrell Kotton (Addgene plasmid # 80316; RRID:Addgene 80316) [13]. hSyn:GCaMP6f.3xNLS.HA was a gift from Hilmar Bading [75]. pLV-EF1a-IRES-Blast was a gift from Tobias Meyer (Addgene plasmid # 85133; RRID:Addgene_85133). pCX-SpiCee (Addgene plasmid # 140836; RRID:Addgene_140836) and pCX-mutSpiCee (Addgene plasmid # 140903; RRID:Addgene_140903) were gifts from Xavier Nicol. hSyn:mCherry.NLS.myc, hSyn:PV.NLS.mCherry, and hSyn:CaMBP4.Flag.mCherry were gifts from Dr. Hilmar Bading.

### Lentivirus production and transfection

psPAX2 (Addgene, Cat# 12260) and pMD2.G (Addgene, Cat# 12259) were gifts from Didier Trono. Lentivirus was produced by co-transfecting HEK293T cells with 1.64 pmol transfer plasmids, 1.3 pmol psPax2 and 0.72 pmol pMD2.G in each 10-cm dish using PEI STAR™ transfection reagent (TOCRIS, Cat# 7854). Lentivirus was collected from the cell culture supernatant 24, 48, and 72 hours after transfection. Lentivirus supernatant was concentrated using 4X Lentivirus Concentrator Solution (40% PEG-8000 weight/volume, 1.2 M NaCl, and 1X PBS). For lentivirus infection, dissociated PDGCs in single-cell suspension were seeded in 6-well plates at a density of 300,000 cells per well and transduced with lentivirus for 24 hours followed by changing the medium. Infected cells were selected by either fluorescence activated cell sorting (FACS) with the SH800Z sorter (Sony Biotechnology) or by puromycin selection (2 μg/mL) for at least 4 days, as appropriate.

### In vitro Ca^2+^ imaging, pharmacology assays, and Ca^2+^ transient analysis

Cytosolic Ca^2+^ imaging utilized GCaMP6s, while nuclear Ca^2+^ signal was imaged with GCaMP6f.3xNLS. Representative optical recordings are shown as videos in **Supplementary Videos**. GBM cells were plated at a density of 1x 10^5^ cells/dish on 10 mm imaging dishes (World Precision Instruments, FD3510-100) pre-coated with Poly-L-Ornithine and Laminin. All the imaging experiments were performed a minimum of 48 hours following seeding on imaging dishes and in DMEM/F-12 supplemented with N2, B27, nonessential amino acids, GlutaMax, 20 ng/ml bFGF, 20 ng/ml EGF and Penicillin-Streptomycin. For pharmacological assays, L-Glutamic Acid (TCI America, Cat# G005925G) was always freshly prepared because of the instability of glutamic acid in water. Other inhibitors or antagonists were dissolved in 5 μL (3% of the total volume) PBS followed by adding to the imaging dish. Concentrations of the neurotransmitters/neuromodulators used in the screen were as follows: 100 μM ATP, 1 mM acetylcholine (Ach), 100 μM norepinephrine (NE), 100 μM dopamine (DA), 100 μM serotonin (5-HT), and 100 μM GABA. Ca^2+^ imaging was performed using an inverted epifluorescent Olympus microscope with a 40x oil dry objective. For time-lapse recordings, images of GFP fluorescence were acquired at 20% LED power at 2-second intervals. Every time series was recorded for 20 minutes in total, unless otherwise specified. Neurotransmitters and other purinergic agonists were added at the 5-minute point unless otherwise specified.

To evaluate the impact of P2RY1 antagonist BPTU (MedChemExpress, Cat# HY-13831) on Ca^2+^ transients, cells were pre-incubated in 10 μM BPTU for 30 minutes prior to imaging. Similarly, cells were pre-incubated in G_q_ antagonist YM354890 (Tocris, Cat# 7352) for 1 hour prior to imaging. For experiments involving caged(ci)-IP_3_ (Tocris, Cat# 6210), cells were pre-incubated in 1 μM ci-IP_3_ for 30 minutes prior to imaging. During recording, IP_3_ was uncaged using UV light (405 nm) at the desired experimental timepoint.

Ca^2+^ transient characterization and visualization were performed with ImageJ, Cellpose [76], and Python. All the scripts for ImageJ and Python can be found on GitHub (Shuai-NY/Calcium-analysis). In brief, ΔF/F_0_ was determined as fluorescence traces from each region of interest (ROI) as defined by Cellpose normalized using a rolling median baseline correction, for which a 120-frame window was used to estimate the local baseline. Traces were denoised using a Savitzky-Golay filter. Ca^2+^ peaks were then detected using an approach based on the SciPy find_peaks and peak_widths functions, with predefined thresholds of minimum prominence ≥ 0.05, amplitude ≥ 0.05, and inter-peak distance ≥ 1 frame. Peak widths were quantified at half-prominence, and only events with physiologically plausible durations (1-30 seconds) were retained. For each retained peak, we extracted data regarding peak amplitude, rise time, decay time, and area under the curve (AUC). Frequency was computed as the number of retained peaks divided by the recording duration.

### In vitro confocal imaging of GRAB_ATP_ in PDGCs

PDGCs expressing ATP sensor GRAB were plated at a density of 1×10^5^ cells/dish on 10 mm imaging dishes (World Precision Instruments, FD3510-100) pre-coated with poly-L-ornithine and laminin. Twenty-four hours after plating, apyrase was added at a final concentration of 10 U/mL. An equal volume of molecular grade water was added to vehicle control dishes. A confocal microscope (Zeiss LSM800) was used to take Z-stacked images of GFP fluorescence at 0.5 µm intervals immediately prior to the addition of apyrase or water, as well as 5 and 15 minutes later. Mean fluorescence of the z-stack projection of each image was determined using Fiji.

### Quantification of extracellular ATP

PDGCs were plated in white opaque 96-well plates at a density of 2×10^4^ cells/well. Twenty-four hours later, apyrase was added to the wells at concentrations ranging from 0-10 U/mL. After 5 minutes, RealTime-Glo™ Extracellular ATP Assay (Promega) substrate was added to each well at a final dilution of 1:4, and luminescence was measured using a Synergy H1 Plate Reader (BioTek).

### Immunofluorescence staining

GBM cells were plated on cover slips (Fisher Scientific, Cat# 50143822) pre-coated with Poly-L-Ornithine and Laminin. Cells were fixed with 4% paraformaldehyde (PFA; Thermo Fisher, Cat# 047340.9M) followed by permeabilizing with 0.1% Triton™ X-100 (Sigma, Cat# T8787) in PBS. Cells were blocked with 5% Bovine Serum Albumin (BSA; Sigma, Cat# A3733) in PBS for 30 minutes. Primary antibody diluted (**Table 4**) in PBS containing 3% BSA was incubated overnight. Cells were then washed using 0.05% Tween-20 (Sigma, Cat# P9416) in PBS (PBS-T) for 5 minutes three times and stained with secondary antibodies (see **Table 4**) for 1 hour at room temperature in the dark. Cells were washed with PBS-T for 5 minutes three times followed by staining with DAPI or Hoechst 33342 (Life Technologies, Cat# H3570) for 15 minutes. Coverslips were mounted with ProLong Gold antifade reagent (Thermo Fisher, Cat# P36934) on microscope slides (Fisher Scientific, Cat# 22042915) for imaging. Pictures were taken using an LSM800 Microscope with Airyscan (Zeiss).

**TABLE 4.**
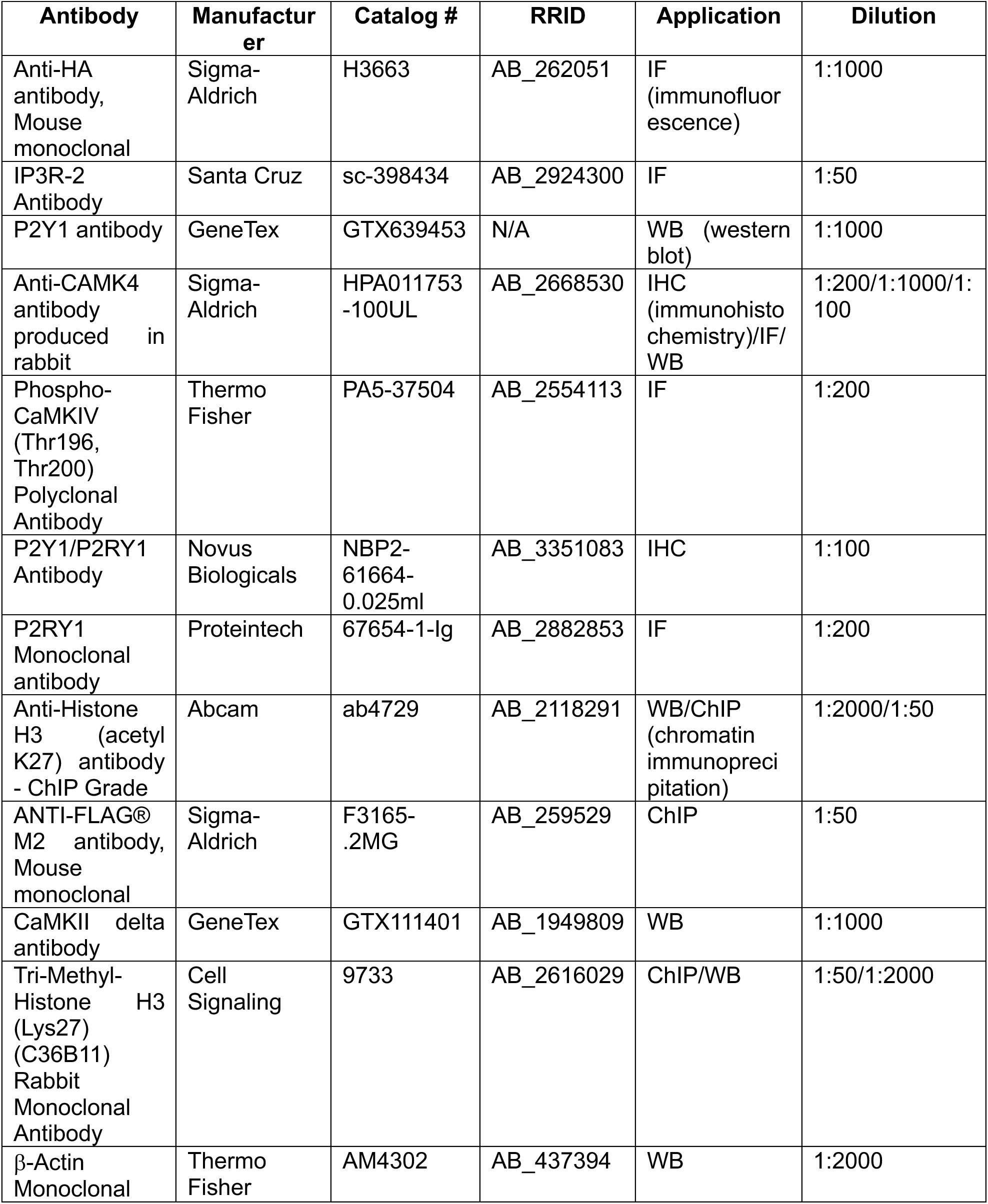
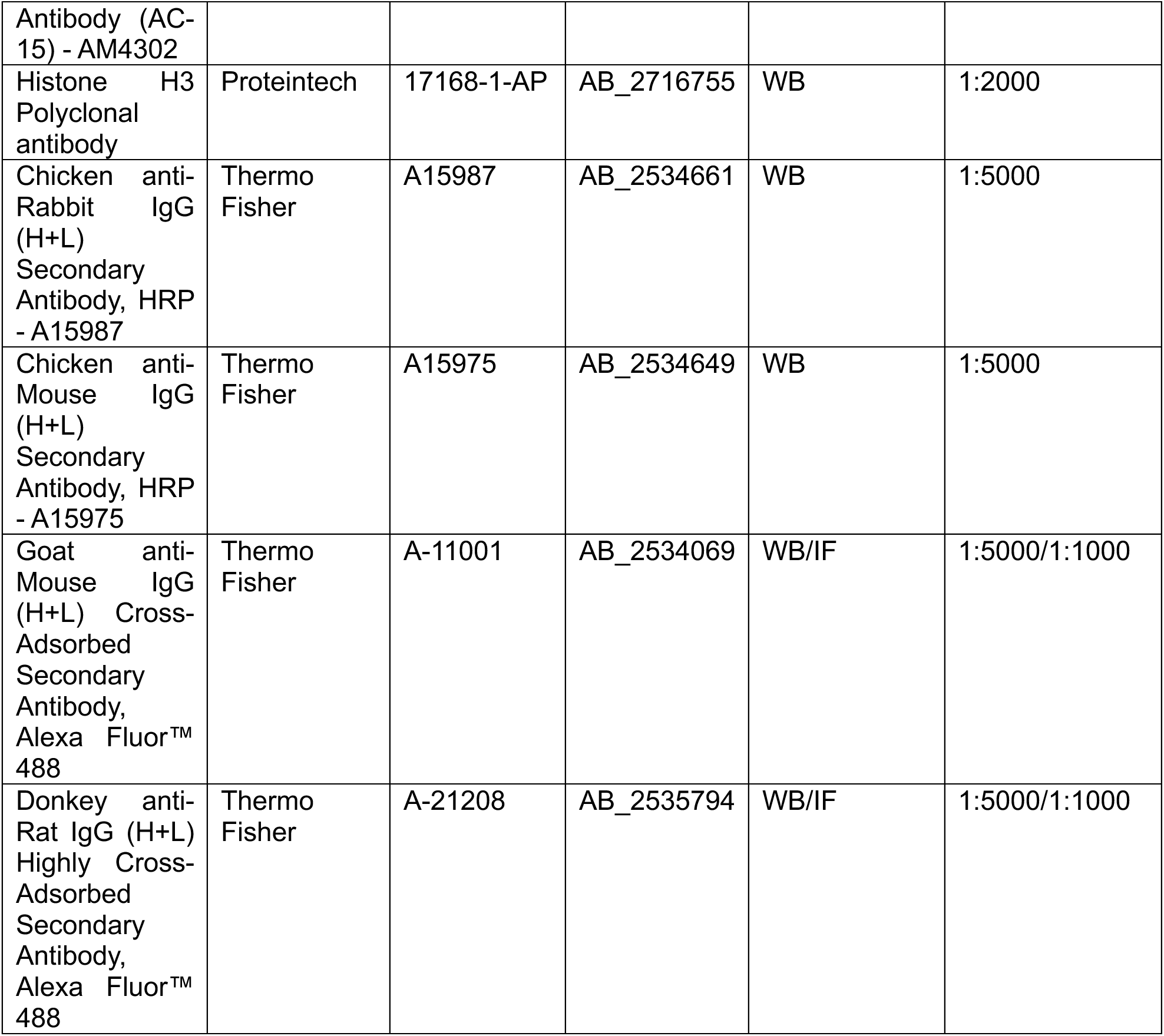

### Protein extraction and Western blot

Cells were lysed using RIPA Lysis and Extraction Buffer (Thermo Fisher, Cat# 89901), supplemented with protease inhibitor (Thermo Fisher, Cat# PIA32963) and phosphatase inhibitor (Thermo Fisher, Cat# 88669). N-dodecyl β-D-maltoside (DDM; Thermo Fisher, Cat# BN2005) was also added at 10% for membrane proteins. Lysates were incubated on ice for 10 minutes followed by ultrasonification using a Vibra-Cell Ultrasonic Processor (Sonics) and centrifugation at 16,000 g for 10 minutes at 4°C. The supernatant was transferred to clean tubes for quantification of protein concentration by DC Protein assay (Bio-Rad, Cat# 5000114). Samples were denatured with 4x Laemmli Sample Buffer (Bio-Rad, Cat# 1610747) with 10% 2-mercaptoethanol (Bio-Rad, Cat# 1610710) and incubated at 95°C for cytoplasmic and nuclear proteins and room temperature for membrane proteins. Protein lysates (10-30 μg) were then resolved by SDS PAGE. Proteins were then transferred from gels to nitrocellulose (Bio-Rad, Cat# 1620112) or PVDF (Bio-Rad, Cat# 1620177) and blocked using bovine serum albumin (Sigma, Cat# A3733-100G) for 1 hour at room temperature. Samples were then incubated with desired primary antibodies (see **Table 4**) at 4°C overnight. Tris-buffered saline with 0.1% Tween 20 (TBST) was then used to wash the samples three times for 10 minutes each at room temperature. The samples were then incubated with desired secondary antibodies toward chemiluminescent or fluorescent visualization (see **Table 4**) at room temperature for 1 hour. Horseradish peroxidase (HRP) signals were developed using SuperSignal^TM^ West Dura Extended Duration Substrate (Thermo Fisher, Cat# 34077) and captured, similarly to fluorescent signals, by the iBright Imaging System (Thermo Fisher).

### Reverse transcription and qRT-PCR

For qRT-PCR using SYBR Green chemistry, RNA was extracted using Monarch® Total RNA Miniprep Kit (NEB, Cat# T2010S), and 1 μg RNA was used to generate cDNA using High-Capacity cDNA Reverse Transcription Kit (Thermo Fisher, Cat# 4368814). Reaction mixes were then prepared using PowerUP SYBR Green Master Mix (Thermo Fisher, Cat# A25742). For TaqMan qRT-PCR, Cells to CT Kit (Cat #1729, ThermoFisher Scientific) was used to obtain RNA and qPCR was performed using TaqMan®FAM™ dye-labeled probes including *CAMK2D* (Thermo Fisher, Cat # Hs00943538_m1), and *HPRT1* (Thermo Fisher, Cat# Hs02800695_m1). qRT-PCR was performed on a Quantstudio 3 or StepOne Real-Time PCR System (Thermo Scientific).

### Tumor sphere formation assay

PDGCs in culture were enzymatically dissociated into single cells using Accutase and counted manually or using Countess II (Thermo Fisher) with Trypan blue (Thermo Fisher, Cat# T10282) staining. Aliquots of 1000-2000 cells in 100 μL complete culture medium were dispensed into non-adherent 96-well plates (Corning, Cat# 3370). Every two days, we added 10 μL fresh culture medium with EGF and bFGF. Tumor spheres were quantified two weeks after seeding. 96-well plates were then imaged by automated well scanning using an EVOS M7000 Cell Imaging System (Thermo Fisher) at 4X magnification. Pictures were processed and tumor spheres larger than 75 μm were counted using ImageJ software with setting circularity to 0.20 - 1. Fiji scripts used to preprocess tumor sphere images can be found on GitHub (https://github.com/Shuai-NY).

### Extreme limiting dilution assays

PDGCs were plated in 96-well plates at 6 separate concentrations with 10 replicates: 50 cells/well, 40 cells/well, 20 cells/well, 10 cells/well, 5 cells/well, 2 cells/well. PDGCs were fed every 2 days with EGF and bFGF. Number of cells with spheres were recorded after 2 weeks and clonogenic frequency of cells were plotted using ELDA online software (https://bioinf.wehi.edu.au/software/elda/).

### In vitro invasion experiments

PDGCs were dissociated into single cells and resuspended in 0.2% Methyl cellulose (Sigma, Cat# M0512) at a density of 3000 cells/50 μL. 50 μL of the single cell suspension was dispensed per well in a 96-well Clear Round Bottom Ultra-Low Attachment Microplate (Corning, Cat# 7007) followed by 200g × 3 minutes centrifugation at room temperature in a swinging bucket rotor. Then the plate was incubated at 37⁰C in a tissue culture incubator for 72 hours to promote spheroid formation. Plates were prechilled on ice for 15 minutes before adding 50 μL Matrigel followed by 300 x g centrifugation at 4⁰C for 5 minutes. Images were taken every day by EVOS M7000 and invasion area was analyzed using ImageJ.

### WST-8 assays

Cells were enzymatically dissociated into single cells using Accutase and counted using Trypan blue (Thermo Fisher, Cat# T10282) staining and Countess III (Invitrogen). 3000-5000 cells were seeded in each well of 96-well plate. Cells were treated with control vehicle (DMSO, PBS, or water) or pharmacological agents of interest at desired concentrations on day 0. Cell viability was tested using WST-8 (MedChemExpress, Cat# HY-K0301) according to the manufacturer’s protocol. Following a two-hour incubation at 37°C, absorbance at 450 nm was measured with a Synergy H1 Plate Reader (BioTek).

### 5-EU incorporation assay

PDGCs were plated on cover slips (Fisher Scientific, Cat# 50143822) pre-coated with Poly-L-Ornithine and Laminin in 24-well plates. Each well was treated with 1 mM 5-ethynyl uridine (5-EU) dissolved in complete culture medium for 1 hour followed by fixation with 1% PFA for 20 minutes and permeabilization with 0.5% Triton X-100 for 10 minutes. Cells were treated with click reaction master mix (5.0 μM AZDye 647, 0.5 mg/mL CuSO_4_ ·5H_2_O, and 20 mg/mL fresh ascorbic acid dissolved in water) for 30 minutes at room temperature in the dark. Coverslips were washed with PBS-T, then counter-stained with 3 μg/mL Hoechst 333342 dye in PBS for 10 minutes in the dark. Fluorescent images were taken using LSM800 microscope with Aryscan (Zeiss) after mounting coverslips with ProLong Gold antifade reagent.

### Immunohistochemistry staining of human GBM tissues

GBM specimens obtained from patients after informed consent (NYU IRB study 12-01130) were fixed and kept in 4% polyformaldehyde (PFA) at 4°C. Fixed tissues were sent to the Experimental Pathology Research Laboratory at NYU Langone (RRID:SCR_017928) for paraffin sectioning. Immunohistochemistry (IHC) staining was performed with IHC Prep & Detect Kit for Rabbit/Mouse Primary Antibody (Proteintech, Cat# PK10019). In brief, paraffin removal and tissue rehydration were performed by immersing slides into xylene and different concentrations of ethanol sequentially followed by antigen retrieval using Tris-EDTA buffer heated in microwave. Primary antibody (see **Table 4**) was diluted in PBS and incubated at 4°C overnight. Slides were incubated with horseradish peroxidase (HRP) anti-Rabbit/Mouse secondary antibody for 30 minutes at room temperature. Immunohistochemistry signals were developed using chromogen and nuclei were counterstained with counter reagent (Proteintech, Cat# HC009). Slides were rinsed with tap water to remove extra counter reagent before mounting and imaging.

### Electrophysiology

Patch clamping was performed at NYU Langone’s Ion Laboratory (RRID: SCR_021754). Membrane potentials of PDGCs were recorded in the current clamp mode with a 0 pA holding current over a 10 s average in artificial cerebrospinal fluid containing (in mM): 119 NaCl, 2.5 KCl, 1.25 NaH_2_PO_4_, 26 NaHCO_3_, 10 D-Glucose, 1.5 MgSO_4_*7H_2_O and 2.5 CaCl_2_*2H2O. The solution was oxygenated with 95% O_2_/5% CO_2_. The standard internal solution consisted of (in mM): 127 K-gluconate, 8 KCl, 10 Phosphocreatine, 10 HEPES, 4 Mg-ATP and 0.3 Na-GTP. To record Ca^2+^ fluorescence (GCaMP6s) signal and resting membrane potential simultaneously in the same cell, individual cells expressing the tdTomato-2A-GCaMP6s construct were identified using a fluorescent microscope to detect red fluorescence from constitutively expressed tdTomato. Patched cells were immersed in DMEM/F12 culture medium used for Ca^2+^ imaging and all the other settings are the same as recording resting membrane potentials. Line graphs of membrane potential and GCaMP6s signals were plotted using R.

### Implantation of GBM cells in the mouse brain

All the animal and surgical procedures were performed according to Institutional Animal Care and Use Committee (IACUC) protocol IA16-00208 approved by NYU Grossman School of Medicine. Immunodeficient NSG (NOD.Cg-*Prkdc^scid^ Il2rg^tm1Wjl^*/SzJ) mice (6–8 weeks of age) were housed within NYU Grossman School of Medicine animal facilities with 12-hour dark and light cycles. Both female and male mice were used in our experiments. PDGCs were dissociated into single cells using Accutase and mixed 1:1 with Matrigel (Corning, Cat# 356237). After mice were injected with buprenorphine and anesthetized with ketamine/xylazine (10 mg/kg and 100 mg/kg, respectively), a vertical incision was made in the scalp and a burr hole was drilled into the skull posterior and to the right of the bregma. Using a Hamilton syringe (Hamilton, Cat# 7653-01) and a stereotaxic frame (Stoelting), the needle was advanced into the brain to a depth of 2.5 mm, followed by 0.5 mm withdrawal, and injection was performed at a rate of 1 µL/minute using an automated injection device (Harvard Apparatus, Cat# 70-4507). Each injection of the cell suspension and Matrigel mixture contained 5.0 × 10^5^ cells in a volume of 4 µL. Skin incisions were sutured after the surgery and mice were euthanized when observed hunched as well as decreased activity.

### Bioluminescent imaging of GBM xenografts

GBM xenografts were monitored with an IVIS Spectrum system (PerkinElmer) at NYU Langone’s Preclinical Imaging Laboratory (RRID:SCR_017937). Tumor bearing mice were weighed and peritoneally injected with 10 μL/g body weight Luciferin substrate solution (D-Luciferin Potassium Salt, LUCK-300, Gold Biotechnology) diluted in DPBS (15 mg/mL). Mice were anesthetized with isoflurane and imaged with Living Image (PerkinElmer) software 10 minutes after luciferin injection using automatic exposure mode. Quantification analysis of total photon flux normalized with exposure (radiance) time was used to estimate the growth rate of GBM xenografts *in vivo*.

### Imaging nuclear Ca^2+^ transients in vivo

To characterize the presence of PDGC nuclear Ca^2+^ transients *in vivo,* we stereotactically injected 5.0 × 10^5^ GCaMP6f.3xNLS-labeled GSC23 and GBML198 cells into the visual cortex (medial–lateral relative to midline (x) −1.50 mm; anterior–posterior relative to bregma (y) −2.20 mm; dorsal–ventral relative to surface of the brain (z) 0.5 mm) of immunodeficient NSG mice. Two to four weeks after PDGC implantation, mice were fitted with cranial widows for in vivo two-photon imaging. Mice were anesthetized using isoflurane (1.5-2.5%) and the scalp removed to expose the skull over visual cortex. A 3 mm craniotomy was made around our previous injection site. A glass cranial window composed of a 5 mm coverslip and a 3 mm coverslip attached with UV optical glue (Norland Optical Adhesive, Norland Inc) was then lowered into the craniotomy, fixed to the skull with dental cement (Metabond, Parkell Inc), and secured with a custom 3D printed headplate. Dental cement was then used to seal the remaining surgical wound. Care was taken to ensure minimal bleeding within the craniotomy for optimal imaging. Animals were given buprenorphine (0.5-1.0mg/kg) 30 min before the end of surgery and meloxicam (2.0-5.0 mg/kg) every 24 hrs for 48 hrs after analgesia. Animals were further monitored for signs of distress. Mice were allowed to recover until ambulatory before being head fixed for *in vivo* Ca^2+^ imaging [77]. Regions of interest (ROIs) were identified using an epifluorescent lamp. ROIs were imaged using a dual resonant galvanometric laser scanning two-photon microscope (Ultima, Bruker). The microscope was coupled to a tunable Ti:Sapphire laser (MaiTai eHP DeepSee, Spectraphysics) at 80 MHz pulse repetition rates and <70 fs pulse width, to excite GCaMP6f at 920 nm. *In vivo* images were acquired at scan speeds of 30 fps, using 512×512 frame size (1.085 μm/pixel resolution) with a resonant scanning galvanometer system mounted on a movable objective Ultima microscope, with an orbital nosepiece coupled to a 16x, 0.8NA, 3 mm water immersion objective (Nikon). The fluorescence signal was detected using high-sensitivity GaAsP photomultiplier tubes (model 7422PA-40 PMTs, Hamamatsu). Recordings were processed using Suite2p [78–80] for motion correction and CellPose [14] for ROI detection. All subsequent Ca^2+^ transient data extraction and analyses were performed as described above for the *in vitro* imaging. Parameters defined differently for the *in vivo* analysis included the following: predefined thresholds of minimum prominence ≥ 0.3, amplitude ≥ 0.2, inter-peak distance ≥ 3 frames, and a physiologically plausible duration of 2-20 seconds optimized by watching live videos to capture all the Ca^2+^ transients.

### Intratumoral delivery of BPTU in vivo

To test if BPTU was effective *in vivo*, we injected 5.0 × 10^5^ luciferase-labeled GSC23 cells into immunodeficient NSG mice as described above. Tumor bearing mice were assigned into control and BPTU treatment groups according to randomization based on tumor volume measured by IVIS two weeks after PDGC injection. Four μg BPTU was dissolved in 10% DMSO + 90% sterilized water to a total volume of 7 μL. BPTU was injected into the same coordinates at which PDGCs were implanted 13 and 21 days prior. Mice were closely monitored every week with IVIS.

### BPTU pharmacokinetics

To evaluate the ability of BPTU to penetrate the blood-brain barrier (BBB), we administered 10 mg/g/day BPTU to NSG mice via oral gavage daily for two weeks. Plasma samples were collected in EDTA-treated tubes and stored at -80⁰C prior to analysis. Brain tissue samples were collected following perfusion with PBS, flash-frozen, and stored at -80⁰C prior to analysis. Samples were kept on dry ice at all times during handling.

### Extraction of BPTU from Mouse Plasma

Prior to extraction, samples were moved from -80 °C storage to wet ice and thawed. Extraction buffer for plasma samples, consisting of 82% methanol (Fisher Scientific) and 512 nM metabolomics amino acid mix standard (Cambridge Isotope Laboratories, Inc.), was prepared and placed on dry ice. Plasma samples were extracted by mixing 25 µL of sample with 975 µL of extraction buffer in 2.0 mL screw cap vials containing ∼100 µL of disruption beads (Research Products International, Mount Prospect, IL). A 7-point standard curve of BPTU (0.1, 0.3, 1, 3, 10, 30, 100 μM) was prepared, extracted, and analyzed as technical duplicates alongside the plasma samples. Each sample was homogenized for 10 cycles on a bead blaster homogenizer (Benchmark Scientific, Edison, NJ). Cycling consisted of a 30 sec homogenization time at 6 m/s followed by a 30 sec pause. Samples were subsequently spun at 21,000 g for 3 min at 4 °C. A fixed volume of each (450 µL) was transferred to a 1.5 mL tube and dried down by speedvac (Thermo Fisher, Waltham, MA). Samples were reconstituted in 50 µL of LC/MS grade water. Samples were sonicated for 2 mins, then spun at 21,000 g for 3 min at 4°C. 20 µL were transferred to LC vials containing glass inserts for analysis. The remaining sample was placed in -80°C for long term storage.

### Extraction of BPTU from Mouse Brain

Prior to extraction, samples were moved from -80 °C storage to wet ice and thawed. Extraction buffer for brain samples, consisting of 80% methanol (Fisher Scientific) and 500 nM metabolomics amino acid mix standard (Cambridge Isotope Laboratories, Inc.), was prepared and placed on dry ice. Brain samples were extracted using a ration of 10mg/mL (tissue to extraction buffer) in 2.0 mL screw cap vials containing ∼100 µL of disruption beads (Research Products International, Mount Prospect, IL). A matrix-controlled 5-point standard curve of BPTU (3, 10, 30, 100, 300 nM) was prepared in null brain then extracted and analyzed as technical duplicates alongside the brain samples. Each sample was homogenized for 10 cycles on a bead blaster homogenizer (Benchmark Scientific, Edison, NJ). Cycling consisted of a 30 sec homogenization time at 6 m/s followed by a 30 sec pause. Samples were subsequently spun at 21,000 g for 3 min at 4 °C. A fixed volume of each (450 µL) was transferred to a 1.5 mL tube and dried down by speedvac (Thermo Fisher, Waltham, MA). Samples were reconstituted in 50 µL of LC/MS grade Ethanol. Samples were sonicated for 2 mins, then spun at 21,000 g for 3 min at 4°C. 20 µL were transferred to LC vials containing glass inserts for analysis. The remaining sample was placed in -80°C for long term storage.

### LC-MS/MS method for BPTU quantification

Samples were subjected to an LCMS analysis to detect and quantify BPTU. The LCMS parameters were adapted from a previously described method [81]. The LC column was a WatersTM BEH-Phenyl (2.1 x150 mm, 1.7 μm) coupled to a Dionex Ultimate 3000TM system and the column oven temperature was 25oC for the gradient elution. A flow rate of 200 μL/min was used with the following buffers: A) 0.1% formic acid in water, and B) 0.1% formic acid in acetonitrile. The gradient profile was as follows; 0-35% B (0-10 min), 35-75% B (10-15 min), 75-99% B (15-15.25 min), 99-99% B (15.25-16.5 min), 99-0% B (16.5-16.75 min), 0-0% B (16.75-20 min). Injection volume was 2 μL for all analyses (20 min total run time per injection).

MS analyses were carried out by coupling the LC system to a Thermo Q Exactive HF™ mass spectrometer operating in heated electrospray ionization mode (HESI). Method duration was 20 min with a polarity switching data-dependent Top 5 method for both positive and negative modes. Spray voltage for both positive and negative modes was 3.5 kV and capillary temperature was 320°C with a sheath gas rate of 35, aux gas of 10, and max spray current of 100 μA. The full MS scan for both polarities utilized 120,000 resolution with an AGC target of 3e6 and a maximum IT of 100 ms, and the scan range was from 95-1000 m/z. Tandem MS spectra for both positive and negative mode used a resolution of 15,000, AGC target of 1e5, maximum IT of 50 ms, isolation window of 0.4 m/z, isolation offset of 0.1 m/z, fixed first mass of 50 m/z, and 3-way multiplexed normalized collision energies (nCE) of 10, 35, 80. The minimum AGC target was 1e4 with an intensity threshold of 2e5. All data were acquired in profile mode.

### Absolute quantification of BPTU

The resulting ThermoTM RAW files were converted to SQLite format using an in-house python script to enable downstream peak detection and quantification. The centroided data were searched using an in-house python script to quantify peak heights (https://github.com/NYUMetabolomics/plz) and the BPTU m/z and retention time (RT) were both confirmed using an external authentic standard at 10 μM during the same batch. BPTU peaks were extracted based on the theoretical m/z of the expected ion type as characterized with the authentic standard (e.g., [M+H]+), with a ±5 part-per-million (ppm) tolerance, and a ± 7.5 second peak apex retention time tolerance within an initial retention time search window of ± 0.5 min across the study samples. BPTU peak detection was determined based on a signal to noise ratio (S/N) of 3X compared to blank controls, with a floor of 10,000 (arbitrary units). Absolute quantification was calculated by plotting the detected peak heights from the known concentration BPTU standard curve points using Graphpad Prism 9 and interpolating the detected sample unknown concentration BPTU peak heights against the linear curve model.

### Single cell RNA sequencing analysis

Single-cell RNA sequencing (scRNA-seq) data from human gliomas were obtained from a previously published study[82] and accessed through the Broad Institute Single Cell Portal. Data preprocessing and analysis were performed using Scanpy (v1.12). Cells with low gene counts, elevated mitochondrial transcript content, or suspected doublets were excluded during quality control. Gene expression counts were normalized on a per cell basis and log-transformed for downstream analyses. Highly variable genes (HVGs; n = 3000) were identified and used for dimensionality reduction and neighborhood graph construction; however, visualization and marker-based annotation analyses were performed using the full gene expression matrix to retain biologically relevant genes not captured within the HVG subset, including immune and lineage-associated markers. Principal component analysis (PCA) was performed for dimensionality reduction, followed by construction of a k-nearest neighbor graph in principal component space to model transcriptional similarity between cells. Unsupervised clustering was performed using the Leiden algorithm, and clusters were visualized using Uniform Manifold Approximation and Projection (UMAP). Cell type annotation was performed at the Leiden cluster level using canonical lineage marker genes visualized across UMAP plots and cluster-level expression patterns. Cluster identities were assigned manually based on established marker expression profiles, including markers for glioma cells (e.g., *EGFR*, *SOX2*, *PDGFRA*, *OLIG2*), myeloid cells (e.g., *PTPRC*, *P2RY12*, *LYZ*), T cells (e.g., *PTPRC*, *CD3D*, *CD3E*), oligodendrocyte-lineage cells (e.g., *MBP*, *PLP1*, *MOG*, *SOX10*), B cells (*PTPRC*, *MS4A1*, *CD19*), endothelial cells (e.g., *PECAM1*, *CLDN5*), and pericytes (e.g., *PDGFRB*, *ACTA2*). Clusters lacking clear lineage-defining markers were classified as “Other.” Differential gene expression analysis between Leiden clusters was performed using the Wilcoxon rank-sum test to identify genes significantly enriched within each cluster relative to all remaining cells. These cluster-enriched genes were used to identify cluster-specific transcriptional signatures.

### Bulk RNA-seq analysis

RNA was extracted using Monarch® Total RNA Miniprep Kit (NEB, Cat# T2010S). RNA was sent to the NYU Langone Genome Technology Center (RRID: SCR_017929) for automated stranded library preparation with poly-A selection and sequencing after RNA integrity was measured. The FASTQC tool was used to verify data quality before downstream analysis. FASTQ files were aligned to GRCh38 using STAR aligner to get row count values using BigPurple HPC Cluster [83]. DeSeq2 [84] package was used for differential analysis, and ClusterProfiler [85] was used for gene set enrichment analysis. Data visualization was performed using ggplot2 package.

### ChIP-seq analysis

ChIP-seq samples were prepared according to the published protocols with modifications to fit our own experiments [86]. In brief, PDGCs were dissociated into single cells and fixed with methanol-free formaldehyde (CST, Cat# 12606) to a final concentration of 1% in the culture medium and shaken to mix at room temperature for 10 minutes. Extra formaldehyde was quenched by adding 1:10 volume of cold 1.25 M glycine (Bio-Rad, Cat# 161-0718) and samples were rocked at room temperature for 5 minutes. Fixed cells were lysed and sheared 30 cycles (30s-on and 30s-off) with Diagenode Bioruptor Sonication System (Diagenode) at 4°C. ChIP grade antibody and magnetic beads (Sigma, Cat# 16-661) were put into ChIP lysates and incubated overnight at 4°C. Antibody-chromatin complexes were eluted, reverse crosslinked, and treated with RNase A (Thermo Fisher, Cat# EN0531), followed by proteinase K (NEB, Cat# P8107S). Input and ChIP DNA were purified using ChIP DNA Clean & Concentrator kit (Zymo research, Cat# D5201). DNA was sent to the NYU Genome Technology Center for library preparation using NEBNext Ultra II (NEB) and sequencing by Novaseq X+ (Illumina). Bowtie2 was used to align FASTQ files to GRCh38 [87]. MACS2 [88] and Deeptools were used for peak calling and data visualization [89].

### DamID sequencing analysis

GBML198 cells were transduced with pLenti-SFFV-mScarI-Dam or pLenti-SFFV-mScarI-Dam.CaMK4 lentivirus. Four days later, mScarlet expression was confirmed with fluorescent microscope before collecting cells for sequencing. Targeted DamID for CamK4 was performed according to published protocol [54, 90]. In short, genomic DNA was isolated and purified using QIAamp DNA Micro Kit (Qiagen, Cat# 56304), then digested with DpnI and DpnII to isolate methylated DNA. DNA fragments were then amplified by PCR and sonicated to generate fragment sizes appropriate for sequencing. The library was sequenced using the Illumina NovaSeq 6000 in the NYU Langone Genome Technology Center. FASTQ files were analyzed according to published pipelines [54].

### Statistical analysis

All the experiments were performed at least 3 independent biological replicates unless otherwise specified. Statistical tests were performed using GraphPad Prism (Version 10.1.2). Normality or log-normality were tested for numerical data. Parametric tests including *t*-test, one-way ANOVA with *post hoc* tests or 2-way ANOVA with *post hoc* tests were used to get statistical results for normally distributed data. Nonparametric tests including Mann-Whitney and Kruskal-Wallis test with multiple comparisons were used for non-normally distributed data. Log-rank test was used for survival analysis. Statistical significance was set at *P* < 0.05. Throughout the manuscript, the following notations were used: * *P* < 0.05, ** *P* < 0.01, *** *P* < 0.001, **** *P* < 0.0001.

## Data, code, and materials availability

RNA-Seq data and ChIP-Seq data were deposited in the Gene Expression Omnibus (GEO) database. Codes used for RNA-Seq analysis, ChIP-Seq analysis, and calcium imaging analysis were deposited in GitHub (https://github.com/Shuai-NY).

## Acknowledgements

We thank Stacy Mahiga, Nata Kakabadze, and Devin Bready for technical assistance. We also thank the Genome Technology Center (RRID: SCR_017929), the Microscopy Laboratory (RRID: SCR_017934), Experimental Pathology Research Laboratory (RRID:SCR_017928), Ion Laboratory (RRID: SCR_021754), Small Instrument Fleet, and the Skirball Mouse Facility at the NYU Grossman School of Medicine. We are grateful to Dr. Hilmar Bading for sharing GCaNP6f.3xNLS, PV.NLS, and CaMBP4-mC plasmids; Drs. Erik Sulman and Jann Sarkaria for sharing the GSC23 and G12 PDGCs; and Dr. Iannis Aifantis and his lab members at NYU Grossman School of Medicine for use of lab equipment. This work was supported by NINDS R21NS126806 (DGP), NINDS R01NS124920 (DGP), the Childhood Brain Tumor Foundation (DGP and JB), NINDS RM1NS132981 (JB), NIA R01AG094086 (JB), Alzheimer’s Association AARGD-NTF-23-1151101 (JB), and the NYU Department of Neurosurgery (DGP).

