## Supplementary figures for "Purinergic signaling promotes gliomagenesis through nuclear calcium transients"

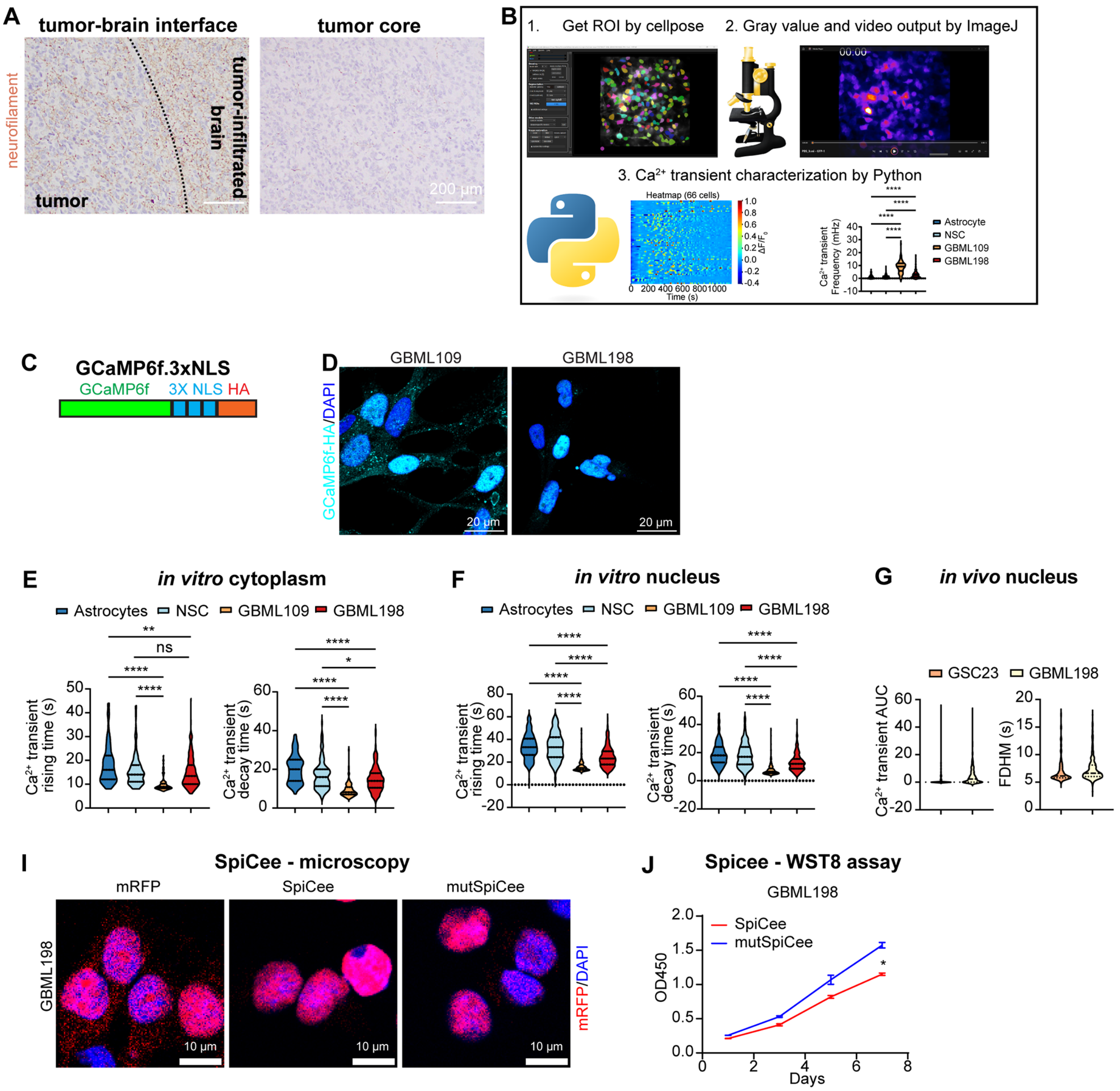


**Supplementary Figure 1: Characterization of Ca$^{2+}$ transients *in vitro* and *in vivo*. (A)** Immunostaining for Neurofilament at the tumor brain interface and tumor core of a GBM operative specimen. **(B)** Schematic diagram of the computational pipeline for analysis of Ca$^{2+}$ transients (ROIs = regions of interest). **(C)** Schematic diagram of nuclear Ca$^{2+}$ sensor GCaMP6f.3xNLS with hemagglutinin (HA) tag. **(D)** Immunofluorescent staining for HA in GBM109 and GBML198 cells transduced with GCaMP6f.3xNLS lentivirus. **(E)** Violin plots showing median and interquartile range of Ca$^{2+}$ transient rising time and decay time in the cytoplasm of immortalized astrocytes (n = 89), NSCs (n = 165), GBML109 (n = 144), and GBML198 (n = 229). Kruskal-Wallis test ($H_{rising}$

= 171.5; $H_{decay}$ = 206.8) with *post hoc* multiple comparisons. **(F)** Violin plots showing median and interquartile range of $Ca^{2+}$ transients rising time and decay time in the nucleus of immortalized astrocytes (n = 263), NSCs (n = 257), GBML109 (n = 233), and GBML198 (n = 296). Kruskal-Wallis test with multiple comparisons. **(G)** Violin plots showing median and interquartile range of $Ca^{2+}$ transient AUC and FDHM in the nucleus of GSC23 ($n_{AUC}$ = 498, $n_{FDHM}$ = 156) and GBML198 ($n_{AUC}$ = 703, $n_{FDHM}$ = 448) cells *in vivo* (n = 4 mice per xenograft model). **(H)** Representative fluorescent images of GBML198 cells transduced with mRFP, SpiCee, or mutSpiCee. **(I)** Line graph depicting OD450 values (y-axis) over time (x-axis) as measured by WST-8 assay in GBML198 cells transduced with SpiCee or mutSpiCee (two-way RM ANOVA with Šídák's multiple comparisons test, n = 3/group, $F_{(3, 12)}$ = 13.06). * $p < 0.05$, ** $p < 0.01$, **** $p < 0.0001$, ns = not significant.

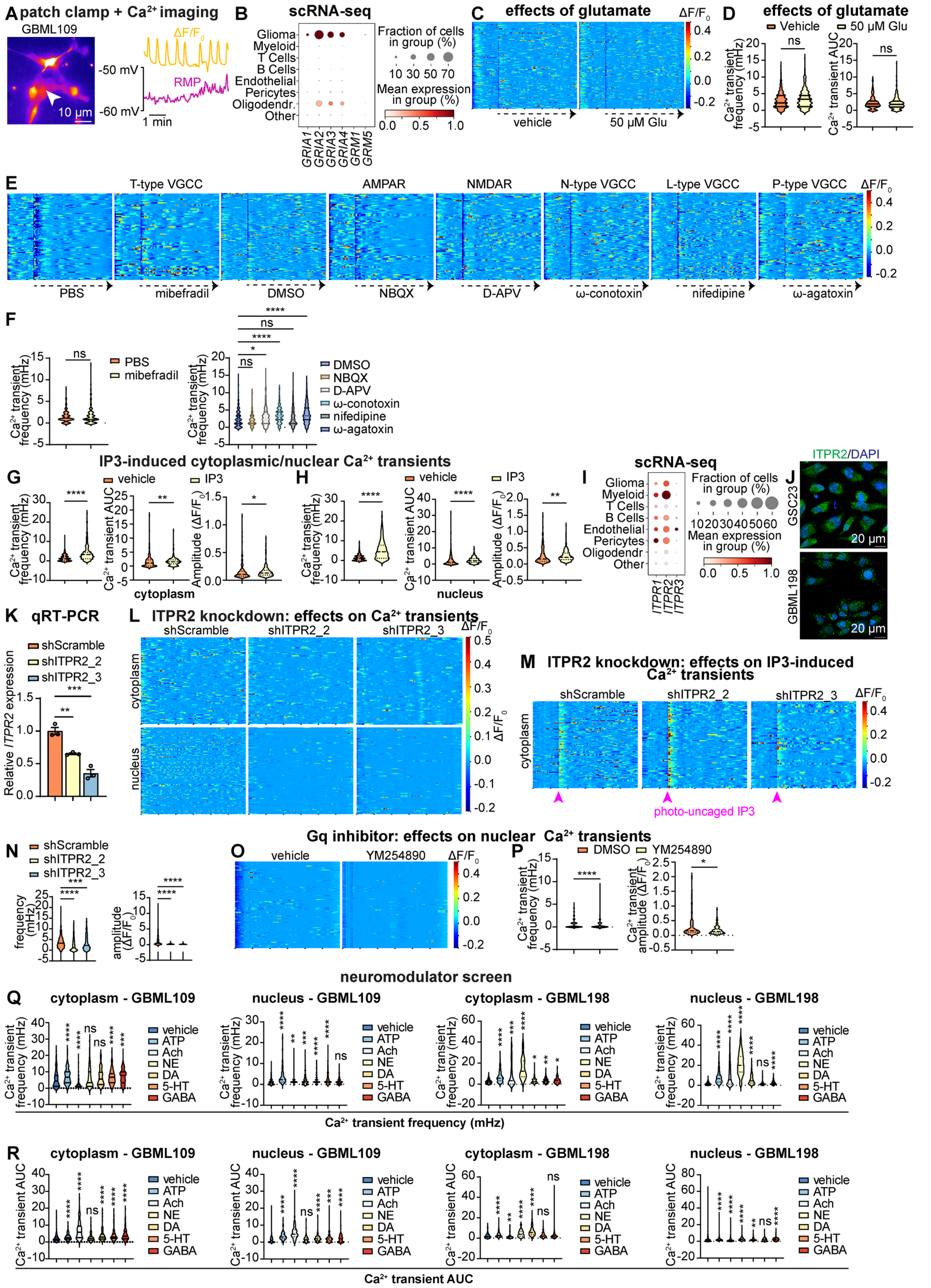


**Supplementary Figure 2: Intracellular $IP_3$ generated by Gq-coupled GPCRs produces $Ca^{2+}$**

**transients. (A)** Representative example of simultaneous $Ca^{2+}$ imaging with GCaMP6s and whole-cell patch-clamp recording of RMP in a GBML109 cell. **(B)** Dot plot showing the expression of ionotropic and metabotropic glutamate receptors in a sc RNA-Seq dataset of GBM specimens (GSE182109). **(C,D)** Response of GCaMP6s-expressing GBML198 cells to 50 µM glutamic acid represented as (**C**) 20-minute heatmaps ($n_{vehicle}$ = 126, $n_{glutamate.}$ = 145) of $\Delta F/F_0$ responses and (**D**) violin plots showing median and interquartile range of $Ca^{2+}$ transient frequency and AUC (Mann Whitney U test; $n_{vehicle}$ = 360, $n_{glutamate.}$ = 424 from 3 biological replicates). **(E)** Heatmaps showing 20-minute recordings of $\Delta F/F_0$ responses to $Ca^{2+}$ channel antagonists in GBML198 cells transduced with GCaMP6s lentivirus ($n_{PBS}$ = 84, $n_{mibefradil}$ = 69, $n_{DMSO}$ = 136, $n_{NBQX}$ = 76, $n_{D\text{-}APV}$ = 86, $n_{\omega\text{-}conotoxin}$ = 120, $n_{nifedipine}$ = 110, $n_{\omega\text{-}agatoxin}$ = 98). The target of each antagonist is indicated above its respective heatmap. **(F)** Violin plots showing median and interquartile range of $Ca^{2+}$ transient frequency responses to $Ca^{2+}$ channel antagonists in GBML198 cells transduced with GCaMP6s lentivirus (left: Mann-Whitney U test, $n_{PBS}$ = 330, $n_{mibefradil}$ = 303; right: Kruskal-Wallis with Dunnett's multiple comparisons test, $n_{DMSO}$ = 368, $n_{NBQX}$ = 194, $n_{D\text{-}APV}$ = 351, $n_{\omega\text{-}conotoxin}$ = 210, $n_{nifedipine}$ = 405, $n_{\omega\text{-}agatoxin}$ = 334)**. (G,H)** Violin plots showing median and interquartile range of $Ca^{2+}$ transient frequency (cytoplasm: $n_{vehicle}$ = 230, $_{nIP3}$ = 249; nucleus: $n_{vehicle}$ = 270, $n_{IP3}$ = 352), AUC (cytoplasm: $n_{vehicle}$ = 230, $n_{IP3}$ = 249; nucleus: $n_{vehicle}$ = 270, $n_{IP3}$ = 352), and amplitude (cytoplasm: $n_{vehicle}$ = 152, $n_{IP3}$ = 222; nucleus: $n_{vehicle}$ = 177, $n_{IP3}$ = 313) in the cytoplasm **(G)** and nucleus **(H)** of GBML198 cells in response to vehicle control or photo-uncaging of 1 µM $IP_3$ (Mann-Whitney U test). **(I)** Dot plot showing the expression of $IP_3$Rs (*ITPR*s) in a sc RNA-Seq dataset of GBM specimens (GSE182109). **(J)** Immunofluorescent staining for ITPR2 in GSC23 and GBML198 cells *in vitro*. **(K)** Bar graph depicting mRNA expression of *ITPR2* relative to *ACTB* as measured by qRT-PCR in GBML198 cells (one-way ANOVA with Dunnett's multiple comparisons test, $F_{(2, 6)}$ = 48.88 across three biological replicates). **(L)** Heatmaps showing 20-minute recordings of $\Delta F/F_0$ in the cytoplasm ($n_{shScramble}$ = 89, $n_{shITPR2\_2}$ = 77, $n_{shITPR2\_3}$ = 70) and nucleus ($n_{shScramble}$ = 141, $n_{shITPR2\_2}$ = 121, $n_{shITPR2\_3}$ = 149) of GBML198 cells transduced with shScramble and two non-overlapping shRNA targeting ITPR2. **(M,N)** Response of GBML198 cells transduced with shScramble and two shRNA targeting ITPR2 and treated with 1 µM photo-uncaged $IP_3$ represented as **(M)** heatmaps ($n_{shScramble}$ = 114, $n_{shITPR2\_2}$ = 74, $n_{shITPR2\_3}$ = 59) of cytoplasmic $Ca^{2+}$ transient $\Delta F/F_0$ in 20 minutes and **(N)** violin plots showing median and interquartile range of $Ca^{2+}$ transient frequency ($n_{shScramble}$ = 303, $n_{shITPR2\_2}$ = 197, $n_{shITPR2\_3}$ = 182) and amplitude ($n_{shScramble}$ = 282, $n_{shITPR2\_2}$ = 112, $n_{shITPR2\_3}$ = 144) (Kruskal-Wallis test (H = 51.53) with *post hoc* multiple comparisons across three separate experiments). **(O,P)** Response of GBML198 cells transduced with GCaMP6f.3xNLS to 2 μM YM254890 represented as **(O)** 20-minute heatmaps ($n_{vehicle}$ = 126, $n_{YM254890}$ = 116) of nuclear $Ca^{2+}$ transient $\Delta F/F_0$ and **(P)** violin plots showing median and interquartile range of $Ca^{2+}$ transient frequency ($n_{vehicle}$ = 352, $n_{YM254890}$ = 477) and amplitude ($n_{vehicle}$ = 135, $n_{YM254890}$ = 117) (Mann-Whitney U test across three separate experiments). **(Q,R)** Violin plots showing median and interquartile range of $Ca^{2+}$ transient frequency **(Q)** and AUC **(R)** in the cytoplasm and nucleus of GBML109 and GBML198 cells treated with vehicle, 100 μM ATP, 1 mM acetylcholine (Ach), 100 μM norepinephrine (NE), 100 μM dopamine (DA), 100 μM serotonin (5-HT), or 100 μM GABA across 3 different recordings (Kruskal-Wallis test (cytoplasm of GBML109: $H_{frequency}$ = 90.71, $H_{AUC}$ = 238.6; nucleus of GBML109: $H_{frequency}$ = 114.6, $H_{AUC}$ = 255.2; cytoplasm of GBML198: $H_{frequency}$ = 338.2, $H_{AUC}$ = 234.3; nucleus of GBML198: $H_{frequency}$ = 1683, $H_{AUC}$ = 293.7) with *post hoc* multiple comparisons; cytoplasm of GBML109: $n_{vehicle}$ = 205, $n_{ATP}$ = 274, $n_{Ach}$ = 168, $n_{NE}$ = 224, $n_{DA}$ = 214, $n_{5\text{-}HT}$ = 200, and $n_{GABA}$ = 148; nucleus of GBML109: $n_{vehicle}$ = 265, $n_{ATP}$ = 75, $n_{Ach}$ = 137, $n_{NE}$ = 253, $n_{DA}$ = 228, $n_{5\text{-}HT}$ = 199, and $n_{GABA}$ = 284; cytoplasm of GBML198: $n_{vehicle}$ = 172, $n_{ATP}$ = 278, $n_{Ach}$ = 229, $n_{NE}$ = 120, $n_{DA}$ = 206, $n_{5\text{-}HT}$ = 203, and $n_{GABA}$ = 188; nucleus of GBML198: $n_{vehicle}$ = 647, $n_{ATP}$ = 487, $n_{Ach}$ = 881, $n_{NE}$ = 629, $n_{DA}$ = 455, $n_{5\text{-}HT}$ = 441, and $n_{GABA}$ = 521). * $p < 0.05$, ** $p < 0.01$, *** $p < 0.001$, **** $p < 0.0001$, ns = not significant.

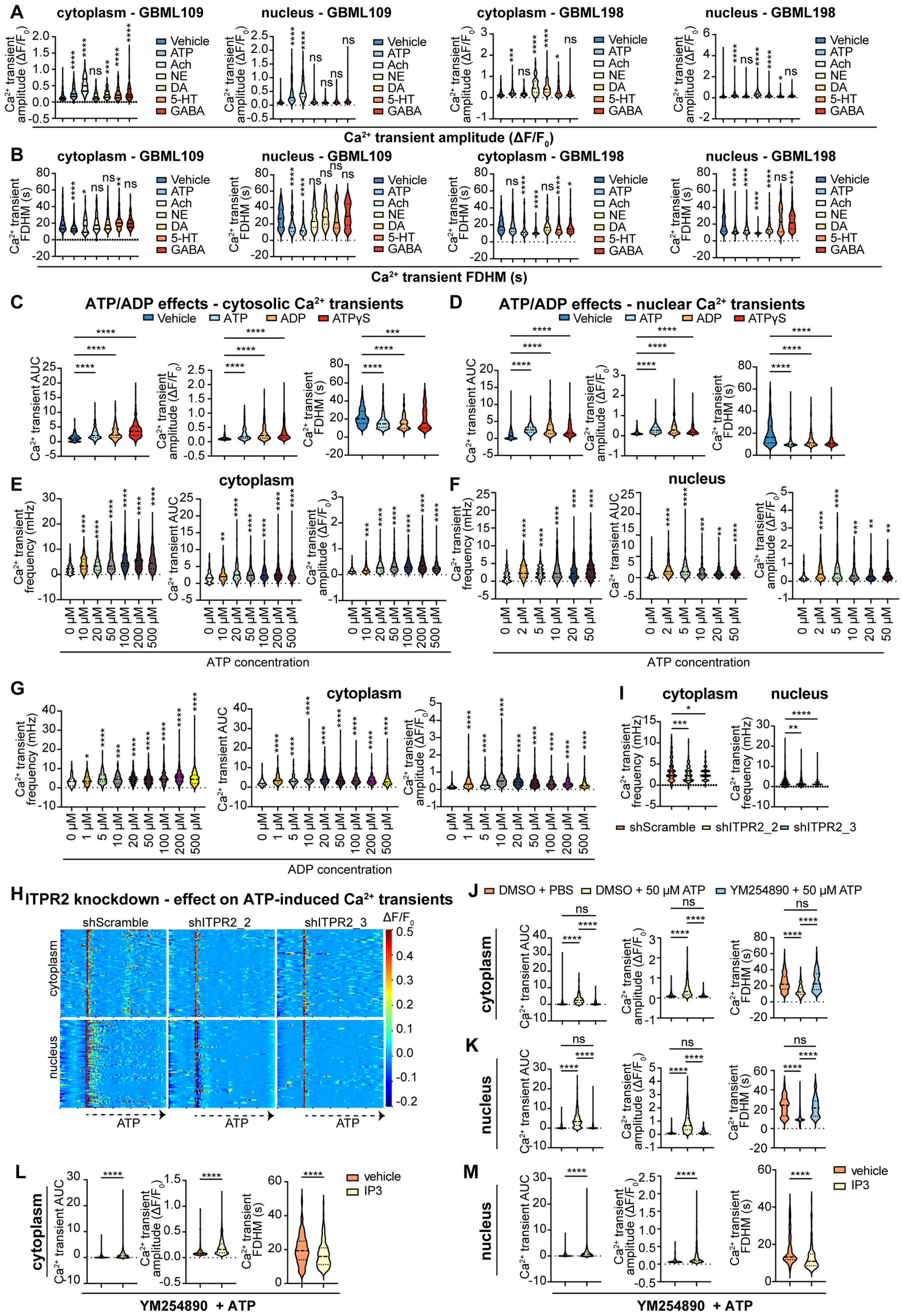


**Supplementary Figure 3: ATP/ADP induces cytoplasmic and nuclear Ca$^{2+}$ transients. (A,B)**

Violin plots showing median and interquartile range of $Ca^{2+}$ transient amplitude **(A)** and FDHM **(B)** in the cytoplasm and nucleus of GBML109 and GBML198 cells treated with vehicle, 100 μM ATP, 1 mM acetylcholine (Ach), 100 μM norepinephrine (NE), 100 μM dopamine (DA), 100 μM serotonin (5-HT), or 100 μM GABA across 3 different recordings (Kruskal-Wallis test (cytoplasm of GBML109: $H_{amplitude}$ = 402.7, $H_{FDHM}$ = 162.0; nucleus of GBML109: $H_{amplitude}$ = 325.4, $H_{FDHM}$ = 205.3; cytoplasm of GBML198: $H_{amplitude}$ = 222.3, $H_{FDHM}$ = 146.0; nucleus of GBML198: $H_{amplitude}$ = 559.7, $H_{FDHM}$ = 1321) with *post hoc* multiple comparisons; cytoplasm of GBML109: $n_{vehicle}$ = 166, $n_{ATP}$ = 262, $n_{Ach}$ = 165, $n_{NE}$ = 182, $n_{DA}$ = 147, $n_{5\text{-}HT}$ = 154, and $n_{GABA}$ = 120; nucleus of GBML109: $n_{vehicle}$ = 149, $n_{ATP}$ = 72, $n_{Ach}$ = 130, $n_{NE}$ = 169, $n_{DA}$ = 193, $n_{5\text{-}HT}$ = 161, and $n_{GABA}$ = 195; cytoplasm of GBML198: $n_{vehicle}$ = 140, $n_{ATP}$ = 272, $n_{Ach}$ = 226, $n_{NE}$ = 117, $n_{DA}$ = 201, $n_{5\text{-}HT}$ = 189, and $n_{GABA}$ = 168; nucleus of GBML198: $n_{vehicle}$ = 490, $n_{ATP}$ = 477, $n_{Ach}$ = 702, $n_{NE}$ = 621, $n_{DA}$ = 434, $n_{5\text{-}HT}$ = 344, and $n_{GABA}$ = 466). **(C,D)** Violin plots showing median and interquartile range of $Ca^{2+}$ transient AUC (cytoplasm: $n_{vehicle}$ = 177, $n_{ATP}$ = 202, $n_{ADP}$ = 168, $n_{ATP\gamma S}$ = 248; nucleus: $n_{vehicle}$ = 282, $n_{ATP}$ = 509, $n_{ADP}$ = 290, $n_{ATP\gamma S}$ = 343), amplitude and FDHM (cytoplasm: $n_{vehicle}$ = 141, $n_{ATP}$ = 195, $n_{ADP}$ = 160, $n_{ATP\gamma S}$ = 244; nucleus: $n_{vehicle}$ = 158, $n_{ATP}$ = 505, $n_{ADP}$ = 288, $n_{ATP\gamma S}$ = 331) in the cytoplasm **(C)** and nucleus (**D**) of GBML198 cells treated with vehicle, 100 μM ATP, 100 μM ADP, or 100 μM ATPγS across 3 different recordings (Kruskal-Wallis test (cytoplasm of GBML198: $H_{AUC}$ = 171.1, $H_{amplitude}$ = 138.3, $H_{FDHM}$ = 47.3; nucleus of GBML198: $H_{AUC}$ = 295.3, $H_{amplitude}$ = 165.8, $H_{FDHM}$ = 96.28) with *post hoc* multiple comparisons). **(E,F)** Violin plots showing median and interquartile range of $Ca^{2+}$ transient frequency and AUC (cytoplasm: $n_{0\mu M}$ = 276, $n_{10\mu M}$ = 277, $n_{20\mu M}$ = 279, $n_{50\mu M}$ = 217, $n_{100\mu M}$ = 266, $n_{200\mu M}$ = 425, $n_{500\mu M}$ = 332; nucleus: $n_{0\mu M}$ = 151, $n_{2\mu M}$ = 160, $n_{5\mu M}$ = 194, $n_{10\mu M}$ = 168, $n_{20\mu M}$ = 284, $n_{50\mu M}$ = 298), and amplitude (cytoplasm: $n_{0\mu M}$ = 226, $n_{10\mu M}$ = 258, $n_{20\mu M}$ = 264, $n_{50\mu M}$ = 214, $n_{100\mu M}$ = 259, $n_{200\mu M}$ = 418, $n_{500\mu M}$ = 318; nucleus: $n_{0\mu M}$ = 96, $n_{2\mu M}$ = 147, $n_{5\mu M}$ = 177, $n_{10\mu M}$ = 146, $n_{20\mu M}$ = 251, $n_{50\mu M}$ = 271) in the cytoplasm **(E)** and nucleus **(F)** of GBML198 cells treated with a range of 0-500 μM ATP across three separate experiments (Kruskal-Wallis test (cytoplasm of GBML198: $H_{frequency}$ = 237.8, $H_{AUC}$ = 94.53, $H_{amplitude}$ = 338.1; nucleus of GBML198: $H_{frequency}$ = 77.06, $H_{AUC}$ = 88.07, $H_{amplitude}$ = 54.37) with *post hoc* multiple comparisons). **(G)** Violin plots showing median and interquartile range of $Ca^{2+}$ transient frequency and AUC ($n_{0\mu M}$ = 196, $n_{1\mu M}$ = 163, $n_{5\mu M}$ = 187, $n_{10\mu M}$ = 166, $n_{20\mu M}$ = 244, $n_{50\mu M}$ = 307, $n_{100\mu M}$ = 206, $n_{200\mu M}$ = 249, $n_{500\mu M}$ = 204), and amplitude ($n_{0\mu M}$ = 181, $n_{1\mu M}$ = 161, $n_{5\mu M}$ = 182, $n_{10\mu M}$ = 164, $n_{20\mu M}$ = 238, $n_{50\mu M}$ = 294, $n_{100\mu M}$ = 200, $n_{200\mu M}$ = 246, $n_{500\mu M}$ = 198) in the cytoplasm of GBML198 cells treated with a range of 0-500 μM ADP across three separate experiments (Kruskal-Wallis test ($H_{frequency}$ = 96.09, $H_{AUC}$ = 178.7) with *post hoc* multiple comparisons). **(H,I)** Response of GBML198 cells transduced with shScramble and two non-overlapping shRNA targeting ITPR2 and treated with 100 μM ATP represented as **(H)** heatmaps of cytoplasmic ($n_{shScramble}$ = 105, $n_{shITPR2\_2}$ = 87, $n_{shITPR2\_3}$ = 80) and nuclear ($n_{shScramble}$ = 76, $n_{shITPR2\_2}$ = 63, $n_{shITPR2\_3}$ = 66) $Ca^{2+}$ transient $\Delta F/F_0$ in 20 minutes and **(I)** violin plots showing median and interquartile range of $Ca^{2+}$ transient frequency (cytoplasm: $n_{shScramble}$ = 277, $n_{shITPR2\_2}$ = 188, $n_{shITPR2\_3}$ = 213; nucleus: $n_{shScramble}$ = 524, $n_{shITPR2\_2}$ = 423, $n_{shITPR2\_3}$ = 418) (Kruskal-Wallis test (cytoplasm: $H_{frequency}$ = 14.79; nucleus: $H_{frequency}$ = 34.91) with *post hoc* multiple comparisons across three separate experiments). **(J,K)** Violin plots showing median and interquartile range of $Ca^{2+}$ transient AUC (cytoplasm: $n_{DMSO + PBS}$ = 353, $n_{DMSO + ATP}$ = 306, $n_{YM254890 + ATP}$ = 292; nucleus: $n_{DMSO + PBS}$ = 400, $n_{DMSO + ATP}$ = 490, $n_{YM254890 + ATP}$ = 412) amplitude and FDHM (cytoplasm: $n_{DMSO + PBS}$ = 282, $n_{DMSO + ATP}$ = 112, $n_{YM254890 + ATP}$ = 144; nucleus: $n_{DMSO + PBS}$ = 137, $n_{DMSO + ATP}$ = 403, $n_{YM254890 + ATP}$ = 146) in the cytoplasm **(J)** and nucleus **(K)** of GBML198 cells pre-incubated with 2 μM $G\alpha_q$ antagonist YM254890 or vehicle and treated with 50 μM ATP (Kruskal-Wallis test (cytoplasm: $H_{AUC}$ = 135.9, $H_{amplitude}$ = 177.1, $H_{FDHM}$ = 145.8; nucleus: $H_{AUC}$ = 399.3, $H_{amplitude}$ = 295.3, $H_{FDHM}$ = 296.7) with *post hoc* multiple comparisons across three separate experiments). **(L,M)** Violin plots showing median and interquartile range of $Ca^{2+}$ transient AUC (cytoplasm: $n_{vehicle}$ = 324, $n_{IP3}$ = 363; nucleus: $n_{vehicle}$ = 397, $n_{IP3}$ =432), amplitude (cytoplasm: $n_{vehicle}$ = 189, $n_{IP3}$ = 343; nucleus: $n_{vehicle}$ = 170, $n_{IP3}$ = 307) and FDHM (cytoplasm: $n_{vehicle}$ = 189, $n_{IP3}$ = 343; nucleus: $n_{vehicle}$ = 170, $n_{IP3}$ = 307) in the cytoplasm **(L)** and nucleus **(M)** of GBML198 cells pre-incubated with 2 μM YM254890 and treated initially with 50 μM ATP and subsequently with 1 μM photo-uncaged $IP_3$ or vehicle control (DMSO) (Mann-Whitney U test across three separate experiments). * $p < 0.05$, ** $p < 0.01$, *** $p < 0.001$, **** $p < 0.0001$, ns = not significant.

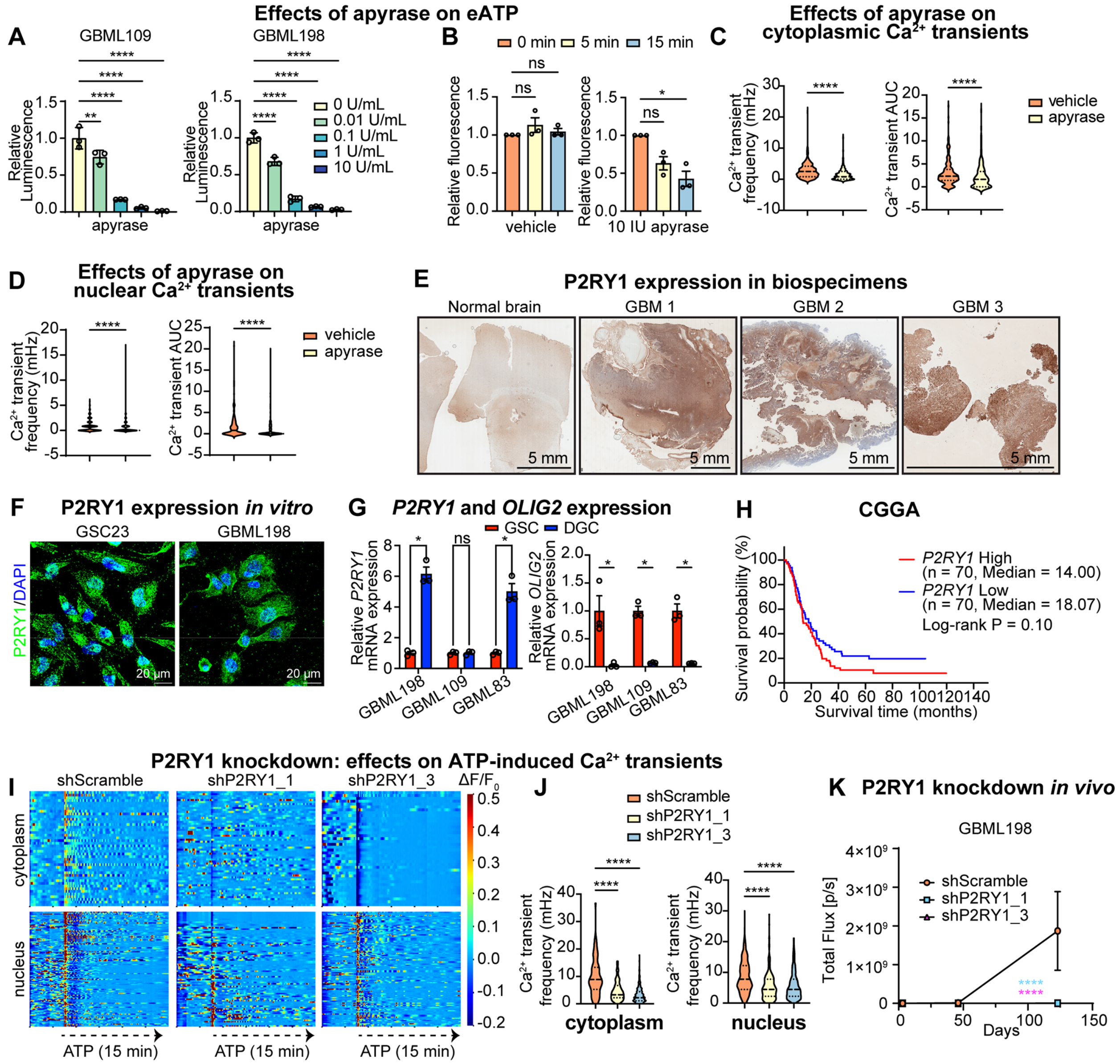


**Supplementary Figure 4: Knockdown of P2RY1 attenuates $Ca^{2+}$ transients and GBM growth.** **(A)** Bar plot depicting relative luminescence measured on eATP assay in response to doses of apyrase ranging from 0-10 IU/mL in GBML109 and GBML198 cells (one way ANOVA with Dunnett's multiple comparisons tests; left: $F_{(4, 10)}$ = 99.79; right: $F_{(4, 10)}$ = 298.7). **(B)** Bar plot depicting relative fluorescence, as measured by confocal microscopy, in GBML198 cells transduced with eATP sensor GRAB 0-15 minutes following treatment with vehicle control or 10 IU/mL apyrase (Kruskal-Wallis test ($H_{vehicle}$ = 3.954; $H_{apyrase}$ = 6.713) with multiple comparisons over three separate experiments). **(C,D)** Violin plots showing median and interquartile range of $Ca^{2+}$ transient frequency and AUC in the cytoplasm **(C)** and nucleus **(D)** of GBML198 cells treated with vehicle control or 10 IU/mL apyrase (Mann-Whitney U test across three separate experiments; cytoplasm: $n_{vehicle}$ = 256, $n_{apyrase}$ = 217; nucleus: $n_{vehicle}$ = 230, $n_{apyrase}$ = 265). **(E)** Immunohistochemistry staining for P2RY1 in three GBM specimens. **(F)** Immunofluorescent staining for P2RY1 in GSC23 and GBML198 cells in vitro. **(G)** Bar graph depicting *P2RY1* mRNA expression relative to *ACTB* in serum-differentiated GBM cells (DGCs) relative to GSCs, as measured by qRT-PCR (n = 3 per group, multiple *t*-test). **(H)** Kaplan-Meier survival plot of GBM patients in the *P2RY1*-high (n = 70) and -low (n = 70) groups from the CGGA database (log-rank test). **(I)** Heatmaps showing 20-minute recordings of cytoplasmic (top panel: $n_{shScramble}$ = 43, $n_{shP2RY1\_1}$= 58, $n_{shP2RY1\_2}$= 40) and nuclear (bottom panel: top panel: $n_{shScramble}$ = 116, $n_{shP2RY1\_1}$= 82, $n_{shP2RY1\_2}$= 106) $\Delta F/F_0$ responses to ATP in GBML198 cells transduced with shScramble or one of two non-overlapping shRNA targeting P2RY1. **(J)** Violin plots showing median and interquartile range of $Ca^{2+}$ transient frequency in the cytoplasm and nucleus of GBML198 cells transduced with shScramble, shP2RY1_2, and shP2RY1_3 lentiviruses and treated with 100 µM ATP across three separate recordings (Kruskal-Wallis test (cytoplasm: $H_{frequency}$ = 124.7; nucleus: $H_{frequency}$ = 146.3) with *post hoc* multiple comparisons; cytoplasm: $n_{shScramble}$ = 102, $n_{shP2RY1\_1}$ = 230, $n_{shP2RY1\_3}$ = 289; nucleus: $n_{shScramble}$ = 578, $n_{shP2RY1\_1}$ = 414, $n_{shP2RY1\_3}$ = 427). **(K)** Line graph depicting total flux (y-axis) over time (x-axis) in an *in vivo* bioluminescent imaging experiment of NSG mice implanted with GBML198 cells transduced with shScramble (n = 10), shP2RY1_2 (n = 8), or shP2RY1_3 (n = 9) lentiviruses (two-way ANOVA with Dunnett's multiple comparisons test after $\log_2$ transformation, $F_{(3.552, 42.63)} = 9.477$). * $p < 0.05$, ** $p < 0.01$, *** $p < 0.001$, **** $p < 0.0001$, ns = not significant.

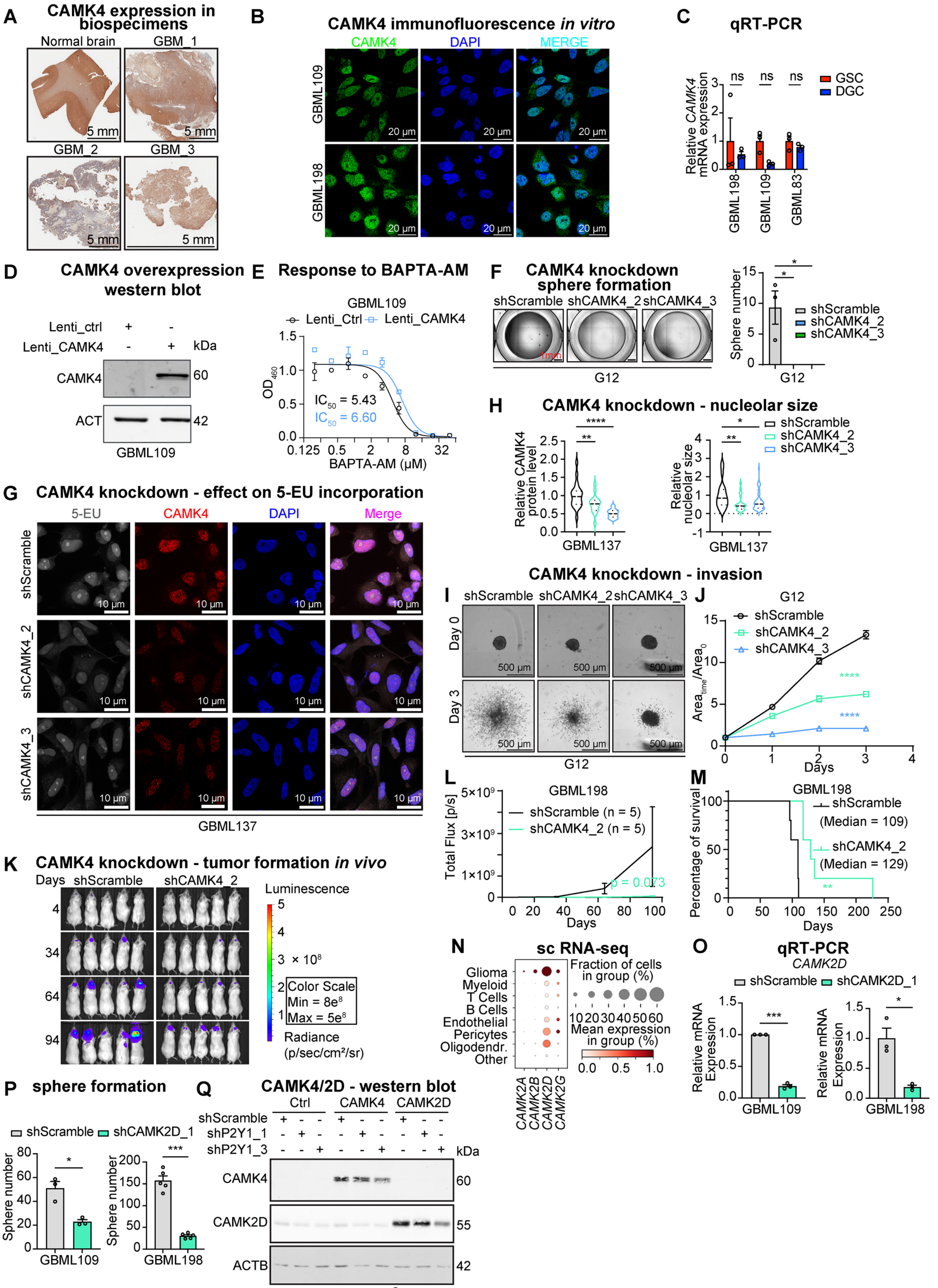


**Supplementary Figure 5: Nuclear $Ca^{2+}$/calmodulin effector CAMK4 is essential for GBM**

**growth. (A)** Immunohistochemistry staining for CAMK4 in three GBM specimens. **(B)** Representative immunofluorescent images of CAMK4 staining in GBML109 and GBML198 cells *in vitro*. **(C)** Bar graph depicting *CAMK4* mRNA expression relative to *ACTB* as measured by qRT-PCR in DGCs relative to GSCs (n = 3/group, multiple *t*-test). **(D)** Immunoblot for CAMK4 and ACTB from GBML109 cells transduced with lenti-ctrl (control) or lenti-CaMK4 lentiviruses. **(E)** Dose-response curve of GBML109 cell viability treated with $Ca^{2+}$ chelator BAPTA-AM (μM) as measured with WST-8 assays. Cells were transduced with lenti-ctrl or lenti-CaMK4 lentiviruses. **(F)** Representative images (left) and bar plot (right) of tumor sphere formation assays in G12 cells transduced with shScramble or two shRNA targeting CAMK4 (one-way ANOVA with Tukey's multiple comparisons test. n = 3/group, $F_{(2, 6)} = 9.673$). **(G)** Representative fluorescent images showing 5-EU (gray), CAMK4 (red), and DAPI (blue) in GBML137 cells transduced with shScramble or one of two shRNA targeting CAMK4. **(H)** Violin plots showing relative CAMK4 protein level (left) and relative nucleolar size (right) measured by 5-EU in GBML137 transduced with shScramble or one of two shRNA targeting CAMK4. Horizontal dash lines indicate the interquartile range, and medians are indicated by a black horizontal line (n = 29-51, one way ANOVA with Dunnett's multiple comparisons tests, left: $F_{(2, 64)} = 26.92$, right: $F_{(2, 100)} = 5.366$). **(I)** Representative images of 3D invasion assays at days 0 and 3 in G12 cells transduced with shScramble or one of two shRNA targeting CAMK4. **(J)** Line graph showing $area_{time}$ /$area_0$ (y-axis) over time (x-axis) of 3D invasion assays in G12 cells shown in (I) (n = 4/group, two-way ANOVA with Šídák's multiple comparisons test, $F_{(6, 36)} = 106.0$). **(K)** Longitudinal bioluminescent images of NSG mice injected with GBML198 cells transduced with shScramble or shCAMK4_2 lentivirus. **(L)** Line graph depicting total flux (y-axis) over time (x-axis) of *in vivo* bioluminescent imaging experiment in mice injected with GBML198 cells transduced with shScramble or shCAMK4_2 (n = 5/group, two-way RM ANOVA with Šídák's multiple comparisons test). **(M)** Kaplan-Meier curve showing survival time of mice xenografted with GBML198 cells transduced with shScramble or shCAMK4_2 (n = 5 mice/group; log-rank test). **(N)** Dot plot showing the expression of *CAMK2A*, *CAMK2B*, *CAMK2D*, and *CAMK2G* in a scRNA-Seq dataset of GBM specimens (GSE182109). **(O)** Bar graph depicting mRNA expression of *CAMK2D* relative to *ACTB* as measured by qRT-PCR in GBML109 and GBML198 cells transduced with shScramble or shCAMK2D_1 (paired *t*-test). **(P)** Bar graph showing tumor sphere formation in GBML109 cultures transduced with shScramble or shCAMK2D_1 (n = 3/group, unpaired t-test). **(Q)** Immunoblot for CAMK4, CAMK2D and ACTB in G12 cells transduced with shScramble or two non-overlapping shRNA targeting P2RY1. These cells were transduced with either lenti-ctrl, lenti-CAMK4, or lenti-CAMK2D lentiviruses. * $p < 0.05$, ** $p < 0.01$, **** $p < 0.0001$, ns = not significant.

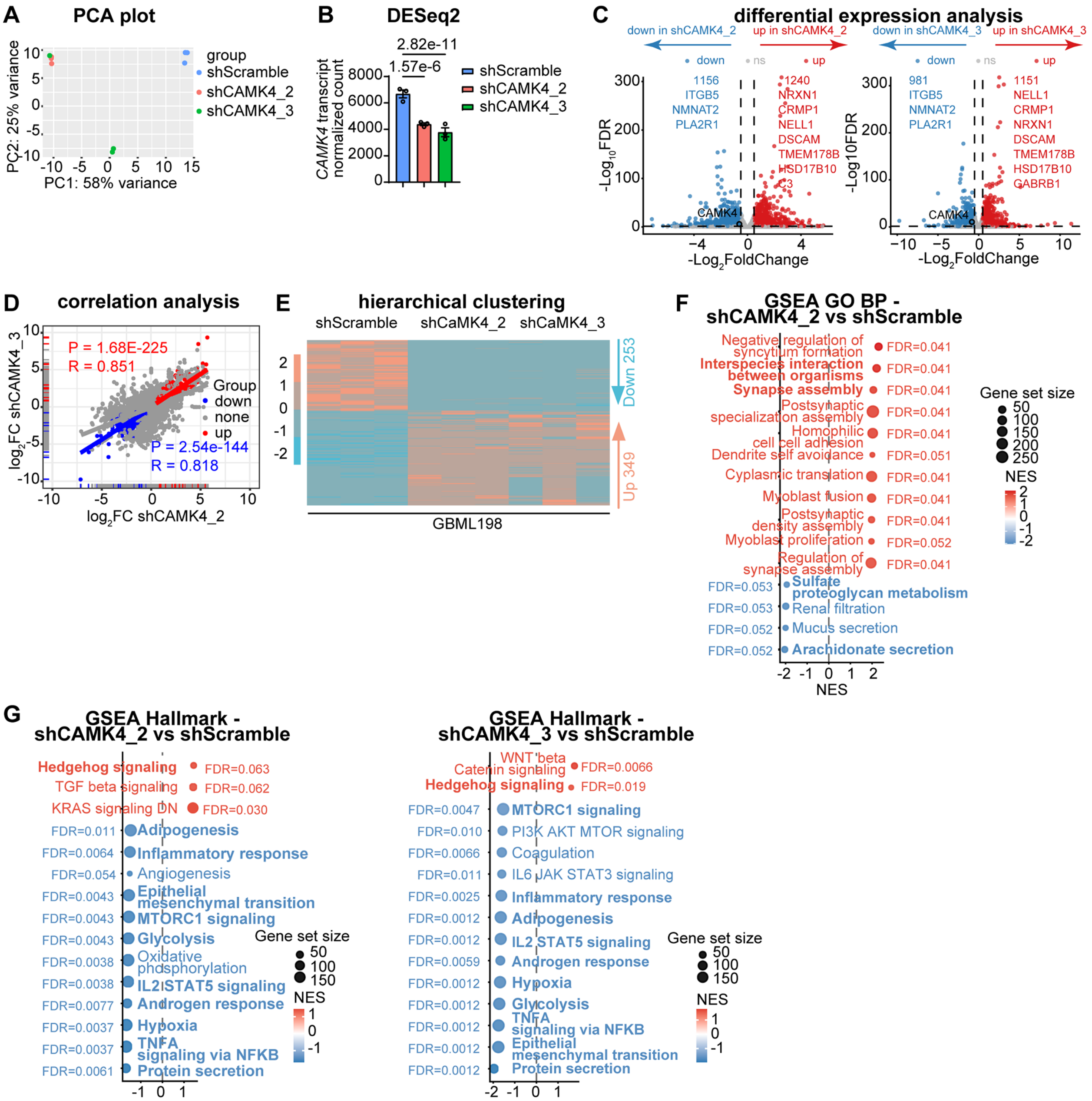


**Supplementary Figure 6: Knockdown of CAMK4 remodels the transcriptome. (A)** Principal component analysis (PCA) of raw counts from RNA-seq data. **(B)** Bar graph showing normalized count value of *CAMK4* in GBML198 transduced with shScramble or one of two non-overlapping shRNA targeting *CAMK4*. **(C)** Volcano plots showing differentially expressed genes in GBML198 cells transduced with shCAMK4_2 (left panel) and shCAMK4_3 (right panel) compared to shScramble. The top 10 upregulated and downregulated genes ranked by *P* value are labeled (|lLog$_2$FC| > 0.5). **(D)** Scatter plot depicting the correlation of Log$_2$FC in shCAMK4_2 and shCAMK4_3. The correlation coefficient and *P* value in upregulated and downregulated groups were determined by Pearson analysis. **(E)** Hierarchical clustering of differentially expressed genes in shCAMK4_2 and shCAMK4_3 compared to shScramble. **(F,G)** Bubble plots showing top ranked significant GO_BP **(F)** and Hallmark **(G)** terms by NES of GSEA results for two non-overlapping shRNA CAMK4 knockdown groups compared to shScramble.

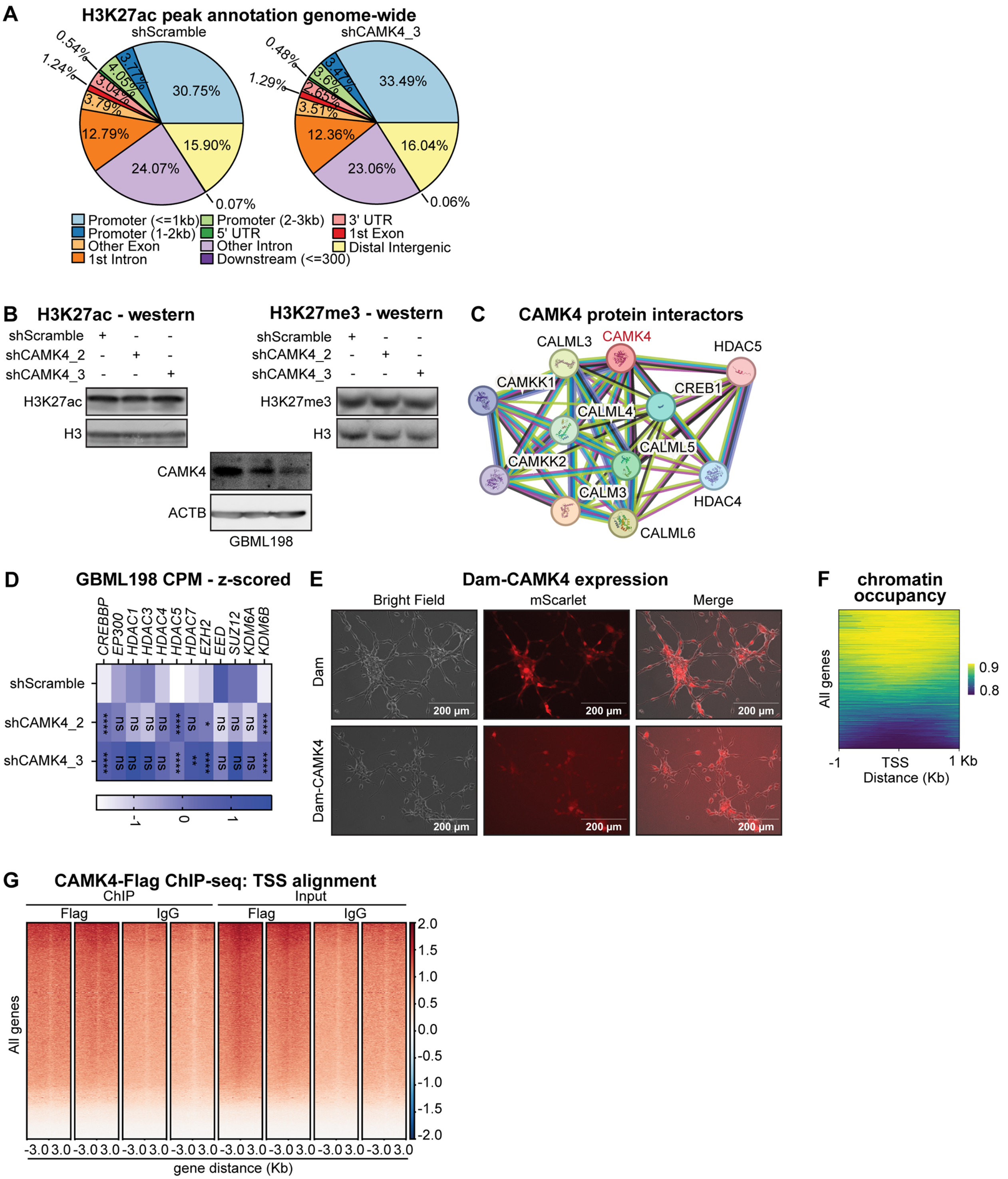


**Supplementary Figure 7: Knockdown of CAMK4 increases H3K27ac deposition at promoter regions. (A)** Pie charts showing annotations of high-confidence H3K27ac peaks across the entire genome in the shScramble and shCAMK4 groups. **(B)** Immunoblot for H3K27ac, H3K27me3, H3, CAMK4 and ACTB from GBML198 cells transduced with shScramble or one of two non-overlapping shRNAs targeting CAMK4. **(C)** STRING network analysis for protein–protein interactions (PPI) of CAMK4. **(D)** Heatmap showing z-scored CPM values of gene transcripts related to regulation of H3K27ac and H3K27me3 modifications as shown by RNA-Seq experiments. **(E)** Live cell imaging showing GBML198 cells transduced with Dam and Dam-CaMK4. **(F)** Heatmaps showing normalized Dam-CAMK4 signals aligned with Transcription Start Site (TSS) ± 1 kb. **(G)** Heatmaps illustrate CAMK4-Flag ChIP-seq signal aligned with TSS ± 3 kb, as compared to IgG ChIP and input control. **** $p < 0.0001$.

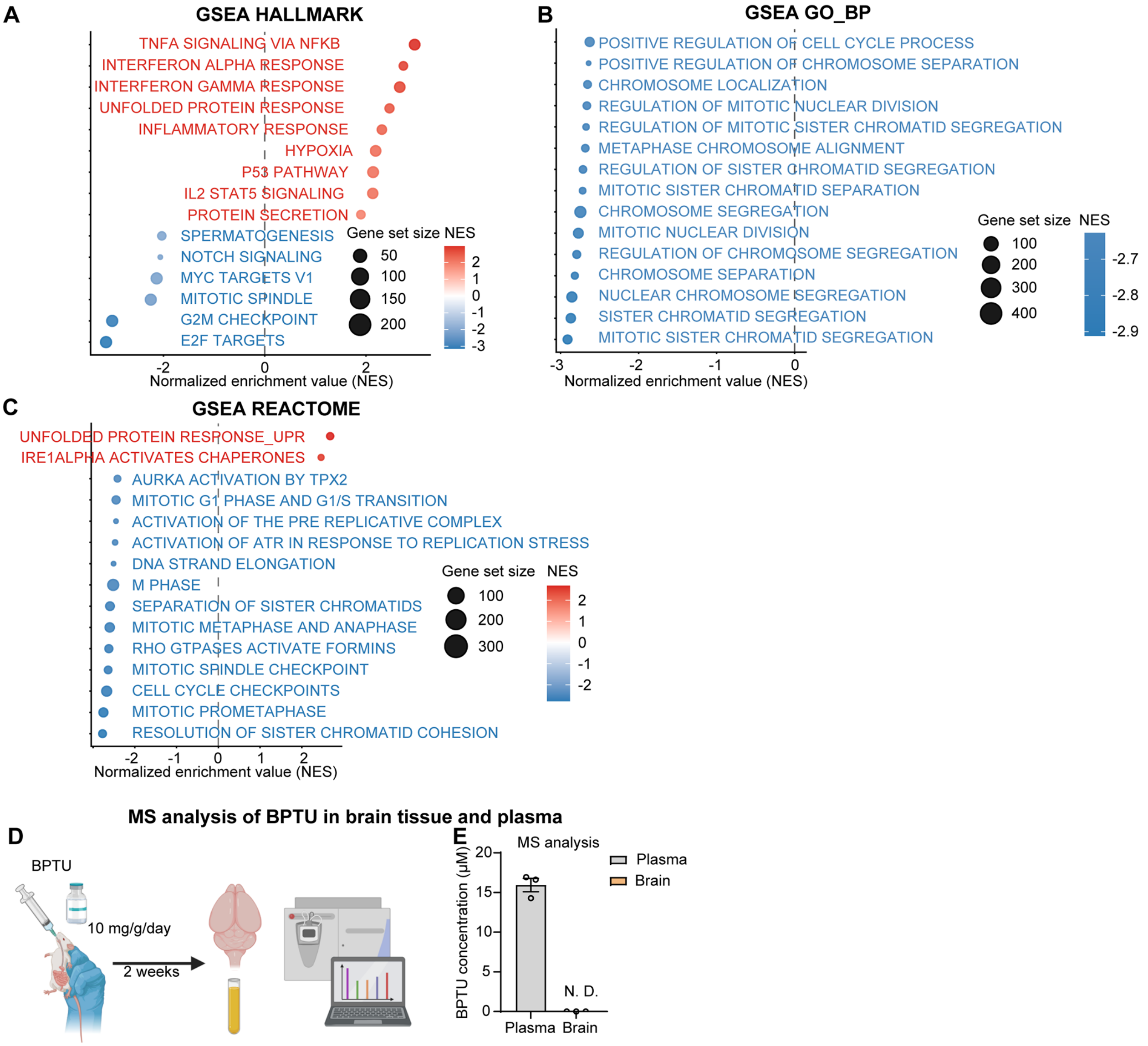


**Supplementary Figure 8: Transcriptomic effects and pharmacokinetic properties of BPTU. (A-C)** Visualization of top genesets enriched or depleted in Hallmark, GO, and Reactome GSEA analysis of RNA-seq data for GBML198 cells treated with BPTU or vehicle control. **(D)** Schematic representation of *in vivo* experiments involving BPTU administration by oral gavage (10 mg/g/day for 2 weeks), followed by mass spectrometry (MS) quantification of BPTU levels in the plasma and brain. **(E)** Bar plot showing MS measurements of BPTU levels in mouse plasma and brain (n = 3/group). N.D. = not detected.
